# Beyond Exons: Linking Noncoding Heritability and Polygenicity across Complex Human Traits and Disorders

**DOI:** 10.64898/2026.04.01.715766

**Authors:** Julian Fuhrer, Alexey A. Shadrin, Timothy Hughes, Nadine Parker, Guy Hindley, Evgeniia Frei, Dat Nguyen, Olav B. Smeland, Srdjan Djurovic, Ole A. Andreassen, Anders M. Dale, Oleksandr Frei

## Abstract

The genetic architecture of complex traits spans a continuum of polygenicity, yet it remains unclear how differences in polygenicity relate to the functional localization of SNP heritability across the genome. We use a MiXeR-based framework to partition heritability across exonic, intronic, and intergenic regions for 34 traits and introduce a likelihood-based annotation contribution score that quantifies annotation-specific impact on heritability. Exons explain a minority of heritability, and their contribution decreases with increasing polygenicity, from an average of 22% in less polygenic somatic diseases and biomarkers to 13% in highly polygenic psychiatric and cognitive phenotypes. Intergenic fractions show the opposite trend, whereas intronic fractions remain relatively stable. Analysis of a broader set of functional annotations reveals systematic differences along the polygenicity axis: highly polygenic traits show stronger contributions from comparative genomics and variant-effect scores, whereas less polygenic traits show stronger contributions in promoter, transcription, and chromatin annotations. Together, these results indicate that the functional partitioning of heritability systematically varies with polygenicity, pointing to a shift from gene-proximal regulatory architectures to architectures shaped by numerous dispersed regulatory effects as a key determinant of differences in polygenicity across traits.

## Introduction

The genetic architecture of complex traits spans a continuum of polygenicity^1^, with traits differing widely in the number of small-effect variants contributing to their heritability^2–9^. These variants occur within a genome that exhibits complex functional features, including gene structure, chromatin states, epigenetic modifications, and signatures of evolutionary constraints, which together regulate gene expression and mediate how genetic variation influences phenotypes^10–13^. Although heritability partitioning across functional annotations has been widely studied^10,14–17,6^, how these distributions differ across traits and how they relate to trait polygenicity remain poorly understood.

Complex phenotypes differ in their genetic and biological organization and regulatory architecture^10,18,19^. Mental and cognitive traits often involve dispersed, distal regulation across many brain regions and developmental windows^18,20–25^, whereas somatic traits have their genetic effects more strongly concentrated in a few specific peripheral tissues (for example, liver, adipose, and muscle), while still retaining multi-organ regulatory components^18,26–28^. Intergenic regions (IGRs) contain diverse regulatory elements and long-range contacts that modulate gene expression, yet much of their context-specific activity and target mapping remains unresolved^11,13^. These distal, noncoding mechanisms are cell-type specific and mediated through dynamic chromatin and epigenetic processes that combine and reuse a limited set of genes in diverse cellular contexts^10,11,13,23,26,27,29–31^. This diversity of regulatory architecture is consistent with models in which natural selection on complex traits shifts heritability toward numerous small, often noncoding effects^5,9,19,32,33^ and may be especially relevant for cognitive traits and mental disorders, placing them toward the distal, brain-focused end of this regulatory spectrum, characterized by particularly complex, highly polygenic architectures^20,21,23–25^. Consistent with this view, recent whole-genome sequencing (WGS) studies indicate that coding variants account for only a minority of heritability, with most attributable to noncoding variants, particularly for behavioral and cognitive phenotypes^34^. Together, these observations suggest that complex traits differ in how their heritability is distributed across coding and noncoding regulatory mechanisms. However, we still lack a systematic, multi-trait analysis that quantifies how heritability is localized across functional annotations (including gene-structure and regulatory features) and relates these patterns to differences in polygenicity.

Early work demonstrated that single nucleotide polymorphisms (SNP) contributions differ systematically across genomic categories: variants in genic elements (5’UTRs, exons, and 3’UTRs) show strong enrichment across phenotypes, followed by introns, whereas intergenic regions are typically depleted on a SNP basis^14^. These observations led to the development of statistical methods to partition heritability across functional annotations using summary statistics from genome-wide association studies (GWAS), including regression-based approaches (e.g., stratified linkage disequilibrium (LD) score regression (sLDSC)^10,35^, sLD4M^5^, and SumHer^36^), likelihood-based models (e.g., MiXeR^2^ and its annotation-informed AI-MiXeR extension^37^, and HDL^38^), and Bayesian regression models that incorporate functional annotations as priors on SNP effects (e.g., SBayesRC^39^ and related frameworks). Empirically, sLDSC analyses have shown that heritability is enriched in coding and regulatory elements and in genes that are specifically expressed in relevant tissues and cell types^18,22,40,41^. AI-MiXeR, using a two-category model that contrasts exonic and non-exonic regions, further demonstrated phenotype-specific differences in how polygenicity is distributed across these categories^37^. However, most existing work has focused on fold enrichment or annotation-specific heritability estimates for individual traits and annotations^6,10,14–17^. These metrics can disproportionately emphasize very small annotations (covering only a small fraction of SNPs) and do not directly quantify the relative contribution of each annotation to the overall genetic architecture. A likelihood-based framework can offer one way to mitigate this imbalance by providing a measure of annotation-specific contributions.

To address this, we build on the MiXeR framework^17,37^ to investigate whether trait-specific localization of heritability across genomic annotations aligns with differences in polygenicity. In contrast to approaches such as sLDSC^10,35^, which often treat functional annotations primarily as covariates to control for confounding (e.g., when estimating cell-type-specific enrichments), we treat annotations as parameters of primary interest and quantify their trait-specific contributions. Specifically, we (i) estimate how SNP heritability is partitioned across broad genomic categories (exons, introns, IGRs), and (ii) introduce a likelihood-based *annotation contribution score* (ACS), to quantify annotation-specific contributions within a broad set of functional annotations^12^, including comparative genomics (evolutionary conservation scores), variant-effect scores (computational predictions of deleteriousness or impact), and regulatory chromatin features (promoters, enhancers, and histone-marker-based states). This framework enables direct comparison of heritability profiles across traits and allows us to assess whether differences in the functional localization of heritability align with differences in polygenicity, potentially defining an axis from gene-proximal, less polygenic architectures to highly polygenic architectures dominated by dispersed distal regulation.

## Results

We extended the MiXeR framework to model a large set of functional annotations and applied it to 34 complex traits and disorders. Across all traits, most heritability was concentrated outside coding sequence, and its partitioning varied systematically across exonic, intronic, and intergenic regions with trait polygenicity. We then used ACS to evaluate this broader annotation panel across traits and relate its contributions to differences in polygenicity.

Briefly, we modeled phenotypic variance as a combination of additive genetic effects and residual variance, with SNP effect-size variance depending on functional annotations (Table S1). Model parameters were inferred from GWAS z-scores using maximum likelihood estimation, accounting for LD. We selected large, well-powered GWAS spanning psychiatric, neurological, cardiometabolic, anthropometric, hematological, immune, and other organ-system traits (Table S2), and then pruned from 70 phenotypes to 34 by clumping at genetic correlation r_g_=0.3, prioritizing higher-heritability traits (Figs. S3–S4). ACS quantifies the relative contribution of each functional annotation, accounting for its size (*Methods*).

### Phenotype’s heritability linked to polygenicity across genomic regions

Across the genome, exons make up 2.55% of base pairs, whereas introns and IGRs together cover 97.45% (introns 45%, IGRs 52.45%). We partitioned heritability across exons, introns, and IGRs for 34 traits and found that exonic regions account for a minority of heritability (mean 14.52±4.72%), with introns and IGRs together explaining the remaining 85.49±4.72% (Table S2; Fig. S4).

When we related these regional fractions to polygenicity, a clear pattern emerged (Fig. 1): With increasing intergenic heritability fraction, the exonic fraction decreases, and the intronic fraction remains approximately constant, while the polygenicity increases. In a pooled regression of regional heritability fractions on −log_10_(π), exonic fractions decreased and intergenic fractions increased significantly, whereas intronic fractions did not significantly vary (exon slope=-4.38, intergenic slope=4.87, intron slope=-0.49). Spearman correlations showed the same pattern (exons r=-0.51, introns r=-0.18, intergenic r=0.59). Across traits, introns alone typically account for roughly half of the heritability, exceeding the exonic and intergenic fractions for most phenotypes. Thus, exonic and intergenic fractions trade off with polygenicity, while intronic heritability remains relatively stable (Table S4).

**Figure 1:**
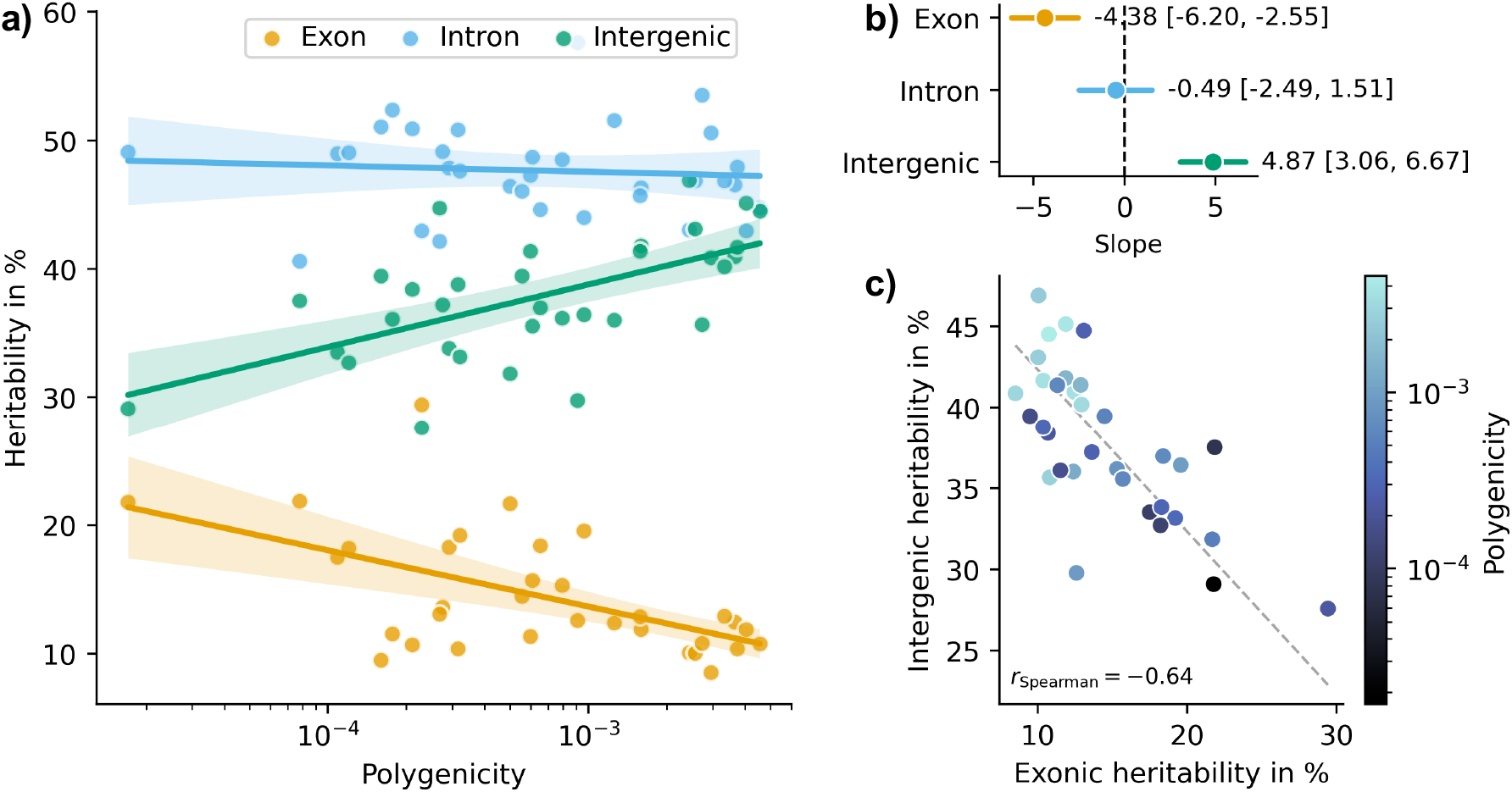
Regional heritability partitions versus polygenicity. **a)** Fractions of heritability across exons, introns and IGRs, with phenotypes sorted by increasing polygenicity. Exonic fractions decrease with polygenicity, intergenic fractions increase, and intronic fractions remain approximately constant. Slopes and confidence intervals are estimated from pooled OLS fits with a Region×log10(π) interaction and trait-clustered standard errors. **b)** Region-specific slopes from the same model (defined as the percentage-point change in regional heritability per 10-fold increase in polygenicity). **c)** Exonic versus intergenic heritability fractions for each trait, colored by polygenicity. More polygenic traits cluster toward lower exonic and higher intergenic fractions. The dashed line indicates the exon-intergenic relationship expected if intronic heritability were fixed at its mean across traits.

For illustration, anorexia nervosa, neuroticism, and schizophrenia had exonic fractions of 10.06%, 10.37%, and 8.51%, respectively, so that introns and IGRs together explained over 89% of heritability in each case (with genic regions, exons plus introns, still contributing more than IGRs). In contrast, height had an exonic fraction of 19.58%, cortical surface area of 21.69%, and sex hormone-binding globulin of 29.44%. Thus, exonic heritability is roughly two-to threefold higher in these somatic traits compared to the mental and cognitive traits, even though introns and IGRs still account for the majority in both groups (Fig. S1).

### Functional annotation panel and its genomic composition

We first describe the functional annotation panel used in our analyses, including its exonic, intronic, and intergenic composition and its grouping into broader functional domains. We constructed a panel of 74 annotations (Table S1) based on the baseline-LD v2.3 model^12^, including high-resolution conservation and variant-effect scores, and expanded regulatory or QTL unions. Consistent with previous analyses^42^, we excluded flanking annotations for interpretability and omitted continuous categories. We further updated gene-structure annotations (coding, introns, 5’UTRs, 3’UTRs) using curated NCBI RefSeq^43^ tracks (adding IGRs as the complement of “WholeGene”). For interpretability, we grouped annotations into seven domains: Genes (coding and gene-structure features), Comparative Genomics (evolutionary conservation scores), Variant Effect and Literature (computational impact predictors and GWAS-or database-derived scores), Enhancer (enhancer markers and unions), Promoter and Transcription (promoter and transcription start site annotations), Chromatin State (integrated chromatin segmentation states), and Epigenetics (histone-markers unions and other epigenomic signal), following UCSC Genome Browser^43^ taxonomy (see Table S1 for brief descriptions and *Methods*).

Fig. 2 shows the genomic composition (i.e., the content of exons, introns, and IGRs) of each annotation and its size (number of SNPs per annotation). Epigenetic annotations (mainly histone marker unions) and enhancer annotations were the largest tracks and were dominated by noncoding nucleotides. Promoter and transcription annotations lay closer to exons but were still predominantly non-exonic. Chromatin state annotations showed moderate overlap, with promoter- and TSS-like chromatin states partially overlapping promoter-associated histone markers and CpG islands, and broader cell-type union chromatin states showing lower overlap (Fig. S2). Comparative genomics and variant-effect and literature annotations were much smaller in size and had mixed composition across broad genomic categories (exon, intron, IGR): Comparative genomics tracks were more exon-enriched yet still largely noncoding, and variant-effect tracks ranged from exon-rich to mostly noncoding high-impact subsets (e.g., extreme tails or binarized annotations; Table S1).

**Figure 2:**
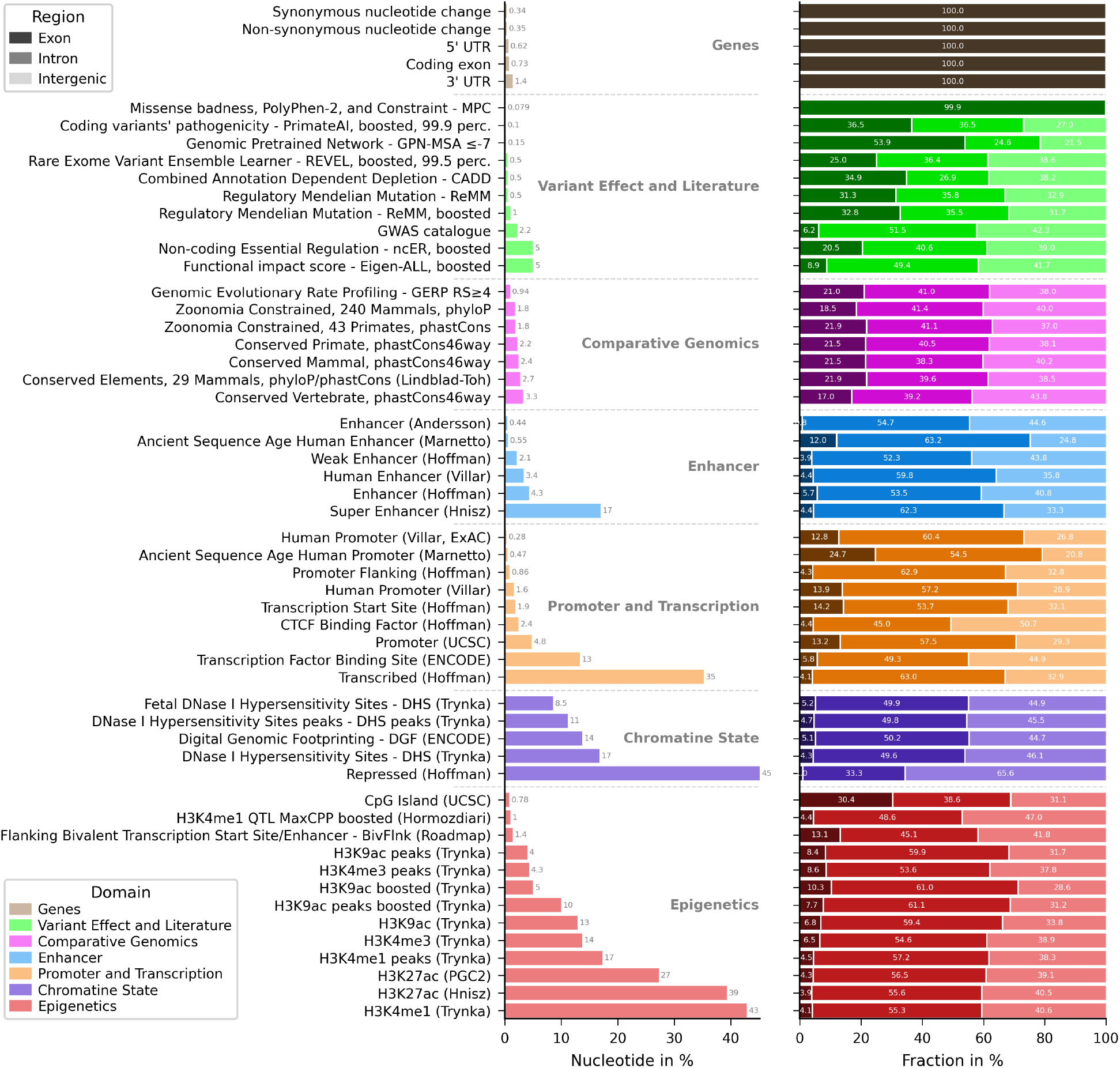
Domain-specific genomic composition of functional annotations. Left: relative nucleotide coverage of functional annotations. Right: respective fractions of broad genomic regions (exons, introns, and IGRs). Functional annotations are grouped by domains and sorted by their respective mean coverage. The nucleotide proportions by genomic regions were exons 2.55%, introns 45.00% and IGRs 52.45%.

### Relative contributions of functional annotations across traits and polygenicity

We next quantify the relative contributions of each functional annotation to heritability across traits using ACS, allowing us to compare how different annotation domains track variation in polygenicity. We clumped all annotations using pairwise Dice coefficients and thresholding at 0.8 (Fig. S2), retaining larger annotations. Treating exons, introns, and IGRs as annotations alongside other functional annotations, we then asked how their ACS profiles vary with polygenicity. When we ordered phenotypes by polygenicity and clustered annotations by similarity of their ACS profiles (Figs. 3, S5), a clear pattern emerged: higher-polygenicity traits loaded more strongly on comparative genomics and variant-effect annotations (e.g., phastCons, GERP, CADD, Eigen, ReMM), indicating that compact, constraint- and impact-based annotations capture a larger fraction of their heritability. On the other hand, less polygenic traits showed stronger contributions from promoter, transcription, and promoter-like chromatin annotations (e.g., Villar promoters, Hoffman TSS, H3K4me3, H3K9ac, CpG islands), suggesting that their genetic architecture is more concentrated in gene-proximal regulatory regions. Several traits (e.g., height, systolic blood pressure) showed mixed profiles with simultaneous loadings across enhancer, promoter, and transcription, and comparative genomics or variant-effect annotations.

**Figure 3:**
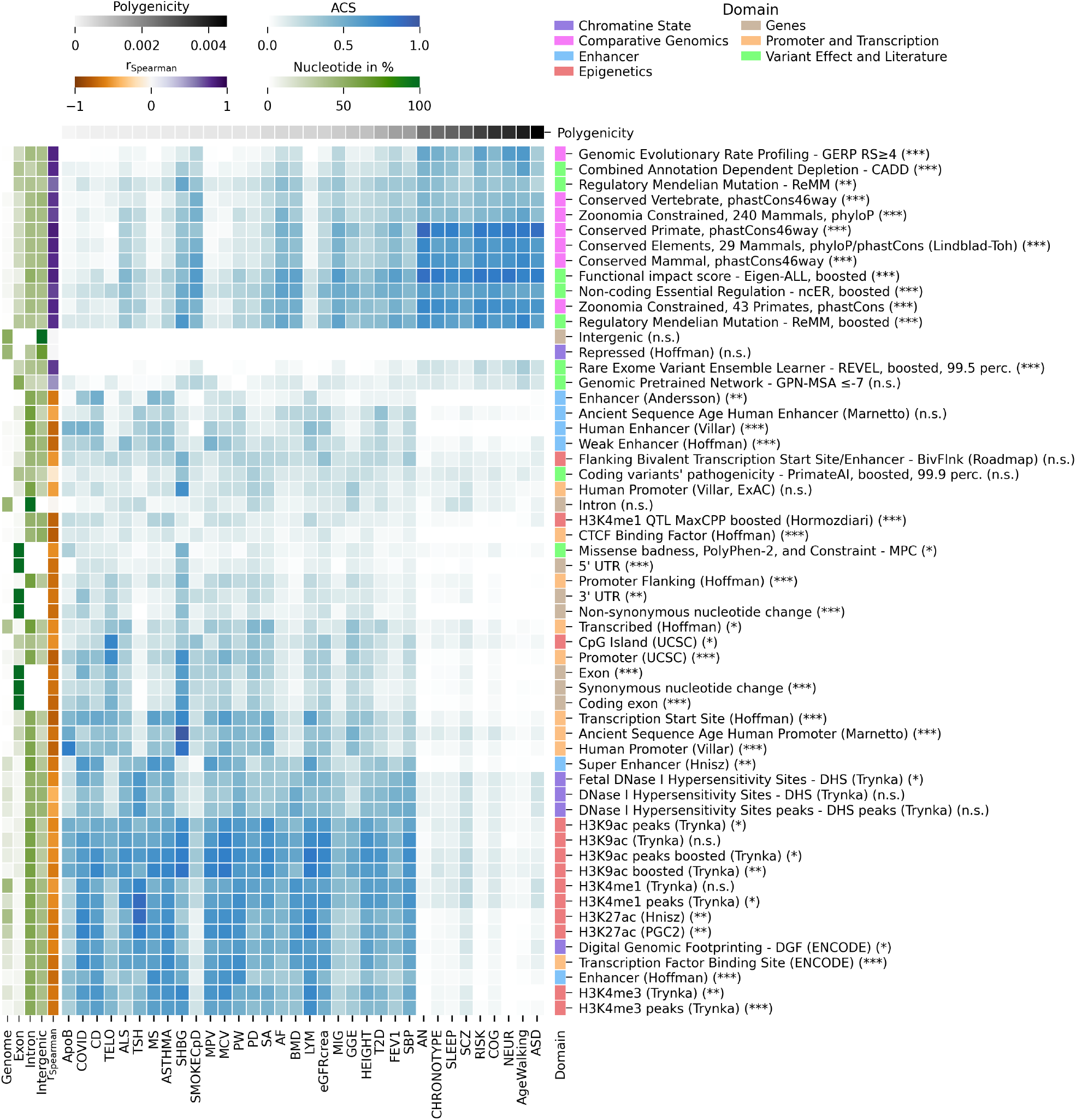
Contribution of individual functional annotations relative to the full model with 74 annotations. Phenotypes are ordered by their polygenicity remaining after pruning with *r*_*g*_=0.3, favoring phenotypes with higher heritability. Annotations are sorted by their Euclidean distance. On the left, next to relative numbers of nucleotides (genome, exon, intron, intergenic), the Spearman rank correlation value between ACS and polygenicity across phenotypes is indicated. The symbols following each functional annotation on the right indicate the significance after Bonferroni correction for the Spearman correlation (significance: ***: p<0.001, **: p<0.01, *: p <0.05, n.s.: p≥0.05).

Across annotations, ACS showed three broad patterns of association with polygenicity (positive, weak/near-zero, and negative; Table S3). Comparative genomics and variant-effect and literature domains showed positive correlations with polygenicity, whereas promoter and transcription, enhancer, chromatin-state, gene, and epigenetic domains were negative on average (Figs. 2, S5). The strongest associations were seen for conserved primate sequence (phastCons46way; positively correlated) and CTCF binding sites (|r| ≥ 0.79, p ≤ 6e-8). Comparing correlations across domains (permutation with Benjamini-Hochberg FDR correction; Tables S4–S5) confirmed that comparative genomics and variant-effect domains track polygenicity more strongly than promoter and transcription, enhancer, and chromatin domains.

As a supporting analysis, we added cell-type union annotations from Finucane et al.^10^ jointly to the full model and evaluated ACS-based contributions (Fig. S16). We observed central nervous system contributions for schizophrenia and cognition, consistent with neuronal or glial involvement, elevated hematopoietic and gastrointestinal enrichments for Crohn’s disease, whereas pancreatic or hepatic signals for type 2 diabetes and lipids were more modest in this broad cell-union framework. These results indicate that our annotation-informed MiXeR framework reproduces established tissue-specific enrichment patterns.

When we examined ACS as a function of annotation size (Figs. 2, 3), ACS values peaked for mid-sized annotations covering roughly 1–15% of SNPs, with slightly smaller or larger categories close behind, whereas very small or large categories contributed less overall (Table S6). Some annotations not biologically connected to exons showed mild exon enrichment (e.g., transcription factor binding site (ENCODE) and chromatin-state annotations). This likely reflects promoter and transcription windows extending into 5’UTR and first exon, fuzzy boundaries from unions of peaks and windowed definitions, and gene-dense regions where regulatory sequence of one gene overlaps the coding sequence of a neighbor. These features are important to keep in mind when interpreting exon-enriched annotations in the functional-annotation analyses.

Crucially, the behavior of exonic annotations helps reconcile the regional and annotation-level results. Consistent with Fig. 1, exonic annotations show relatively greater ACS in low-polygenicity phenotypes (Fig. 3), but their limited genomic coverage (Fig. 2) constrains their absolute contribution, so they occupy only mid-range positions in terms of overall heritability explained. Thus, even when exonic categories are highly enriched on a SNP basis in less polygenic traits, diffuse noncoding annotations still dominate total heritability and become increasingly important as traits become more polygenic.

Despite their relatively small genomic coverage, comparative genomics and variant-effect and literature annotations account for a disproportionate fraction of ACS in highly polygenic phenotypes, whereas large epigenetic and chromatin-state annotations contribute more prominently to less polygenic traits (Fig. 3). More generally, compact, information-rich annotations can show strong per-SNP enrichment yet contribute less total heritability than broad regulatory categories that cover larger genomic regions. Interpreting enrichment, therefore, requires considering both effect-size concentration and genomic coverage, especially when comparing traits that differ in polygenicity.

### Sensitivity analyses

We performed several sensitivity analyses to examine model assumptions and the robustness of our findings (*Supplementary Results, Methods*). First, we compared heritability fractions estimated from sLDSC’s infinitesimal model versus sLD4M’s regression model^5^, which allows both polygenicity and per-SNP effect size variance to vary across individual functional categories. Estimates were nearly identical across phenotypes and annotations (Pearson r=0.996; Figs. S8a, S9). Second, polygenicity estimates from univariate MiXeR^2,3^ and sLD4M were highly correlated (Spearman r=0.9; Fig. S8b). Third, ACS estimates from infinitesimal and non-infinitesimal annotation-informed MiXeR fits were strongly concordant (Pearson r=0.92; Figs. S10–S13), with differences confined mainly to lower-polygenicity traits and no change in domain-level rankings. Finally, clustering traits by cosine similarity of annotation-specific heritability profiles showed that traits with similar polygenicity have more similar heritability landscapes than expected by chance (Figs. S14–S15). Including a GWAS Catalog annotation track improved model fit and yielded the largest relative improvement in log-likelihood across all traits, however, due to potential circularity, it is reported only as a sensitivity analysis (Fig. S7).

## Discussion

Our results show that the distribution of heritability across functional genomic partitions is closely tied to the polygenicity of a trait. Two robust patterns emerged. First, the partitioning of heritability across exons, introns, and IGRs shifts systematically with polygenicity: as traits become more polygenic, the intergenic heritability fraction increases while the exonic fraction decreases. Exons, although enriched relative to their small genomic span, explain a minority of heritability, especially in cognitive and mental traits, whereas intronic sequence consistently accounts for roughly half of the heritability across all traits. Second, functional annotation domains can be stratified by polygenicity: comparative genomics and variant-effect scores dominate higher-polygenicity traits, whereas regulatory chromatin tracks contribute more, and show more phenotype-specific contribution patterns in less polygenic traits.

Taken together, these patterns indicate that complex traits do not share a single genetic architecture but instead occupy distinct positions along a spectrum from gene-proximal to highly dispersed, distal regulation. Biochemical and hematological traits such as lipids, glycemic indices, and many blood cell measures reflect relatively specific physiological pathways and often depend more on a limited set of coding and proximal regulatory variants, consistent with strong constraints on gene function^18,19,44–48^. In contrast, cognition and psychiatric disorders are higher-order phenotypes that integrate information across many brain regions, cell types, and developmental trajectories. For these traits, genetic influences are expected to act largely through dispersed noncoding regulation of shared gene sets^15,20– 25,32,33,40,41^

In our results, this phenotypic variation is mirrored in the major genomic partitions. Traits with more dispersed, context-dependent biology show higher fractions of intergenic heritability and lower exonic fractions (Figs. 1, S1), whereas the intronic fraction remains relatively stable, and typically exceeds the exonic and intergenic fractions, consistent with a gene-proximal baseline of regulatory and splice-related elements (for example, first-intron enhancers, promoter-proximal regions, splice sites)^5,19,49^. Because such intronic regulatory and splice features are present around most expressed genes and are reused across traits, their aggregate contribution scales relatively uniformly with overall heritability, leading to a stable intronic fraction even as exonic and intergenic contributions trade off with polygenicity. This intronic dominance suggests that regulatory and splice-related variation within gene bodies contributes more heritability than is often appreciated in simple “coding versus intergenic regulation” models. Increasing intergenic fractions of heritability are associated with higher polygenicity beyond a simple size effect because distal annotations carry higher ACS values (Fig. 3)^10,6,19,16^. As genetic effects become more numerous and more dispersed across cell types and developmental contexts, they are increasingly mediated by distal elements in IGRs (enhancers, silencers, insulators, long-range contacts), naturally elevating the intergenic fraction of heritability while exonic contributions decline.

Several evolutionary mechanisms plausibly contribute to these phenotype-related differences in genetic architecture. Negative selection purges strongly deleterious variants, induces MAF- and LD-dependent architectures, and shifts heritability toward numerous small effects, many of them noncoding^5,9,19,44–48,50^. Under stabilizing selection, fitness effects are spread across many loci and traits, favoring architectures dominated by numerous small regulatory effects^32,33^ and leading to broad pleiotropy across traits^15^. In parallel, trait biology exhibits clear cell-type specificity, for example, CNS enrichment for psychiatric phenotypes, immune for autoimmune and inflammatory, and adipose or hepatic for cardiometabolic traits^10,23–28^, supported by our supplementary cell-type union analyses (Fig. S16). Within this context, our results support a qualitative distinction between evolutionarily “basic” and “higher-order” traits: basic physical and physiological traits rely more heavily on a constrained set of coding and proximal regulatory variants, whereas evolutionarily complex, human-specific traits such as cognition and psychiatric disorders show architectures dominated by dispersed noncoding regulation of largely shared gene repertoires. Consistent with this interpretation, evolutionary modeling of GWAS hits suggests that CNS-mediated traits have larger mutational target sizes and are under stronger negative selection than other complex traits^51^.

At the domain level defined by the UCSC-Genome-Browser, ACS values concentrate in “umbrella” annotations of comparative genomics and variant-effect and literature scores as polygenicity increases, whereas regulatory chromatin domains show stronger contributions in lower-polygenicity traits. Comparative genomics and variant-effect domains preferentially tag evolutionarily constrained sequence, including both conserved noncoding regions and sites where variation is predicted to strongly affect splicing or protein sequence^43,52^. At first glance, this may seem at odds with the exon-IGR heritability pattern, because exons are among the most highly conserved regions and contain many high variant-effect scores, whereas regulatory chromatin annotations are largely intergenic. However, conservation and impact scores also capture extensive conserved noncoding sequence and deleterious regulatory variants outside exons, and are defined on relatively compact, information-rich subsets of the genome. In contrast, chromatin annotations tend to be broad, often spanning large bands of intergenic and intronic sequence with heterogeneous function. When effects are diffuse across cell types and contexts, these broad conservation- and impact-based annotations appear to generalize better than any single cell-specific regulatory track. They efficiently capture intergenic signal, in part because distal enhancers and other intergenic regulatory elements are typically conserved across species and therefore constitute a major subset of conserved noncoding sequences^43,52^. In contrast, lower-polygenicity, more gene-proximal traits (lipids, glycemic indices, hematological measures) show balanced loadings across promoter, transcription, and chromatin domains, with regional heritability concentrated near genes^18,52,53^. Traits such as height, systolic blood pressure, atrial fibrillation, estimated glomerular filtration rate, and bone mineral density occupy intermediate positions with mixed contributions, probably reflecting blended distal and proximal mechanisms.

Methodologically, our ACS metric quantifies trait-specific functional annotation contributions while balancing annotation size against fold enrichment (average heritability within an annotation relative to the genome-wide average). This reduces the tendency of standard fold enrichment to overemphasize very small, high-specificity categories that contribute little to total heritability. Across traits, mid-sized annotations (covering roughly 10–15% of SNPs) yield the highest ACS values, suggesting that such compact, information-rich annotations may be particularly useful as priors for fine-mapping and prediction^16,39,54^. An aggregated GWAS Catalog track improved model fit and was the strongest single annotation contributor (Fig. S7), consistent with the predominance of noncoding GWAS associations. We treated this as a sensitivity analysis because of potential circularity with our GWAS inputs. Nevertheless, it has potential as a general prior when used out-of-sample^12,43^.

Our findings also have implications for study design. Genome-wide arrays with imputation are cost-effective for common SNPs and benefit from annotation-informed models, but they indirectly capture noncoding architecture. Whole-exome sequencing (WES) is efficient when effects concentrate in coding regions, whereas WGS is most informative for regulatory or noncoding signal but at higher cost^55^. Long-read WGS (e.g., nanopore) further improves mapping, phasing and structural variant detection in IGRs but at higher cost and lower throughput^56^. Given that coding SNPs explain only a minority of heritability and that noncoding contributions increase with trait polygenicity^34^, especially for behavioral, mental and cognitive phenotypes, these patterns argue for prioritizing sequencing designs that maximize noncoding coverage (for example, WGS over WES when feasible) and for using functionally informed priors to extract more information from array-based GWAS in future largescale studies.

Limitations include reliance on European-ancestry reference panels and a focus on relatively broad functional annotations with incomplete coverage of chromosome X. A next step is to apply this framework at finer resolution to dissect noncoding contributions within specific gene sets, pathways, and regulatory networks, and to replicate analyses in diverse ancestries. Our conclusions also depend on polygenicity estimates from MiXeR and sLD4M, which showed strong concordance across methods in sensitivity analyses.

Together, these results provide a novel organizing framework for complex trait genetic architecture: less-polygenic traits tend to have heritability concentrated in gene-proximal coding and regulatory regions, with introns accounting for a large, relatively stable fraction, whereas more polygenic traits tend to have heritability distributed across dispersed distal regulatory elements. In evolutionary terms, this is consistent with a model in which basic somatic functions are encoded by relatively few, highly constrained genes and their proximal regulatory elements, whereas higher-order mental and cognitive traits are tuned by many small, largely noncoding effects distributed across regulatory networks. This explains why noncoding regions can dominate heritability even when individual categories show modest fold enrichment, and why broad evolution-or impact-based annotations often explain more than single enhancer or promoter tracks for highly polygenic traits. Conceptually, this organizing framework unifies diverse enrichment and contribution signals with polygenic architecture. Practically, it informs about when broad functional priors aid fine-mapping and prediction, and when specific annotations in coding and nearby regulatory regions are most informative. More broadly, clarifying how polygenicity maps onto the functional genome may help to better understand the molecular mechanisms of complex traits and diseases and ultimately shape strategies for therapeutic and precision-medicine applications.

## Methods

### MiXeR model, heritability, and annotation contribution score

We work within the unified MiXeR framework with spike-and-slab priors on SNP effects (see *Supplementary Note*). Throughout, the index 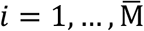 runs over all modeled variants (typically SNPs in a reference panel), while the index *j* = 1, …, M runs over GWAS tag SNPs with available z_j_-scores. By construction, the set of tag SNPs is a subset of the modeled variants on which the additive effects βi are defined on. In the main analyses, we use the infinitesimal case of this framework (π_i_ = 1 for all modeled variants), so that 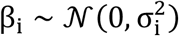 and each z_j_ is marginally Gaussian. The effect-size variance 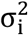 is allowed to depend on a set of functional annotations: Let A_p_ (*p* = 1, …, N_a_) denote the functional categories and introduce a core category A_0_ including all variants, then the additive functional-annotation model can be defined as 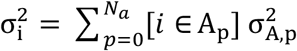, where [*i* ∈ A_p_] is 1 if SNP *i* belongs to category A_p_ and 0 otherwise. For the analyses reported in the main text, the MAF-dependence parameter is fixed (*S* = 0; *Supplementary Note*) when fitting annotation models and regional heritability fractions. Given GWAS z_j_, effective sample sizes N_j_, LD correlations r_ij_, and heterozygosities H_i_, we precompute heterozygosity-adjusted LD scores for each tag SNP *j* and annotation *p* which reduces to 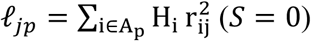. Under the infinitesimal constraint, the functional-annotation log-likelihood expression can be then formulated as

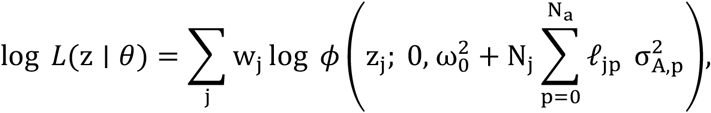

where ϕ(⋅; 0, s^2^) is the 𝒩 (0, s^2^) density, w_j_ are LD-based weights, 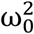 is a residual variance term capturing non-polygenic inflation in z-scores, and the parameter vector is 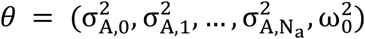. All MiXeR models in the main text are fitted by direct maximization of this univariate log-likelihood in log-variance space 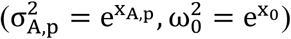, using analytical gradients.

### Core, partial, and full models

Using the log-likelihood log*L*(z | *θ*) formulation, we fit three infinitesimal models (core, partial, full) for each trait, differing how 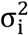 is parameterized:

1. **Core model**: all SNPs share a single variance parameter 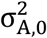 (no annotation effects), i.e., 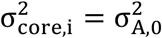, defining log *L*_core_.
2. **Partial model**: SNP effect-size variance is modeled as a shared “core” component plus an additional component for SNPs in annotation *C*: 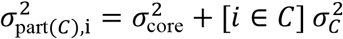 with the variance 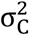 attributable to membership in *C*, defining log *L*_part(c)_.
3. **Full model**: Uses the entire set of functional annotations, that is to say, 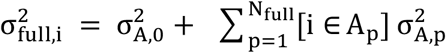, defining log *L*_full_.

### Heritability fractions in exons, introns, and IGRs

We partition the heritability across RefSeq exonic, intronic, and intergenic regions using the fitted full model. Let R_1_, R_2_, R_3_ denote the three RefSeq categories (exon, intron, intergenic). Using the variant-based heritability identity (*Supplementary Note*), the fitted full model implies SNP-specific heritability contributions 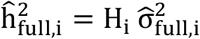 and the full and regional contributions for the three RefSeq categories respectively are

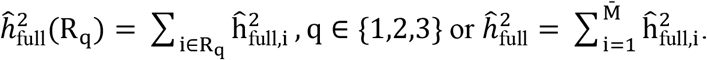

The reported full-model heritability fractions are

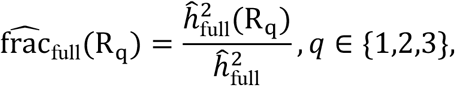

so that the full model is fitted on the complete annotation set, but the final readout is collapsed back to the three RefSeq categories (exon, intron, intergenic).

### Annotation Contribution Score

To quantify how much of the full model’s log-likelihood improvement over the core model can be captured by a single functional annotation, we introduce the annotation contribution score (ACS) for each trait and annotation *C* using log *L*_core_, log *L*_full_, and log *L*_part (c)_ as

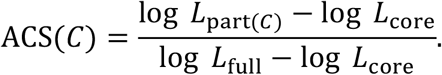

ACS measures the fraction of the full model’s total improvement in model fit (relative to the core model with no annotation effects) that is recovered by a model in which only annotation *C* has its own variance parameter. This is conceptually similar to fold enrichment in stratified sLDSC^10^, but here we use the change in log-likelihood in a joint model as the primary summary statistic. The approach is related to the GSA-MiXeR framework^17,57^, which compares full and core likelihood models to estimate gene-set fold enrichment. In our case, we apply a similar likelihood-based idea to a heterogeneous panel of functional annotations and report normalized log-likelihood improvements rather than fold-enrichment ratios. In practice, ACS integrates both annotation size and effect-size scaling through the joint likelihood, which is particularly useful when annotation sizes differ greatly, as it avoids ranking extremely small, high-specificity annotations disproportionately highly when they explain only modest fractions of total heritability. In addition to the infinitesimal fits, we performed a sensitivity analysis in which we explicitly fitted the polygenicity parameters π under the causal-mixture prior, while keeping the heritability profile fixed. This analysis was carried out separately for the core, partial, and full models, and the resulting likelihoods and ACS values closely matched the infinitesimal results, indicating that our conclusions are robust to relaxing the infinitesimal assumption.

### Functional Annotations

We utilized the set of 97 binary annotations designed for stratified LD score regression (baseline-LD v2.3)^10,12^. We restricted to binary annotations and further excluded 33 categories that extended the main annotations by 500 base pairs upstream and downstream as these categories are primarily intended as statistical controls rather than for direct inference^42^. Gene-structure annotations (exons, introns, 5’UTRs, 3’UTRS and IGRs, where IGRs are defined as the complement of the “WholeGene” track) were obtained from the curated National Center for Biotechnology Information Reference Sequence Database (NCBI RefSeq)^43^ and used instead of the corresponding gene-structure annotations in the baseline-LD v2.3 set. Nucleotides with an ambiguous exon-intron status (due to multiple transcripts) were considered exonic to ensure non-overlapping genomic regions. Given the strong signals for the continuous annotation “CpG Content 50kb” in UK Biobank phenotypes^42^, we added a binary CpG-island annotation from the UCSC Genome Browser^43^. For variant-effect prediction, we additionally derived a binary genomic pretrained network with multiple-sequence alignment (GPN-MSA) annotation^58^, based on a recent DNA language model that provides genome-wide variant effect scores. For each base pair, we averaged scores across the four possible single-nucleotide substitutions and then binarized the annotation using a threshold of -7 (the recommended deleteriousness cutoff). To identify highly similar annotations, we calculated nucleotide-based pairwise dice coefficients (Fig. S2) followed by clumping using a dice threshold of 0.8, favoring larger annotations. In addition, for sensitivity analyses we used a “GWAS Catalog” annotation track from the UCSC Genome Browser, which is based on the NHGRI-EBI GWAS Catalog^59^ of reported genome-wide association loci. We restricted this track to variants reported in European-ancestry cohorts, which was then added as an extra category on top of the baseline-LD annotations.

### 1000 Genomes Phase3 reference (LD structure)

We made use of the 1000 Genomes Phase3 (1kG) reference panel. Our 1kG reference has 9,997,232 SNPs and 489 samples after basic QC procedure. During sample QC, we select individuals of European ancestry defined from the first two principal components of 1000 Genomes Phase3 data. Further, we prune related individuals using KING software (‘king --unrelated --degree 2’). For variant QC, we use ‘plink --maf 0.001 --hwe 1e-10 --geno 0.1’ parameters and further exclude markers without reference SNP (RS) IDs and excluding duplicated RS IDs. The calculation of allelic correlation (LD r^2^) was done from hard calls, separately for each chromosome, with a window size of 10 MB, and using a minimum LD r^2^ threshold of 0.01.

### Phenotypes and GWAS summary statistics

Table S2 summarizes the phenotypes and publicly available GWAS summary statistics used in this study. We assembled a broad panel of 70 phenotypes spanning anthropometric measures, clinical outcomes, quantitative biomarkers and metabolites, psychiatric and neurodevelopmental conditions, neurological disorders, cognitive and brain-structure measures, immune and inflammatory conditions, endocrine traits, hematology traits, respiratory and musculoskeletal phenotypes, infection outcomes, substance u-use measures, sleep or chronotype traits, and functional or aging measures. Our goals in selecting this range of phenotypes were, first, to capture multiple levels of biology from molecular biomarkers to clinical outcomes, and second, to include phenotypes with different genetic and pleiotropic architectures, maximizing the potential for discovery and biological interpretation. We further selected the latest, publicly available, European-ancestry summary statistics. When several summary statistics for a phenotype were available, we prioritized the most recent study and otherwise selected the largest one. Summary statistics were harmonized to dbSNP with the cleansumstats pipeline^60^, excluding SNPs with MAF below 5% in the reference panel, strand-ambiguous SNPs, indels, SNPs without an rsID, and the extended major histocompatibility complex (MHC; chr6: 25652429–33368333 MB, GRCh37) due to long-range LD. For Alzheimer’s disease, chromosome 19 was excluded^2,21^. Summary statistics were pruned via inverse LD-weighting (*Supplementary Note*). For the log-likelihood-based analysis, we selected the phenotypes by thresholding at r_g_=0.3 and preferring phenotypes with higher heritability based on global LD score regression^35^.

## Acknowledgments

We thank the research participants, employees, and researchers for making this research possible.

## Author Contributions

Conceptualization: JF, OF, AS, GH, NP, AMD, OA; Data curation: JF, OF, AS, NP, TH; DN, Formal analysis: JF, OF, AS, NP; Funding acquisition: OAA, AMD; Software: JF, OF, AS AMD; Visualization: JF, OF, TH, GH, AS; Writing – original draft: JF, OF; Writing – review & editing: all listed authors.

## Data and Code Availability Statement

For peer review, the relevant code for all analyses can be accessed via the supplied archive named “XXX.zip”, which will be made public upon publication.

