## Supplementary material for "Beyond Exons: Linking Noncoding Heritability and Polygenicity across Complex Human Traits and Disorders": SI

**Supplementary Notes – MiXeR** 4

**Supplementary Results** 18

Robustness to Model Assumptions and Polygenicity Estimation . . . . . . . . . . . . . . . . . . 18
Minor-Allele-Frequency Bins as Model Covariates . . . . . . . . . . . . . . . . . . . . . . . . 19
Functional Annotation Properties and Overlap . . . . . . . . . . . . . . . . . . . . . . . . . . 19

#### Supplementary Figures

S1 Fractions of SNP heritability partitioned across exons, introns and IGRs with phenotypes sorted by polygenicity. The solid line shows polygenicity estimates. With increasing polygenicity, heritability decreases in exons, increases in IGRs, and stays invariant in introns. . . . . . . . . . . . . . . . . . . . . . . . . . . . . . . . . . . . . . . . . . . 21
S2 Pairwise Dice coefficient comparing the similarity between functional annotations using the 1kG template. . . . . . . . . . . . . . . . . . . . . . . . . . . . . . . . . . . . . . . 26
S3 Pairwise genetic correlation coefficient $r_g$ using LDSC score regression[83] across the full phenotype panel. . . . . . . . . . . . . . . . . . . . . . . . . . . . . . . . . . . . . . 27
S4 Pairwise genetic correlation coefficient $r_g$ using LDSC score regression[83] after pruning at 0.3 favoring phenotypes with higher heritability. . . . . . . . . . . . . . . . . . . . . 28
S5 Contribution of individual functional annotations relative to the full model with 74 annotations showing all phenotypes, i.e., without pruning. Phenotypes are ordered by their polygenicity and annotations are sorted by their Euclidean distance. On the left, next to relative numbers of nucleotides (genome, exon, intron, intergenic), the Spearman rank correlation value between ACS and polygenicity across phenotypes is indicated. The symbols following each functional annotation on the right indicates the significance after Bonferroni correction for the Spearman correlation (significance: ***: p<0.001, **: p<0.01, *: p <0.05, n.s.: p≥0.05). . . . . . . . . . . . . . . . . . . . . . . . . . . 29
S6 Contribution of MAF bin annotations relative to the full model with 74 annotations (phenotype pruning at rg=0.3). Phenotypes are ordered by their polygenicity. Contributions are fairly low with a max value of 0.098 for COVID. . . . . . . . . . . . . . . . . . . 29
S7 Contribution of GWAS catalouge jointly to the full model with 74 annotations across all phenotypes which are ordered by their polygenicity. For all phenotypes, its contribution is greather than one. . . . . . . . . . . . . . . . . . . . . . . . . . . . . . . . . . . . 29

S8 Comparison of (a) heritability fractions and (b) polygenicity estimates between sLD4M regression[1] (incorporating polygenicity estimation) and naive sLDSC style regression (SLD4M infinitesimal). . . . . . 30
S9 $h^2_{SNP}$ fractions from sLD4M[1] or sLDSC for pruned phenotypes. (AF) . . . . . . 30
S10 Contribution of individual functional annotations relative to the full model with 74 annotations. Phenotypes are ordered by their polygenicity remaining after pruning with $r_g$=0.3 favoring phenotypes with higher heritability. The annotation are sorted in the same way as in Fig. 2. Results show log-likelihood values based on infinitesimal assumption. . . . 39
S11 Contribution of individual functional annotations relative to the full model with 74 annotations. Phenotypes are ordered by their polygenicity remaining after pruning with $r_g$=0.3 favoring phenotypes with higher heritability. The annotation are sorted in the same way as in Fig. 2. Results show log-likelihood values based on non-infinitesimal assumption, i.e., with fitted polygenicity. . . . . . 40
S12 Relative log-likelihood-based ACS across pruned phenotypes and functional annotations when using an infinitesimal approach (x-axis) versus and non-infinitesimal approach (y-axis). . . . . . 41
S13 Relative log-likelihood-based ACS across individual pruned phenotypes and functional annotations when using an infinitesimal approach versus and non-infinitesimal approach (AF) . . . . . . 43
S14 Cross-trait SNP heritability cosine similarity heatmaps with traits ordered by decreasing polygenicity. . . . . . 54
S14 Cross-trait SNP heritability cosine similarity heatmaps with traits ordered by decreasing polygenicity. . . . . . 55
S14 Cross-trait SNP heritability cosine similarity heatmaps with traits ordered by decreasing polygenicity. . . . . . 56
S14 Cross-trait SNP heritability cosine similarity heatmaps with traits ordered by decreasing polygenicity. . . . . . 57
S15 Permutation test of cross-trait per-SNP heritability similarity. Traits were binned by -$\log_{10}(\pi)$ (median: 17/17, tertiles: 12/11/11). Histograms show the null distribution of "within-between" differences of cosine similarity matrix from 1e4 label shuffles. Vertical dashed lines indicate the observed within- and between-bin means in the respective panels. 58
S16 Contribution of 221 individual cell types from Finucane *et al.*[84] added jointly to the full model with 74 annotations. Phenotypes are ordered by their polygenicity remaining after pruning with $r_g$=0.3 favoring phenotypes with higher heritability. , There is an increased CNS ACS for SCZ and COG, elevated hematopoietic and gastrointestinal signals for MS and CD, and comparatively modest signals in liver or pancreas for T2D or lipids. . . . . 58

#### Supplementary Tables

S1 Functional annotations with sources. The bulk of the annotations stems from the curated set "baseline-LD v2.3"[7]. Annotations are grouped into higher-level functional domains used in the main text: **Genes** (coding and UTR sequence), **Variant Effect and Literature** (variant-level predicted impact scores and GWAS-informed priors), **Comparative Genomics** (evolutionary conservation scores across species), **Enhancer** (enhancer and super-enhancer annotations), **Promoter and Transcription** (promoter, TSS and transcription-related elements), **Chromatin State** (open or repressed chromatin markers, DHS and footprints), and **Epigenetics** (histone modifications and CpG islands). . . . 21
S2 Overview of phenotypes and GWAS summary statistics panel. The sample size can be the total or the effective sample size. The latter is defined for for case-control studies as $N_{eff} = 4/(1/N_{cases} + 1/N_{controls})$). . . . . . 22
S3 Spearman correlation between ACS and polygenicity estimates for pruned set of phenotypes ($r_g = 0.3$). . . . . . 24

S4 Association between partitioned heritability and polygenicity by genomic region, from a pooled ordinary least squares model with a genomic region $\times \log_{10}(\pi)$ interaction and trait-clustered standard errors (SEs). Corresponding Spearman correlation coefficients between partitioned heritability and polygenicity are also shown. . . . . . . . . . . . . . . 25
S5 Per-domain mean correlation with polygenicity. Spearman $r$ per annotation was Fisher z-transformed. Means and 95% bootstrap CIs (percentile, B=1e4) are reported for both z and r = tanh(z). . . . . . . . . . . . . . . . . . . . . . . . . . . . . . . . . . . . . . . 41
S6 Pairwise differences in mean correlation with polygenicity across functional domains. For each domain, r denotes the Spearman correlation between ACS and phenotype polygenicity (Fisher z-transformed and averaged across traits). $\Delta r = \tanh(\bar{z}_{\mathrm{A}}) - \tanh(\bar{z}_{\mathrm{B}})$ gives the difference in mean correlation (Domain A − Domain B). CIs are 95% bootstrap (percentile, $B = 10^4$). $p$-values are obtained from 1e5 label permutations. An asterisk indicates that the comparison remains significant after Benjamini–Hochberg FDR correction at $q < 0.05$. Positive $\Delta r$ indicates Domain A > Domain B in terms of mean correlation with polygenicity. . . . . . . . . . . . . . . . . . . . . . . . . . . . . . . . 42
S7 90th-percentile of $\log \mathrm{L}_{\mathrm{rel}}$ across annotation-specific relative nucleotide count bins. Bins were chosen to align with nucleotide distributions and maintain counts per bin, showing highest number in mid-size annotations. . . . . . . . . . . . . . . . . . . . . . . . . . . . 42
S8 MAF deciles used in the analysis based on 1000 Genome reference panel. . . . . . . . . 42

### Supplementary Notes – MiXeR

#### Contents

**1 Simple additive genetic model for complex traits**

**2 Probability model for GWAS z-scores**

**3 Likelihood function of the unified MiXeR model**
3.1 Infinitesimal log-likelihood function . . . . . . . . . . . . . . . . . . . . . . . . .
3.2 Likelihood approximation via Monte Carlo sampling . . . . . . . . . . . . . . . . . .
3.3 Likelihood evaluation via characteristic functions and the convolution theorem . . .

**4 Heritability models**
4.1 Full model with functional annotations . . . . . . . . . . . . . . . . . . . . . . . . .
4.2 Log-likelihood function with functional annotations . . . . . . . . . . . . . . . . . .
4.3 Gradient of the log-likelihood function . . . . . . . . . . . . . . . . . . . . . . . . .
4.4 Cosine similarity between heritability models . . . . . . . . . . . . . . . . . . . . .

**5 Implementation details**
5.1 Estimating heritability fractions in exon/intron/intergenic regions using full model . .
5.2 Estimating annotation contribution scores using log-likelihood metric . . . . . . . .
5.3 Sensitivity analysis with causal mixture likelihood . . . . . . . . . . . . . . . . . . .

### 1 Simple additive genetic model for complex traits

Assuming a simple additive genetic model, a quantitative phenotype $y \in \mathbb{R}$ is modeled as a linear combination of genotype dosages $g_i$ with weights $\beta_i$:

$$y = \sum_{i=1}^{\bar{M}} g_i \beta_i + \epsilon_y ,$$

where the index $i$ runs over $\bar{M}$ genetic variants, weights $\beta_i \in \mathbb{R}$ are known as *additive effects of allele substitution*, and the $\epsilon_y \sim \mathcal{N}(0, 1 - h^2)$ term is a normally distributed residual, which reflects contributions from the environment and non-additive genetic effects. The quantitative phenotype $y$ is assumed to be centered and scaled, $E[y] = 0, \mathrm{Var}[y] = 1$, implying that the variance $\mathrm{Var}[\sum_i \beta_i g_i]$ is equal to each trait's SNP heritability, $h^2 \in [0, 1]$. Genotype dosages $g_i$ are assumed to be centered but not scaled, i.e., $g_i \in \{0 - 2f_i, 1 - 2f_i, 2 - 2f_i\}$, where $f_i$ is allele frequency of the $i$-th variant.

Specific prior distribution on effect sizes $\beta_i$ encodes various genetic architectures of complex traits, for example $\beta_i \sim \mathcal{N}(0, \sigma_\beta^2)$ assumption is widely known as *infinitesimal model*. Here $\mathcal{N}(\mu, \sigma^2)$ denotes a Gaussian random variable with mean $\mu$ and variance $\sigma^2$, and we also let $\phi(x; \mu, \sigma^2) = \frac{1}{\sigma\sqrt{2\pi}} \exp(-\frac{(x-\mu)^2}{2\sigma^2})$ denote the probability density of a normal distribution at point $x$. Under infinitesimal model, $\sigma_\beta^2$ parameter is closely related to trait's heritability, $h^2 = \sum_i \sigma_\beta^2 H_i$, where $H_i = 2f_i(1 - f_i)$ is the heterozygosity of $i$-th variant. Indeed, by definition of heritability:

$$h^2 = \mathrm{Var}(\sum_{i=1}^{\bar{M}} g_i \beta_i)/\mathrm{Var}(y).$$

Then, as $g_i$ and $\beta_i$ are independent zero mean random variables, $\mathrm{Var}(y) = 1$, and $\mathrm{Var}(g_i) = H_i$, the expression for variant-based heritability is

$$h^2 = \sum_{i=1}^{\bar{M}} H_i \mathrm{Var}(\beta_i),$$

and, for a genomic region $G \subseteq \{1, \dots, \bar{M}\}$, the partitioned heritability is

$$h_G^2 = \sum_{i \in G} H_i \mathrm{Var}(\beta_i).$$

Various extensions of this basic model are possible by modifying the prior distribution of effect sizes $\beta_i$, allowing for the mixture of null and non-null components, and for distribution of heritability to depend on SNP's allele frequency, functional annotations. We will describe these extensions in the following sections, and refer to them as *heritability models*, as they define how heritability is distributed across the genome.

### 2 Probability model for GWAS z-scores

To infer numeric parameters defining the heritability model (e.g. $\sigma_\beta^2$ as in $\beta \sim \mathcal{N}(0, \sigma_\beta^2)$ we use z-scores ($z_j$) from GWAS summary statistics, where the index $j = 1, \dots, M$ runs over GWAS tag SNPs (imputed or genotyped). Note that the set of GWAS tag SNPs must be a subset of the variants used in modeling additive genetic effects $\beta_i$, with $i = 1, \dots, \bar{M}$.

The following equations relate additive effects of allele substitution $\beta_i$ to $z_j$-scores from GWAS summary statistics.

*Let $\hat{\beta}'_j$ be GWAS estimate of the marginal effect size for $j$-th tag SNP, assessed via univariate linear regression, and $z_j$ be the corresponding z-score, $z_j = \hat{\beta}'_j / \hat{se}(\beta'_j)$. Then*

$$
\begin{aligned}
z_j &= \sqrt{N_j}\delta_j + \epsilon_j, \\
\delta_j &= \sum_{i=1}^{\bar{M}} a_{ij}\beta_i, \text{ where } a_{ij} = \sqrt{H_i} r_{ij}, \\
\epsilon_j &\sim \mathcal{N}(0, \omega_0^2) ,
\end{aligned} \tag{1}
$$

*where $N_j$ is the number of subjects with non-missing genotype information on $j$-th variant; $H_i = 2f_i(1 - f_i)$ is the heterozygosity of $i$-th variant; $r_{ij} = corr(\mathbf{v_i}, \mathbf{v_j})$ is an LD allele count correlation between genotype vectors $\mathbf{v_i}$, and $\mathbf{v_j}$ (two column-vectors running across individuals, containing dosages $g_i$ and $g_j$, of variants i and j respectively, across all individuals), parameter $\omega_0^2$ accounts for non-polygenic inflation in GWAS z-scores, and $\epsilon_j$ denotes a normally distributed residual.*

**Proof**. For this proof only, let $\mathbf{y}$, $\mathbf{e}$, and $\mathbf{v}_i$ denote the phenotype, residual, and genotype-dosage vectors restricted to the subjects included in the univariate regression for tag SNP $j$. Thus all inner products below are taken over the same set of $N_j$ subjects. In the absence of covariates, the least squares estimate $\hat{\beta}'_j$ from regressing $\mathbf{y}$ on $\mathbf{v}_j$ is

$$
\hat{\beta}'_j = \frac{\mathbf{v_j^T y}}{\mathbf{v_j^T v_j}}.
$$

Substituting the additive model $\mathbf{y} = \sum_{i=1}^{\bar{M}} \mathbf{v}_i\beta_i + \mathbf{e}$ gives

$$
\hat{\beta}'_j = \sum_{i=1}^{\bar{M}} \frac{\mathbf{v}_j^T\mathbf{v}_i}{\mathbf{v}_j^T\mathbf{v}_j}\beta_i + \frac{\mathbf{v}_j^T\mathbf{e}}{\mathbf{v}_j^T\mathbf{v}_j}.
$$

As $N_j \to \infty$, the sample covariance ratio converges to the population quantity:

$$
\frac{\mathbf{v}_j^T\mathbf{v}_i}{\mathbf{v}_j^T\mathbf{v}_j} \simeq \frac{\mathrm{Cov}(v_i, v_j)}{\mathrm{Var}(v_j)} = \sqrt{\frac{H_i}{H_j}} r_{ij}.
$$

Therefore

$$
\hat{\beta}'_j \simeq \sum_{i=1}^{\bar{M}} \sqrt{\frac{H_i}{H_j}} r_{ij}\beta_i + \frac{\mathbf{v}_j^T\mathbf{e}}{N_j H_j}.
$$

To connect this with the z-score, we use two standard approximations for the univariate regression on SNP $j$:

$$
\mathbf{v}_j^T\mathbf{v}_j \simeq N_j H_j, \qquad \sigma_{e,j}^2 \simeq 1,
$$

where $\sigma_{e,j}^2$ is the residual variance of that marginal regression. The second approximation uses that the phenotype is standardized, $\mathrm{Var}(y) = 1$, and any single SNP explains only a negligible fraction of phenotypic variance. Therefore the usual univariate OLS standard error satisfies

$$
\hat{se}(\beta'_j) \simeq \sqrt{\frac{\sigma_{e,j}^2}{\mathbf{v}_j^T\mathbf{v}_j}} \simeq \frac{1}{\sqrt{H_j N_j}}.
$$

Defining

$$\epsilon_j = \frac{\mathbf{v}_j^T \mathbf{e}}{\sqrt{H_j N_j}},$$

we obtain

$$z_j = \frac{\hat{\beta}'_j}{\hat{se}(\beta'_j)} \simeq \sqrt{N_j} \sum_{i=1}^{\bar{M}} \sqrt{H_i} r_{ij} \beta_i + \epsilon_j.$$

This is Eq. (1), with $a_{ij} = \sqrt{H_i} r_{ij}$ and $\epsilon_j \sim \mathcal{N}(0, \omega_0^2)$. ■

In a matrix form:

$$\mathbf{z} = \mathbf{N}^{1/2}\, \boldsymbol{R}\, \mathbf{H}^{1/2}\, \boldsymbol{\beta} + \boldsymbol{\epsilon}, \qquad \boldsymbol{\epsilon} \sim \mathcal{N}(\mathbf{0}, \omega_0^2 I_M), \tag{2}$$

where

$\mathbf{z} = (z_1, \ldots, z_M)^\top$, vector of GWAS z-scores

$\mathbf{N}^{1/2} = \mathrm{diag}\big(\sqrt{N_1}, \ldots, \sqrt{N_M}\big)$, $N_j$ is per-SNP effective sample size for tag SNP $j$,

$\mathbf{H}^{1/2} = \mathrm{diag}\big(\sqrt{H_1}, \ldots, \sqrt{H_{\bar{M}}}\big)$,

$H_i = 2f_i(1 - f_i)$ is heterozygosity of the $i$-th modeled variant,

$\mathbf{R} = (r_{ij})_{j=1,\ldots,M,\; i=1,\ldots,\bar{M}}$

LD correlation matrix between tag SNPs (rows) and modeled variants (columns),

$\boldsymbol{\beta} = (\beta_1, \ldots, \beta_{\bar{M}})^\top$,

$\boldsymbol{\epsilon} \sim \mathcal{N}(\mathbf{0}, \omega_0^2 I_M)$ is independent noise.

Note that the heterozygosity vector $\mathbf{H}$ has length $\bar{M}$, matching the modeled-variant index of the columns in $\mathbf{R}$. When the set of tag SNPs coincides with the set of modeled variants ($M = \bar{M}$), $\mathbf{R}$ reduces to the familiar symmetric LD correlation matrix.

Let $\mathcal{T} \subseteq \{1, \ldots, \bar{M}\}$ denote the subset of modeled variants with available GWAS z-scores, so that $|\mathcal{T}| = M$. Inverse LD weighting is then defined using only LD between tag SNPs:

$$\ell_j^{(w)} = \sum_{i=1}^{\bar{M}} [i \in \mathcal{T}] r_{ij}^2 = \sum_{i \in \mathcal{T}} r_{ij}^2, \qquad w_j = \frac{1}{\ell_j^{(w)}}, \qquad j = 1, \ldots, M.$$

That is, while the MiXeR likelihood involves all modeled variants $i = 1, \ldots, \bar{M}$ through sums such as $\sum_{i=1}^{\bar{M}} a_{ij}\beta_i$, the inverse-LD weight for tag SNP $j$ is computed only from variants with available z-scores.

### 3 Likelihood function of the unified MiXeR model

Unified MiXeR model defines per-SNP parameters of the $p(\beta_i)$ prior via spike-and-slab distribution:

$$\beta_i \sim (1 - \pi_i)\delta_0 + \pi_i \mathcal{N}(0, \sigma_i^2)\;, \tag{3}$$

where $\pi_i \in [0, 1]$ defines weights in a two-component mixture distribution, and is interpreted as the prior probability of $i$-th genetic variant to have a non-zero effect, and $\sigma_i^2$ is prior variance of its effect size. $\delta_0$ indicates a distribution with probability mass at 0 (the Dirac delta function). This is a generalization of the standard MiXeR model, where $\pi_i = \pi_1$ and $\sigma_i^2 = \sigma_\beta^2$ are constant across all variants.

Below we derive the likelihood function for the unified MiXeR model, as a function of model parameters $\theta = \{\pi_i, \sigma_i^2, \omega_0^2\}$ (including $\omega_0^2$ accounts for non-polygenic inflation in GWAS z-scores). Unified MiXeR model is not intended to be fitted directly, but rather serves as a general framework to derive likelihood functions for various MiXeR model extensions, such as standard MiXeR with constant $\pi_i = \pi_1$ and $\sigma_i^2 = \sigma_\beta^2$ (Eq. (3)), functional MiXeR with $\sigma_i^2$ defined via functional annotations (see next section), and others.

We now introduce the latent variables $u_i \in \{0, 1\}$ given by a Bernoulli distribution, $p(u_i) = Bern(u_i|\pi_i)$, so that the full probabilistic model can be written as follows:

$$
\begin{aligned}
&p(z_j, \vec{\beta}, \vec{u}|\theta) = p(z_j|\vec{\beta}, \theta) \cdot p(\vec{\beta}|\vec{u}, \theta) \cdot p(\vec{u}|\theta) \;, \\
&p(z_j|\beta_1, \dots, \beta_{\bar{M}}, \theta) = \phi\Big(z_j; \sqrt{N_j} \sum_{i=1}^{\bar{M}} a_{ij}\beta_i, \omega_0^2\Big) \;, \\
&p(\beta_i|u_i = 0, \theta) = \phi(\beta_i; 0, 0) \;, \quad p(\beta_i|u_i = 1, \theta) = \phi(\beta_i; 0, \sigma_i^2) \;, \\
&p(u_i|\theta) = Bern(u_i|\pi_i) \;.
\end{aligned}
$$

A tricky part here is that $z_j$ may depend on multiple $\beta_i$. After observing $\vec{z} = (z_1, \dots, z_M)^T$, we are aiming to do inference on $\theta$ by the maximum likelihood:

$$p_j(z_j|\theta) = \int_u \int_\beta p(z_j, \vec{\beta}, \vec{u}|\theta) du d\beta \;, \tag{4}$$

$$\log L(\boldsymbol{z}|\theta) = \sum_j w_j \log p_j(z_j|\theta) \to \max_\theta, \tag{5}$$

where weights $w_j$ are introduced to avoid over-counting contribution from large LD blocks, with weights $w_j$ computed either from random pruning, or from inverse LD weighting, $w_j = 1/\ell_j^{(w)}$. On a technical note, it is impossible to store all elements of the $a_{ij}$ matrix ($a_{ij} = \sqrt{H_i} r_{ij}$) due to its size. Nevertheless, most of the elements are close to zero, and we use a sparse matrix (e.g. Compressed Sparse Row (CSR) format) to store all $a_{ij}$ with $r_{ij}^2 \geq r_{min}^2$, by default $r_{min}^2 = 0.01$, considering all pairs of variants within each chromosome regardless of the distance between them, but neglecting potential correlations between variants on different chromosomes.

#### 3.1 Infinitesimal log-likelihood function

An important special case of the unified MiXeR model is the infinitesimal constraint $\pi_i = 1$ for all modeled variants. In that case the latent variables $u_i$ are no longer needed and $\beta_i \sim \mathcal{N}(0, \sigma_i^2)$. Because the genetic effects are independent and centered, the marginal distribution of $z_j$ remains Gaussian:

$$p_j(z_j|\theta) = \phi(z_j; 0, s_j^2), \text{ where } s_j^2 = \omega_0^2 + N_j \sum_{i=1}^{\bar{M}} a_{ij}^2 \sigma_i^2. \tag{6}$$

Therefore the corresponding weighted log-likelihood approximation takes the form

$$\log L(\boldsymbol{z}|\theta) = \sum_j w_j \log \phi\Big(z_j; 0, \omega_0^2 + N_j \sum_{i=1}^{\bar{M}} a_{ij}^2 \sigma_i^2\Big), \tag{7}$$

where $\phi(z; 0, s^2) = \frac{1}{\sqrt{2\pi}s} e^{-z^2/2s^2}$ stands for the probability density function of centered normal distribution.

#### 3.2 Likelihood approximation via Monte Carlo sampling

A simple way of computing (5) is to approximate the integral by drawing a large number (e.g. $K = 20,000$) samples from $p(u_i)$ for each latent variable $u_i$. For $j$-th variant and $k$-th sampling iteration, let $U_{jk}$ be the set of variants with $u_i = 1$. For each realization of latent variables the distribution over $p(z_j|U_{jk}, \theta)$ becomes normal zero-mean distribution with a simple analytical formula for its variance:

$$p(z_j|U_{jk}, \theta) = \phi(z_j; 0, \Sigma^2_{jk}) \,,$$
$$\Sigma^2_{jk} = \omega_0^2 + N_j \sum_{i \in U_{jk}} a_{ij}^2 \sigma_i^2, \ .$$

With this approximation, the per-SNP likelihood term in (4) is estimated by Monte Carlo sampling, leading to

$$\log L \approx \sum_j w_j \log\left(\frac{1}{K} \sum_k \phi(z_j; 0, \omega_0^2 + N_j \sum_{i \in U_{jk}} a_{ij}^2 \sigma_i^2)\right) , \quad (8)$$

where $\phi(z; 0, s^2) = \frac{1}{\sqrt{2\pi}s} e^{-z^2/2s^2}$ stands for the probability density function of centered normal distribution. In some cases it is useful to implement right-censoring of large z-scores, replacing $\phi(z; 0, s^2)$ with $q(z; 0, s^2)$ for z-scores that exceed certain threshold $|z_j| \geq z_{max}$, with $q(z_j; 0, s^2) = 2\Phi(-z_{max}; 0, s^2) = erfc\left(\frac{z_{max}}{\sqrt{2}s}\right)$. A typical value for $z_{max}$ is 5.45, which corresponds to conventional genome-wide significance threshold $\alpha = 5 \times 10^{-8}$.

#### 3.3 Likelihood evaluation via characteristic functions and the convolution theorem

To handle the general case $\pi_i \neq 1$, we use a characteristic-function representation of the likelihood based on Fourier inversion. The title of this subsection refers to the convolution theorem: because $z_j$ is represented as a sum of independent random contributions, its characteristic function factorizes into a product of characteristic functions, which is the Fourier-domain counterpart of convolution in the original variable. We briefly recall the generic ingredients of this construction. For a random variable with probability density $p(z)$ and characteristic function

$$\psi(t) = \mathrm{E}[e^{itz}],$$

the inverse Fourier transform gives

$$p(z_0) = \frac{1}{2\pi} \int_{-\infty}^{\infty} e^{-itz_0} \psi(t) dt$$
$$= \frac{1}{2\pi} \int_{-\infty}^{\infty} \cos(tz_0) \psi(t) dt - \frac{i}{2\pi} \int_{-\infty}^{\infty} \sin(tz_0) \psi(t) dt.$$

If $p(z)$ is symmetric around zero, then $\psi(t)$ is an even function and the sine term vanishes, so

$$p(z_0) = \frac{1}{\pi} \int_0^{\infty} \cos(tz_0) \psi(t) dt.$$

We now translate this generic identity to the likelihood calculation from the previous section. Under the unified MiXeR model, for a fixed tag SNP $j$ we can write

$$z_j = \epsilon_j + \sum_{i=1}^{\bar{M}} \xi_{ij},$$

where $\epsilon_j \sim \mathcal{N}(0, \omega_0^2)$ and

$$\xi_{ij} = \sqrt{N_j} a_{ij} \beta_i, \qquad \beta_i \sim (1-\pi_i)\delta_0 + \pi_i \mathcal{N}(0, \sigma_i^2).$$

Therefore the contribution of the $i$-th modeled variant to $z_j$ is

$$\xi_{ij} \sim \begin{cases} 0, & 1-\pi_i, \\ \mathcal{N}(0, N_j a_{ij}^2 \sigma_i^2), & \pi_i. \end{cases}$$

Because the terms are independent, the characteristic function of $z_j$ factorizes:

$$\psi_{z_j}(t) = \psi_{\epsilon_j}(t) \prod_{i=1}^{\bar{M}} \psi_{\xi_{ij}}(t).$$

$$\begin{aligned} \psi_{\xi_{ij}}(t) &= (1-\pi_i) + \pi_i e^{-\frac{1}{2} t^2 N_j a_{ij}^2 \sigma_i^2}, \\ \psi_{\epsilon_j}(t) &= e^{-\frac{1}{2} t^2 \omega_0^2}. \end{aligned}$$

Substituting these expressions into the inverse Fourier formula yields the exact per-SNP likelihood

$$p(z_j|\theta) = \frac{1}{\pi} \int_0^\infty \cos(t z_j)\, e^{-\frac{1}{2} t^2 \omega_0^2} \prod_{i=1}^{\bar{M}} \left[ (1-\pi_i) + \pi_i e^{-\frac{1}{2} t^2 N_j a_{ij}^2 \sigma_i^2} \right] dt,$$

and the full objective remains

$$\log L(\boldsymbol{z}|\theta) = \sum_j w_j \log p(z_j|\theta).$$

To evaluate this integral numerically, we map the semi-infinite interval to a bounded domain and then apply adaptive cubature. In practice, we use the Cubature package of Steven G. Johnson for adaptive multidimensional integration over hypercubes.[1]

For a semi-infinite interval one may use the change of variables $x = a + \frac{u}{1-u}$:

$$\int_a^\infty f(x)dx = \int_0^1 f\Big(a + \frac{u}{1-u}\Big) \frac{1}{(1-u)^2} du.$$

For a two-sided infinite interval one may use $x = \frac{u}{1-u^2}$:

$$\int_{-\infty}^\infty f(x)dx = \int_{-1}^1 f\Big(\frac{u}{1-u^2}\Big) \frac{1+u^2}{\left(1-u^2\right)^2} du.$$

[1] `https://github.com/stevengj/cubature`

Note the Jacobian factors multiplying $f(\cdots)$ in both integrals, and also that the integration limits are different in the two cases.

**Bounded re-parametrization of improper integrals.** For the convolution likelihood above, only the one-sided transformation is needed because symmetry has already reduced the Fourier inversion to the interval $[0, \infty)$. Using $t = \frac{u}{1-u}$, we obtain

$$p(z_j|\theta) = \frac{1}{\pi}\int_0^1 \cos\left(\frac{u}{1-u}z_j\right)\psi_{z_j}\left(\frac{u}{1-u}\right)\frac{1}{(1-u)^2}\,du.$$

This bounded-domain form is what we pass to the adaptive cubature routine.

### 4 Heritability models

This section describes *full model* accounting non-uniform distribution of heritability across functional annotations. This extends *core model*, which is defined by $\beta_i \sim \mathcal{N}(0, \sigma_\beta^2 H_i^S)$, where the $S$ parameter controls the effect size distribution with regards to allele frequency, similarly to LDAK and SumHer models. We note that effect sizes $\beta_i$ are defined w.r.t. centered but not scaled genotypes $g_i$, so that $\mathrm{Var}(g_i) = H_i = 2f_i(1-f_i)$, thus under our core model trait's SNP heritability $h^2 = \sum_{i=1}^{M} \sigma_\beta^2 H_i^{S+1}$. The original GCTA model assumes $S = -1$, so that each SNP contributes equally to heritability regardless of its allele frequency.

The full model extends the core heritability model by retaining its MAF-dependent factor $H_i^S$, and modulating it with an additive combination of contributions from functional annotation categories, so that annotations determine how heritability is redistributed across variants beyond the core allele-frequency dependence.

#### 4.1 Full model with functional annotations

In the full model per-SNP variance $\sigma_i^2$ is defined by the following expression:

$$\sigma_i^2 = \left(\sigma_{A,0}^2 + \sum_{p=1}^{N_a}[i \in A_p]\sigma_{A,p}^2\right)H_i^S \qquad (9)$$

The index $p$ runs across functional annotation categories $\{A_1, A_2, \dots, A_{N_a}\}$, while the parameter $\sigma_{A,p}^2$ represents the contribution of the $p$-th annotation category to the variance of $i$-th genetic variant, given that the $i$-th variant belongs to the $p$-th category, as specified in 9 via the indicator variable $[i \in A_p]$, using square brackets notation to transform a logical statement from true or false into 1 and 0, respectively. If genetic variant $i$ belongs to multiple annotation categories, the variance parameters of those annotations will be added together in an additive manner. To simplify notation we will introduce "base" category $A_0$ including all variants, and run index $p = 0, \dots, N_a$:

$$\sigma_i^2 = \sum_{p=0}^{N_a}[i \in A_p]\sigma_{A,p}^2 \qquad (10)$$

In a matrix notation this is equivalent to $\boldsymbol{\beta} \sim \mathcal{N}(\mathbf{0}, \Sigma_\beta)$ where

$$\Sigma_\beta \;=\; \mathrm{diag}\!\left(U\,\boldsymbol{\sigma}_A^2\right),$$

$$U = (u_{ip}) \in \{0,1\}^{\bar{M}\times(N_a+1)}, u_{ip} = 1 \text{ iff SNP } i \in A_p,$$

$$\boldsymbol{\sigma}_A^2 = (\sigma_{A,0}^2, \sigma_{A,1}^2, \dots, \sigma_{A,N_a}^2)^\top,$$

$$\text{so that} \quad \sigma_i^2 = \sum_{p=0}^{N_a} u_{ip}\, \sigma_{A,p}^2,$$

$$\text{and therefore} \quad \Sigma_\beta = \operatorname{diag}(\sigma_1^2, \dots, \sigma_{\bar{M}}^2), \quad \operatorname{Cov}(\beta_i, \beta_j) = 0 \ (i \neq j).$$

#### 4.2 Log-likelihood function with functional annotations

Under the infinitesimal constraint $\pi_i = 1$, equation (7) specializes to the functional-annotation model as follows. The log-likelihood function of observing GWAS z-scores $\mathbf{z} = (z_j)$ given model parameters, $\theta = (\{\sigma_{A,p}^2\}, \omega_0)$, can be written as follows:

$$\begin{aligned} \log L(\mathbf{z}|\theta) &= \sum_j w_j \log \phi(z_j; 0, \omega_0^2 + N_j \sum_{i=1}^{\bar{M}} a_{ij}^2 \sigma_i^2) = \\ &= \sum_j w_j \log \phi(z_j; 0, \omega_0^2 + N_j \sum_{p=0}^{N_a} \sum_{i=1}^{\bar{M}} a_{ij}^2 [i \in A_p] H_i^S \sigma_{A,p}^2) = \\ &= \sum_j w_j \log \phi(z_j; 0, \omega_0^2 + N_j \sum_{p=0}^{N_a} \ell_{jp} \sigma_{A,p}^2), \end{aligned} \tag{11}$$

where at the last step we've used expression for $a_{ij}$, and introduced heterozygosity-adjusted LD scores $\ell_{jp}$ for each functional category $p$ and tag SNP $j$:

$$\ell_{jp} = \sum_{i=1}^{\bar{M}} a_{ij}^2 [i \in A_p] H_i^S = \sum_{i \in A_p} H_i^{S+1} r_{ij}^2. \tag{12}$$

Weights $w_j$ are introduced to avoid over-counting contribution from large LD blocks, and $w_j$ can be computed from inverse LD weighting ($w_j = 1/\ell_j^{(w)}$, default), or as probabilities of selecting $j$-th SNP in random pruning procedure; $\phi(z; 0, s^2) = \frac{1}{\sqrt{2\pi s}} e^{-z^2/2s^2}$ stands for the probability density function of centered normal distribution, $N_j$ is GWAS sample size for SNP $j$.

Under fixed value of $S$ parameter, equation (11), $E[z_j^2] = \omega_0^2 + N_j \sum_{p=0}^{N_a} \ell_{jp} \sigma_{A,p}^2$, is closely related to stratified LDSC regression model, with regression parameters $\vec{x} = \sigma_{A,p}^2$ and intercept $\omega_0^2$.

#### 4.3 Gradient of the log-likelihood function

For optimization, we re-parametrize $\sigma_{A,p}^2 = e^{x_{A,p}}$ to ensure positive value for $\sigma_{A,p}^2$ while $x_{A,p}$ parameter can be unconstrained. With this re-parametrization, the gradients can be computed as follows:

$$\frac{\partial \log L}{\partial x_{A,p}} = \frac{\partial\big(\sum_j w_j \log \phi(z_j; 0, s_j^2)\big)}{\partial x_{A,p}} = \sum_j \frac{w_j}{p_j} \frac{\partial p_j}{\partial x_{A,p}},$$

where $p_j = \phi(z_j; 0, s_j^2)$, $s_j^2 = \omega_0^2 + \sum_{p=0}^{N_a} v_{jp} e^{x_{A,p}}$, and $v_{jp} = N_j \ell_{jp}$. Derrivative of the gaussian distribution density w.r.t. variance parameter is as follows:

$$\frac{\partial \phi(z; 0, s^2)}{\partial s^2} = \frac{z^2 - s^2}{2s^4} \phi(z; 0, s^2),$$

therefore

$$\frac{\partial \log L}{\partial x_{A,p}} = \sum_j \frac{w_j}{p_j} \frac{(z_j^2 - s_j^2) p_j}{2s_j^4} \frac{\partial s_j^2}{\partial x_{A,p}} = \sum_j \frac{w_j (z_j^2 - s_j^2)}{2s_j^4} N_j \ell_{jp} \sigma_{A,p}^2.$$

Similarly, for $\omega_0^2 = e^{x_0}$,

$$\frac{\partial \log L}{\partial x_0} = \sum_j \frac{w_j}{p_j} \frac{(z_j^2 - s_j^2) p_j}{2s_j^4} \frac{\partial s_j^2}{\partial x_0} = \sum_j \frac{w_j (z_j^2 - s_j^2)}{2s_j^4} \omega_0^2.$$

##### 4.4 Cosine similarity between heritability models

For two models, denoted $A$ and $B$, let $h_{Ai}^2$ and $h_{Bi}^2$ denote the SNP-specific heritability contributions implied by the corresponding heritability models (for example, heritability models for two different traits):

$$\begin{aligned} h_{Ai}^2 &= H_i \pi_{iA} \sigma_{Ai}^2, \\ h_{Bi}^2 &= H_i \pi_{iB} \sigma_{Bi}^2 \end{aligned} \tag{13}$$

This provides a summary measure of similarity between the two heritability distributions across the genome: if the two models allocate heritability to similar sets of variants then the cosine similarity is high, whereas if they imply different heritability distributions then the cosine similarity is low. We then estimate cosine similarity between vectors $\vec{u} = (\sqrt{h_{A0}^2}, \ldots, \sqrt{h_{AM}^2})$ and $\vec{v} = (\sqrt{h_{B0}^2}, \ldots, \sqrt{h_{BM}^2})$ as

$$\cos_{\text{sim}}(A, B) = \frac{\sum_i \sqrt{h_{Ai}^2} \sqrt{h_{Bi}^2}}{\sqrt{\sum_i h_{Ai}^2} \cdot \sqrt{\sum_i h_{Bi}^2}}$$

#### 5 Implementation details

##### 5.1 Estimating heritability fractions in exon/intron/intergenic regions using full model

This subsection describes how total SNP heritability is partitioned across exon, intron, and intergenic regions. Throughout this subsection, the *complete set of functional categories* is defined as the union of LDSC baseline v2.3 annotations and RefSeq categories (exon, intron, intergenic). The procedure described below was repeated independently for each analyzed trait. The implementation proceeds as follows:

1. **Fit the full model on the complete annotation set.** Let $A_1, \ldots, A_{N_a}$ denote the complete set of functional categories, with three of these categories corresponding to RefSeq exon, intron,

and intergenic regions; denote these three RefSeq categories by $R_1, R_2, R_3$. The per-SNP variance model is the additive functional-annotation model from (10), with the MAF-dependence parameter constrained to $S = 0$:

$$\sigma^2_{full,i} = \sum_{p=1}^{N_a} [i \in A_p] \sigma^2_{A,p},$$

and parameters are optimized by the same unconstrained maximum-likelihood procedure as in the core model.

2. **Aggregate the fitted full model back to exon, intron, and intergenic regions.** Although the full model is fitted using the complete annotation set, this model is then used to estimate heritability fractions withi the three RefSeq categories. Using the variant-based heritability identity from Section 1, the fitted full model implies per-SNP heritability contribution $H_i \hat{\sigma}^2_{full,i}$, and the regional contributions are

$$\hat{h}^2_{full}(R_q) = \sum_{i \in R_q} \hat{h}^2_{full,i}, \qquad \hat{h}^2_{full,i} = H_i \hat{\sigma}^2_{full,i}, \qquad q \in \{1, 2, 3\}.$$

The total full-model heritability is

$$\hat{h}^2_{full} = \sum_{i=1}^{\bar{M}} \hat{h}^2_{full,i}.$$

The reported heritability fractions are

$$\widehat{\text{frac}}_{full}(R_q) = \frac{\hat{h}^2_{full}(R_q)}{\hat{h}^2_{full}}, \qquad q \in \{1, 2, 3\}.$$

**Optimization procedure.** For both the core and full models, parameters are estimated by direct maximization of the univariate log-likelihood (11). To ensure positivity of variance parameters, the optimization is performed in log scale,

$$\sigma^2_{A,p} = e^{x_{A,p}}, \qquad \omega^2_0 = e^{x_0},$$

so that the resulting optimization problem is unconstrained. In the MATLAB implementation, optimization is carried out with `fminunc` using the trust-region algorithm and analytical first-order derivatives. The corresponding gradient formulas are those given in the subsection *Gradient of the log-likelihood function*. In practice, an LDSC-style weighted regression is used to provide initial values, after which the likelihood is refined by direct optimization.

**Algorithm 1:** Workflow for estimating full-model heritability fractions in exon, intron, and intergenic regions

**Data:** GWAS z-scores $z_j$, sample sizes $N_j$, LD correlations $r_{ij}$, heterozygosities $H_i$, complete annotation set $A_p$, RefSeq regions $R_q \in \{\text{exon, intron, intergenic}\}$
**Result:** Full-model heritability fractions $\widehat{\text{frac}}_{full}(R_q)$

```
/* Full model fit                                                     */
```

**foreach** *tag SNP j, annotation p* **do**
  $\ell_{jp} \leftarrow \sum_{i \in A_p} H_i r_{ij}^2$;
**end**
Initialize $(\omega_0^2, \sigma_{A,p}^2)$ from LDSC-style weighted regression on $\ell_{jp}$;
Optimize full-model log-likelihood (11) in log-parameter space;

```
/* Regional readout                                                   */
```

**foreach** *variant i* **do**
  $\hat{h}^2_{full,i} \leftarrow H_i \hat{\sigma}^2_{full,i}$;
**end**
**foreach** *RefSeq region $R_q$* **do**
  $\widehat{\text{frac}}_{full}(R_q) \leftarrow \Big( \sum_{i \in R_q} \hat{h}^2_{full,i} \Big) / \Big( \sum_{i=1}^{\bar{M}} \hat{h}^2_{full,i} \Big)$;
**end**

#### 5.2 Estimating annotation contribution scores using log-likelihood metric

The goal is to quantify how much of the *full* model likelihood improvement over the *core model* can already be captured by a *partial* model that includes one annotation at a time. The procedure described below was repeated independently for each analyzed trait. Model fitting then proceeds in three stages:

1. **Core model.** The core model is the $S = 0$ special case of the framework from Section 2, with a single global variance parameter:
$$\sigma^2_{core,i} = \sigma^2_{A,0}.$$
This defines the likelihood $\log L_{core}$.

2. **Partial model for annotation $C$.** For each focal annotation $C$, a separate partial model is fitted:
$$\sigma^2_{part(C),i} = \sigma^2_{core}. + [i \in C]\sigma^2_C.$$
Equivalently, this is the functional-annotation model (10) restricted to a shared core component plus one focal annotation. The variance attributable to membership in $C$ is captured by $\sigma^2_C$, and the resulting likelihood is denoted $\log L_{part(C)}$.

3. **Full model.** The full model uses the complete set of functional categories, i.e. RefSeq plus LDSC baseline v2.3:
$$\sigma^2_{full,i} = \sigma^2_{A,0} + \sum_{p=1}^{N_{full}} [i \in A_p]\sigma^2_{A,p}.$$
This gives the likelihood $\log L_{full}$.

In the current implementation, these models are fitted by direct maximization of the univariate log-likelihood (11), using the same unconstrained log-parameterization and analytical gradients as in Section 3.1. An LDSC-style weighted regression is used only to provide initial values for the subsequent likelihood optimization. The readout of interest is the fraction of the total full-model log-likelihood improvement that is already explained by the partial model:

$$\text{ACS}(C) = \frac{\log L_{part(C)} - \log L_{core}}{\log L_{full} - \log L_{core}}.$$

We refer to ACS($C$) as the *annotation contribution score* of annotation $C$. Values close to 1 indicate that the focal annotation alone explains most of the likelihood improvement obtained by the complete full model, while values close to 0 indicate little additional contribution beyond the core model.

**Algorithm 2:** Workflow for annotation contribution scores

**Data:** GWAS z-scores $z_j$, sample sizes $N_j$, LD correlations $r_{ij}$, heterozygosities $H_i$, complete annotation set $A_p$
**Result:** Annotation contribution scores ACS($C$) for all focal annotations $C$

```
/* Reference models */
```

Fit the core model and record $\log L_{core}$;
Fit the full model and record $\log L_{full}$;

```
/* Partial models */
```

**foreach** *focal annotation C* **do**
  Fit the partial model for $C$ and record $\log L_{part(C)}$;
  Compute ACS($C$) $\leftarrow \frac{\log L_{part(C)} - \log L_{core}}{\log L_{full} - \log L_{core}}$;
**end**

#### 5.3 Sensitivity analysis with causal mixture likelihood

Sensitivity analysis is performed by replacing the infinitesimal prior ($\pi_i = 1$) with the causal mixture prior from (3), while keeping the fitted per-SNP heritability profile fixed. This is done separately for the core model and for each fitted partial or full model.

Let $\hat{\sigma}_i^2$ denote the per-SNP variance profile estimated under the infinitesimal model. For a causal mixture model with prior probability $\pi_i$ of a non-zero effect, the non-null variance is re-parametrized as

$$\sigma_i^{2,mix} = \frac{\hat{\sigma}_i^2}{\pi_i},$$

so that

$$\pi_i \sigma_i^{2,mix} = \hat{\sigma}_i^2.$$

Therefore the expected per-SNP contribution to heritability,

$$H_i \pi_i \sigma_i^{2,mix} = H_i \hat{\sigma}_i^2,$$

is kept fixed while varying the degree of polygenicity.

The practical procedure is as follows:

1. **Core-model sensitivity.** For the core model, a single global parameter $\pi$ is used:

$$\pi_i = \pi \qquad \text{for all } i.$$

   The implementation evaluates the unified univariate likelihood (5) on a logarithmic grid $\pi \in [10^{-4}, 1]$ and interpolates the resulting curve to obtain the minimizing value $\hat{\pi}_{core}$.

2. **Full-model sensitivity.** For the complete full model, the same single-parameter grid search is applied:

$$\pi_i = \pi \qquad \text{for all } i.$$

This yields $\hat{\pi}_{full}$ and the corresponding optimized likelihood.

3. **Partial-model sensitivity (AI-MiXeR parameterization).** For a focal annotation $C$, the current implementation uses two polygenicity parameters:

$$\pi_i = \begin{cases} \pi_{in}, & i \in C, \\ \pi_{out}, & i \notin C. \end{cases}$$

In practice, $\pi_{out}$ is fixed to the best-fitting core-model value $\hat{\pi}_{core}$, and $\pi_{in}$ is optimized on the same logarithmic grid. This yields a partial-model sensitivity analysis with separate polygenicity inside and outside the focal annotation, while preserving the infinitesimal-model heritability profile through the constraint $\pi_i \sigma_i^{2,mix} = \hat{\sigma}_i^2$.

4. **The *annotation contribution score* of annotation $C$** is computed as before, using causal mixture likelihood function:

$$\mathrm{ACS}(C) = \frac{\log L_{part(C)} - \log L_{core}}{\log L_{full} - \log L_{core}}.$$

At the last step, likelihood evaluation follows the causal-mixture machinery developed earlier in Section 2. In the current implementation, the native plugin combines the characteristic-function / convolution approach for tag SNPs with $|z_j| < 5.45$ and the sampling-based approximation from Section 3.2 for tag SNPs with $|z_j| \geq 5.45$, because the latter also supports right-censoring of extreme z-scores. The resulting optimized likelihood values are then compared against the infinitesimal fits to assess robustness of the annotation contribution scores to deviations from the $\pi_i = 1$ assumption.

### Supplementary Results

In this section, we present additional analyses that assess the robustness of our main findings to model assumptions and design choices, and that characterize properties of the functional annotations used in the study. First, we compare polygenicity-informed versus infinitesimal model specifications and cross-check polygenicity estimates across methods. We then describe the properties of individual functional annotations and domains, summarize overlap, size, and composition features of the functional annotation panel.

#### Robustness to Model Assumptions and Polygenicity Estimation

The sLD4M[1] framework leverages fourth moments to simultaneously estimate heritability and polygenicity for functional annotations. This extends sLDSC[2], which operates under the infinitesimal assumption (effectively $\pi = 1$) and estimates only annotation-specific heritability. Because the impact of this assumption is central to our study, we conducted a sensitivity analysis comparing sLD4M to an infinitesimal specification. Specifically, we compared annotation-level heritability fractions from sLD4M (polygenicity-enabled) to sLD4M with the polygenicity component disabled (infinitesimal), allowing a direct output comparison. The resulting estimates were highly correlated ($r = 0.996$; Fig. S8a,S9).

We further evaluated concordance between global polygenicity estimates from univariate MiXeR[3] (one genome-wide estimate per phenotype) and those from sLD4M (which yields a global polygenicity while modeling functional annotations). We observed high correlation across phenotypes (Spearman r = 0.91; Fig. S8b).

We further examined the impact of the infinitesimal assumption on the MiXeR log-likelihood evaluations (i.e., setting $\pi = 1$). To this end, we used annotation-informed MiXeR (AI MiXeR)[4] to allow polygenicity estimation within the Fig. 2 framework. Relative to our infinitesimal fitting, we made two pragmatic preprocessing changes: (i) we increased the minor allele frequency (MAF) filter from 0.005 to 0.05 to reduce instability from very rare variants and improve the normal approximation for test statistics, and (ii) we censored GWAS $z$-scores at $|z|=5.45$ to mitigate undue influence of outliers and heavy tails on the likelihood surface (i.e., Winsorization to stabilize optimization under model misspecification). In addition, to reduce redundancy and computational burden while preserving signal, we applied hard linkage disequilibrium (LD) pruning prior to inverse LD weighting, removing SNPs in high-LD pairs with $r^2>0.8$, and then computed inverse LD weights within the retained set. For polygenicity fitting, we performed a one-dimensional grid search over 30 values (within $[0, -4]$ on a log scale) with spline interpolation to locate the optimum cost (minimum negative log-likelihood). We further assumed the polygenicity outside a given annotation to equal the global (baseline) polygenicity. This yielded two model outputs, one from the infinitesimal specification (Fig. S10) and one from the non-infinitesimal specification (Fig. S11). The resulting ACS estimates were strongly correlated (Fig. S12; r=0.92), with the largest deviations observed for lower-polygenicity phenotypes (Fig. S13). Importantly, these differences did not alter domain-level rankings or the principal conclusions regarding the alignment of comparative genomics and variant-effect annotations with polygenicity.

To test whether the polygenicity ordering and the infinitesimal specification could spuriously induce the pattern observed in Fig. 2, we performed a surrogate analysis based on cross-trait similarity of SNP heritability profiles. For each phenotype, we constructed a genome-wide vector of SNP heritabilities from the stratified infinitesimal fit. We then computed pairwise cross-trait similarities using a cosine similarity applied to square root transformed SNP heritabilities. Similarities were computed genome-wide and separately within RefSeq exonic, intronic, and intergenic strata (see resulting similarity matrices in Fig. S14a–d). We next asked whether traits with similar polygenicity exhibit greater similarity in their per-SNP heritability landscapes than expected by chance. Traits were partitioned into bins by the empirical distribution of -log($\pi$)-transformed (median split or tertiles). We compared the mean within-bin similarity to the mean between-bin similarity and assessed significance using a permutation test that preserves bin sizes while shuffling trait labels. Across 1e−4 permutations, we observed a significant "within-greater-than-between" effect for the genome-wide SNP heritability (Fig. S15; tertiles split: $T_{obs}$=0.056, z=7.28, p=1e−4, bin sizes 12/11/11, median split: $T_{obs}$=0.0528, z=7.35, p=4e−4, bin sizes 17/17). Results were similar for the median split and were robust across RefSeq strata (exon, intron, intergenic; Fig. S14b–d). These findings indicate that traits with similar polygenicity share more similar per-SNP heritability landscapes than expected under label shuffling, arguing against a presentation artifact due to the ordering by polygenicity.

#### Minor-Allele-Frequency Bins as Model Covariates

Regarding MAF bins, we defined our them as deciles that partitioned the genome into ten equally sized MAF bins. Their labels increased with allele frequency (Table S8). Note that we defined custom MAF bins instead of using the ones given in the baseline-LD v2.3 model, as they are based on a MAF cutoff of 0.05, excluding rare SNPs[5]. MAF-bin log-likelihood contributions are modest and trait-specific. Chronic kidney disease shows the largest contributions in rare bins (bin 1=0.58, bin 2=0.23), fasting glucose peaks at high MAF (bin 10=0.14), COVID-19 at mid-to-high MAF (bins 6–7=0.06–0.10), and higher-polygenicity brain phenotypes are essentially flat across MAF (Fig. S6; Table S8). This pattern aligns with[5], where explicit MAF (LD) modeling absorbs frequency-driven artifacts. In practice, our MAF bins might stabilize the stratified model, rather than have large standalone MAF contributions.

#### Functional Annotation Properties and Overlap

To characterize redundancy and overlap among functional annotations, we computed pairwise Dice coefficients for all 74 tracks (Fig. S2). Chromatin-state annotations showed moderate pairwise overlap, reflecting shared features across related states. For example, transcription start site (TSS) or promoter tracks moderately colocalized with promoter-associated histone markers (e.g., H3K4me3, H3K9ac) and CpG islands, whereas broad cell-type union annotations exhibited lower Dice similarity due to the union across heterogeneous tissues and cell types. These patterns confirm that related regulatory features cluster as expected, yet no single chromatin track fully captures the others.

We additionally considered a "GWAS Catalog" annotation track, provided by the UCSC Genome Browser, which aggregates loci reported in the NHGRI-EBI GWAS Catalog[6]. As expected, this track primarily tagged SNPs outside exons, consistent with the predominance of noncoding GWAS associations. It showed relatively low Dice similarity to most other annotations (Fig. S2), indicating that Catalog loci only partially overlap with the functional categories used here. Including the GWAS Catalog track as an additional annotation in the full model improved overall model fit and yielded the largest single-annotation gain in log-likelihood across traits (Fig. S7). However, because this track is derived from GWAS that partially overlap the summary statistics analyzed in the present study, we excluded it from the primary analyses to avoid potential circularity. We retain it as a sensitivity analysis quantifying the relative contribution of previously reported loci compared with functional annotations.

Several variant-effect and literature annotations were originally defined as continuous scores (e.g., CADD, Eigen, ncER, ReMM, PrimateAI). To obtain interpretable discrete categories and reduce the influence of extreme outliers, we binarized these scores into high-impact subsets (e.g., "boosted" tracks or extreme percentile tails; see Materials and Methods). This procedure yields compact, information-rich annotations with reduced genomic coverage. For example, CADD high-impact SNPs had approximately 34.9% exonic content, whereas Eigen-ALL boosted SNPs were about 8.9% exonic (Table S1). Thus, variant-effect annotations span a continuum from relatively exon-enriched to largely noncoding, but as a group they capture functionally prioritized subsets within the broader intronic and intergenic space.

To assess how annotation size influences log-likelihood ACS values, we binned annotations by the fraction of SNPs they cover. Bin boundaries were chosen to match the empirical distribution of annotation sizes while maintaining adequate numbers of annotations per bin (Fig. 3). We then examined the distribution (particularly the 90th percentile) of ACS values within each size bin. ACS peaked in the 10–15% coverage bin, with 15–25% and 3–6% bins close behind, whereas very small and very large categories tended to yield lower ACS values on average (Table S6). These results indicate that mid-sized annotations, such as many conservation- and impact-based tracks, are especially informative per nucleotide, whereas extremely broad annotations (e.g., large histone unions) dilute signal across many SNPs and very small annotations have limited power.

A subset of regulatory annotations not biologically defined as exonic (for example, transcription factor binding site tracks from ENCODE and chromatin-state annotations) showed mild exon enrichment. Several mechanisms likely contribute. First, promoter and transcription windows often extend into the 5′ UTR and first exon, leading regulatory peaks to overlap exonic sequence. Second, unioned peak calls and window-based definitions blur precise boundaries between promoter, UTR, and coding segments. Third, in gene-dense regions, regulatory sequence of one gene can overlap the coding sequence of a neighboring gene. Together, these effects produce modest exonic enrichment in some regulatory tracks and should be considered when interpreting exon-enriched categories in the functional-annotation analyses.

We partitioned fractions of SNP heritability across exons, introns, and IGRs for 34 complex traits and disorders (Table S2; Fig. S4), using the GSA-MiXeR-based annotation framework with the full functional annotation model (all annotation categories) and then aggregating SNP heritability into exons, introns and IGRs, so that regional heritability fractions sum to the genome□wide total.

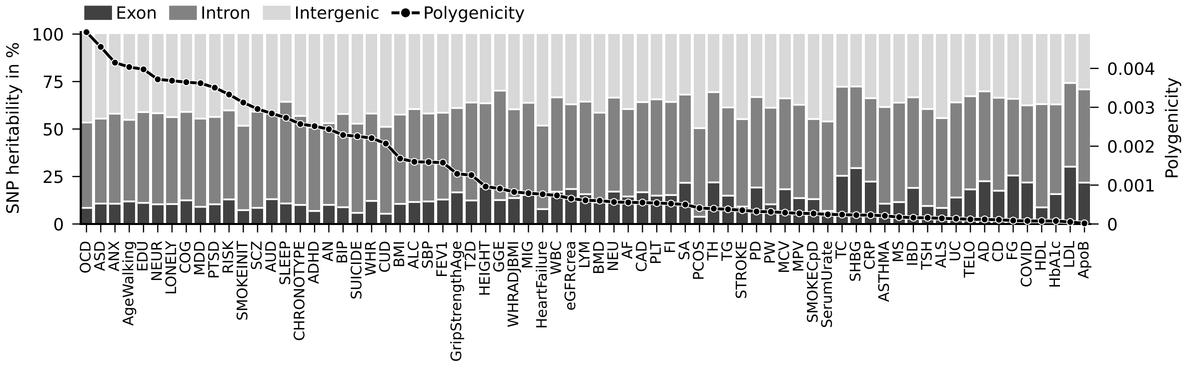


**Figure S1:** Fractions of SNP heritability partitioned across exons, introns and IGRs with phenotypes sorted by polygenicity. The solid line shows polygenicity estimates. With increasing polygenicity, heritability decreases in exons, increases in IGRs, and stays invariant in introns.

**Table S1:** Functional annotations with sources. The bulk of the annotations stems from the curated set "baseline-LD v2.3"[7]. Annotations are grouped into higher-level functional domains used in the main text: **Genes** (coding and UTR sequence), **Variant Effect and Literature** (variant-level predicted impact scores and GWAS-informed priors), **Comparative Genomics** (evolutionary conservation scores across species), **Enhancer** (enhancer and super-enhancer annotations), **Promoter and Transcription** (promoter, TSS and transcription-related elements), **Chromatin State** (open or repressed chromatin markers, DHS and footprints), and **Epigenetics** (histone modifications and CpG islands).

| Functional Annotation | Source |
|---|---|
| **Genes** | |
| Synonymous nucleotide change | 8 |
| Non-synonymous nucleotide change | 8 |
| 5' UTR | 9 |
| Coding exon | 9 |
| 3' UTR | 9 |
| **Variant Effect and Literature** | |
| Missense badness, PolyPhen-2, and Constraint - MPC | 10 |
| Coding variants' pathogenicity - PrimateAI, boosted, 99.9 perc. | 11 |
| Genomic Pretrained Network - GPN-MSA $\leq -7$ | 12 |
| Rare Exome Variant Ensemble Learner - REVEL, boosted, 99.5 perc. | 13 |
| Combined Annotation Dependent Depletion - CADD | 14 |
| Regulatory Mendelian Mutation - ReMM | 15 |
| Regulatory Mendelian Mutation - ReMM, boosted | 15 |
| GWAS catalogue | 9 |
| Non-coding Essential Regulation - ncER, boosted | 16 |
| Functional impact score - Eigen-ALL, boosted | 17 |
| **Comparative Genomics** | |
| Genomic Evolutionary Rate Profiling - GERP RS $\geq 4$ | 9 |
| Zoonomia Constrained, 240 Mammals, phyloP | 7 |
| Zoonomia Constrained, 43 Primates, phastCons | 7 |
| Conserved Primate, phastCons46way | 9 |
| Conserved Mammal, phastCons46way | 9 |
| Conserved Elements, 29 Mammals, phyloP/phastCons (Lindblad-Toh) | 18 |
| Conserved Vertebrate, phastCons46way | 9 |
| **Enhancer** | |

| Functional Annotation | Source |
|---|---|
| Enhancer (Andersson) | 19 |
| Ancient Sequence Age Human Enhancer (Marnetto) | 20 |
| Weak Enhancer (Hoffman) | 21 |
| Human Enhancer (Villar) | 22 |
| Enhancer (Hoffman) | 21 |
| Super Enhancer (Hnisz) | 23 |
| **Promoter and Transcription** | |
| Human Promoter (Villar, ExAC) | 22 |
| Ancient Sequence Age Human Promoter (Marnetto) | 20 |
| Promoter Flanking (Hoffman) | 21 |
| Human Promoter (Villar) | 22 |
| Transcription Start Site (Hoffman) | 21 |
| CTCF Binding Factor (Hoffman) | 21 |
| Promoter (UCSC) | 9 |
| Transcription Factor Binding Site (ENCODE) | 24 |
| Transcribed (Hoffman) | 21 |
| **Chromatin State** | |
| Fetal DNase I Hypersensitivity Sites - DHS (Trynka) | 25 |
| DNase I Hypersensitivity Sites peaks - DHS peaks (Trynka) | 25 |
| Digital Genomic Footprinting - DGF (ENCODE) | 24,26 |
| DNase I Hypersensitivity Sites - DHS (Trynka) | 25 |
| Repressed (Hoffman) | 21 |
| **Epigenetics** | |
| CpG Island (UCSC) | 9 |
| H3K4me1 QTL MaxCPP boosted (Hormozdiari) | 27 |
| Flanking Bivalent Transcription Start Site/Enhancer - BivFlnk (Roadmap) | 28 |
| H3K9ac peaks (Trynka) | 25 |
| H3K4me3 peaks (Trynka) | 25 |
| H3K9ac boosted (Trynka) | 25 |
| H3K9ac peaks boosted (Trynka) | 25 |
| H3K9ac (Trynka) | 25 |
| H3K4me3 (Trynka) | 25 |
| H3K4me1 peaks (Trynka) | 25 |
| H3K27ac (PGC2) | 29 |
| H3K27ac (Hnisz) | 23 |
| H3K4me1 (Trynka) | 25 |

**Table S2:** Overview of phenotypes and GWAS summary statistics panel. The sample size can be the total or the effective sample size. The latter is defined for for case-control studies as $N_{eff} = 4/(1/N_{cases} + 1/N_{controls})$.

| Phenotype | Abbreviation | Sample Size | Source |
|---|---|---|---|
| Age at onset of walking | AgeWalking | 70560 | 30 |
| Alcohol consumption | ALC | 112176 | 31 |
| Alcohol use disorder | AUD | 352373 | 32 |

| Phenotype | Abbreviation | Sample Size | Source |
|---|---|---|---|
| Alzheimer's disease | AD | 306866 | [33] |
| Amyotrophic lateral sclerosis | ALS | 80713 | [34] |
| Anorexia nervosa | AN | 52042 | [35] |
| Anxiety disorders | ANX | 359587 | [36] |
| Apolipoprotein B | ApoB | 88329 | [37] |
| Asthma | ASTHMA | 156857 | [38] |
| Atrial fibrillation | AF | 1840340 | [39] |
| Attention-deficit/hyperactivity disorder | ADHD | 128214 | [40] |
| Autism spectrum disorder | ASD | 53761 | [41] |
| Bipolar disorder | BIP | 220416 | [42] |
| Body mass index | BMI | 795640 | [43] |
| Bone mineral density (heel) | BMD | 426824 | [44] |
| C-reactive protein | CRP | 575531 | [45] |
| Cannabis use disorder | CUD | 375180 | [46] |
| Chronotype | CHRONOTYPE | 345148 | [38] |
| Cigarettes per day | SMOKECpD | 263956 | [47] |
| Cognitive performance | COG | 269867 | [48] |
| Coronary artery disease | CAD | 1165720 | [49] |
| Cortical surface area | SA | 32877 | [50] |
| Cortical thickness | TH | 32877 | [50] |
| COVID-19 hospitalization | COVID | 128057 | [51] |
| Crohn's disease | CD | 36151 | [52] |
| Educational attainment | EDU | 765283 | [53] |
| Estimated glomerular filtration rate | eGFRcrea | 1004040 | [54] |
| Fasting glucose | FG | 200622 | [55] |
| Fasting insulin | FI | 151013 | [55] |
| Forced expiratory volume in 1 s | FEV1 | 404165 | [56] |
| Genetic generalized epilepsy | GGE | 23363 | [57] |
| Glycated hemoglobin | HbA1c | 146806 | [55] |
| Hand grip strength (low, 60+ age) | GripStrengthAge | 256523 | [58] |
| HDL cholesterol | HDL | 1244580 | [59] |
| Heart failure | HeartFailure | 964057 | [60] |
| Height | HEIGHT | 360388 | [38] |
| Inflammatory bowel disease | IBD | 58331 | [52] |
| LDL cholesterol | LDL | 1231289 | [59] |
| Loneliness | LONELY | 262818 | [61] |
| Lymphocyte count | LYM | 524923 | [62] |
| Major depressive disorder | MDD | 1152650 | [63] |
| Mean corpuscular volume | MCV | 544127 | [62] |
| Mean platelet volume | MPV | 460935 | [62] |
| Migraine | MIG | 281344 | [64] |
| Multiple sclerosis | MS | 38093 | [65] |
| Neurocriticism | NEUR | 390278 | [66] |
| Neutrophil count | NEU | 519288 | [62] |
| Obsessive-compulsive disorder | OCD | 92032 | [67] |
| Parkinson's disease | PD | 257876 | [68] |
| Placental weight | PW | 65405 | [69] |
| Platelet count | PLT | 542827 | [62] |
| Polycystic ovary syndrome | PCOS | 22880 | [70] |

| Phenotype | Abbreviation | Sample Size | Source |
|---|---|---|---|
| Post-traumatic stress disorder | PTSD | 641533 | [71] |
| Risk tolerance | RISK | 466571 | [72] |
| Schizophrenia | SCZ | 126282 | [73] |
| Serum urate | SerumUrate | 677373 | [74] |
| Sex hormone-binding globulin | SHBG | 368929 | [75] |
| Sleep Duration | SLEEP | 384225 | [38] |
| Smoking initiation | SMOKEINIT | 632807 | [47] |
| Stroke | STROKE | 278025 | [76] |
| Suicide attempt | SUICIDE | 100907 | [77] |
| Systolic blood pressure | SBP | 1017460 | [78] |
| Telomere length | TELO | 438351 | [79] |
| Thyroid-stimulating hormone | TSH | 482863 | [80] |
| Total cholesterol | TC | 1320016 | [59] |
| Triglycerides | TG | 1253277 | [59] |
| Type 2 diabetes | T2D | 751755 | [81] |
| Ulcerative colitis | UC | 36527 | [52] |
| Waist-to-hip ratio | WHR | 697734 | [82] |
| Waist-to-hip ratio (BMI adj.) | WHRADJBMI | 694649 | [82] |
| White blood cell count | WBC | 562243 | [62] |

**Table S3:** Spearman correlation between ACS and polygenicity estimates for pruned set of phenotypes ($r_g = 0.3$).

| Functional annotation | $r_{Spear}$ | p |
|---|---|---|
| Conserved Primate, phastCons46way | 0.85 | 3e-10 |
| Functional impact score - Eigen-ALL, boosted | 0.83 | 1.7e-09 |
| Genomic Evolutionary Rate Profiling - GERP RS≥4 | 0.82 | 3.4e-09 |
| Conserved Elements, 29 Mammals, phyloP/phastCons (Lindblad-Toh) | 0.81 | 5.3e-09 |
| Zoonomia Constrained, 43 Primates, phastCons | 0.80 | 1.1e-08 |
| Combined Annotation Dependent Depletion - CADD | 0.80 | 2.1e-08 |
| Conserved Mammal, phastCons46way | 0.80 | 2.2e-08 |
| Conserved Vertebrate, phastCons46way | 0.77 | 1.3e-07 |
| Regulatory Mendelian Mutation - ReMM, boosted | 0.77 | 6.6e-08 |
| Rare Exome Variant Ensemble Learner - REVEL, boosted, 99.5 perc. | 0.75 | 4.4e-07 |
| Zoonomia Constrained, 240 Mammals, phyloP | 0.71 | 2e-06 |
| Non-coding Essential Regulation - ncER, boosted | 0.71 | 1.2e-06 |
| Regulatory Mendelian Mutation - ReMM | 0.60 | 9.6e-05 |
| Genomic Pretrained Network - GPN-MSA≤-7 | 0.48 | 0.0037 |
| Intergenic | 0 | – |
| all | 0 | – |
| Repressed (Hoffman) | 0 | – |
| Intron | -0.068 | 0.9 |
| Coding variants’ pathogenicity - PrimateAI, boosted, 99.9 perc. | -0.14 | 0.46 |
| DNase I Hypersensitivity Sites - DHS (Trynka) | -0.43 | 0.013 |
| Ancient Sequence Age Human Enhancer (Marnetto) | -0.48 | 0.005 |
| DNase I Hypersensitivity Sites peaks - DHS peaks (Trynka) | -0.49 | 0.0038 |
| Human Promoter (Villar, ExAC) | -0.50 | 0.003 |
| H3K4me1 (Trynka) | -0.52 | 0.0017 |
| Flanking Bivalent Transcription Start Site/Enhancer - BivFlnk (Roadmap) | -0.54 | 0.0014 |
| H3K9ac (Trynka) | -0.56 | 0.001 |

**Table S3**

| Functional annotation | $r_{Spear}$ | p |
|---|---|---|
| CpG Island (UCSC) | -0.56 | 0.00078 |
| H3K4me1 peaks (Trynka) | -0.57 | 0.0005 |
| Transcribed (Hoffman) | -0.58 | 0.00044 |
| H3K9ac peaks boosted (Trynka) | -0.58 | 0.00045 |
| Fetal DNase I Hypersensitivity Sites - DHS (Trynka) | -0.58 | 0.00033 |
| H3K9ac peaks (Trynka) | -0.59 | 0.00034 |
| Missense badness, PolyPhen-2, and Constraint - MPC | -0.60 | 0.00063 |
| Digital Genomic Footprinting - DGF (ENCODE) | -0.60 | 0.0002 |
| H3K27ac (Hnisz) | -0.64 | 4.8e-05 |
| Enhancer (Andersson) | -0.64 | 3.6e-05 |
| H3K9ac boosted (Trynka) | -0.65 | 4.6e-05 |
| Super Enhancer (Hnisz) | -0.65 | 3.3e-05 |
| H3K27ac (PGC2) | -0.66 | 2.8e-05 |
| H3K4me3 (Trynka) | -0.66 | 3.1e-05 |
| Synonymous nucleotide change | -0.67 | 1.3e-05 |
| 3' UTR | -0.67 | 1.9e-05 |
| H3K4me3 peaks (Trynka) | -0.67 | 1.3e-05 |
| Ancient Sequence Age Human Promoter (Marnetto) | -0.69 | 1.2e-05 |
| Transcription Factor Binding Site (ENCODE) | -0.69 | 9.8e-06 |
| 5' UTR | -0.69 | 7.1e-06 |
| Exon | -0.70 | 7.3e-06 |
| Non-synonymous nucleotide change | -0.71 | 3.5e-06 |
| Promoter Flanking (Hoffman) | -0.71 | 4.1e-06 |
| Enhancer (Hoffman) | -0.72 | 2.8e-06 |
| Weak Enhancer (Hoffman) | -0.73 | 9.8e-07 |
| Human Enhancer (Villar) | -0.73 | 1.1e-06 |
| Coding exon | -0.74 | 9.7e-07 |
| H3K4me1 QTL MaxCPP boosted (Hormozdiari) | -0.75 | 4.3e-07 |
| Human Promoter (Villar) | -0.75 | 3.7e-07 |
| Promoter (UCSC) | -0.77 | 2e-07 |
| Transcription Start Site (Hoffman) | -0.77 | 9.6e-08 |
| CTCF Binding Factor (Hoffman) | -0.79 | 6e-08 |

**Table S4:** Association between partitioned heritability and polygenicity by genomic region, from a pooled ordinary least squares model with a genomic region $\times \log_{10}(\pi)$ interaction and trait-clustered standard errors (SEs). Corresponding Spearman correlation coefficients between partitioned heritability and polygenicity are also shown.

| Genomic region | Slope | 95% CI | $p$-value | Spearman $r$ ($p$-value) |
|---|---|---|---|---|
| Exonic | −4.38 | [−6.20, −2.55] | 2.6e−6 | −0.51 ($p$ = 2.08e−3) |
| Intronic | −0.49 | [−2.49, 1.51] | 0.63 | −0.18 ($p$ = 0.30) |
| Intergenic | 4.87 | [3.06, 6.67] | 1.4e−7 | 0.59 ($p$ = 2.57e−4) |

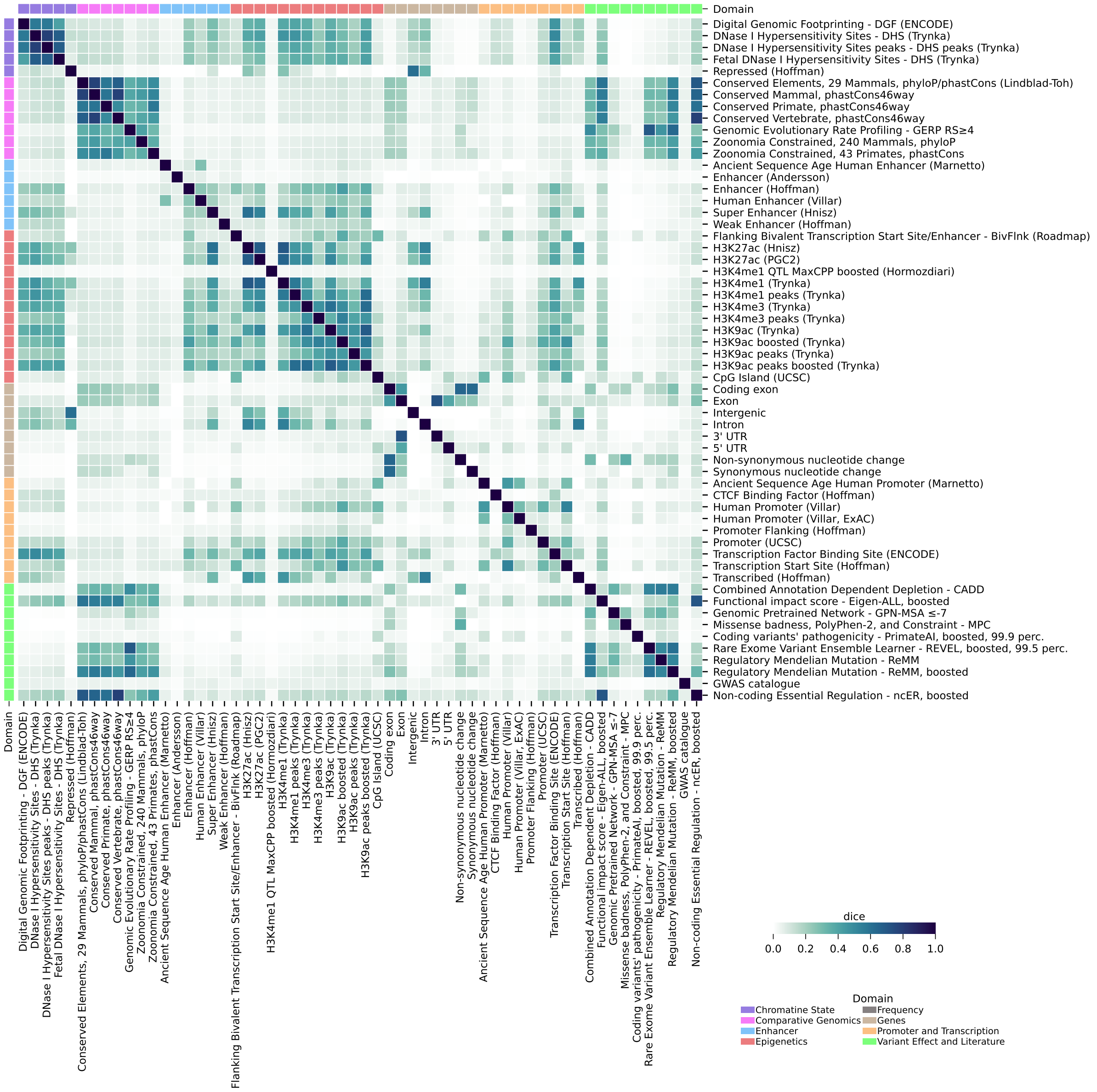


**Figure S2:** Pairwise Dice coefficient comparing the similarity between functional annotations using the 1kG template.

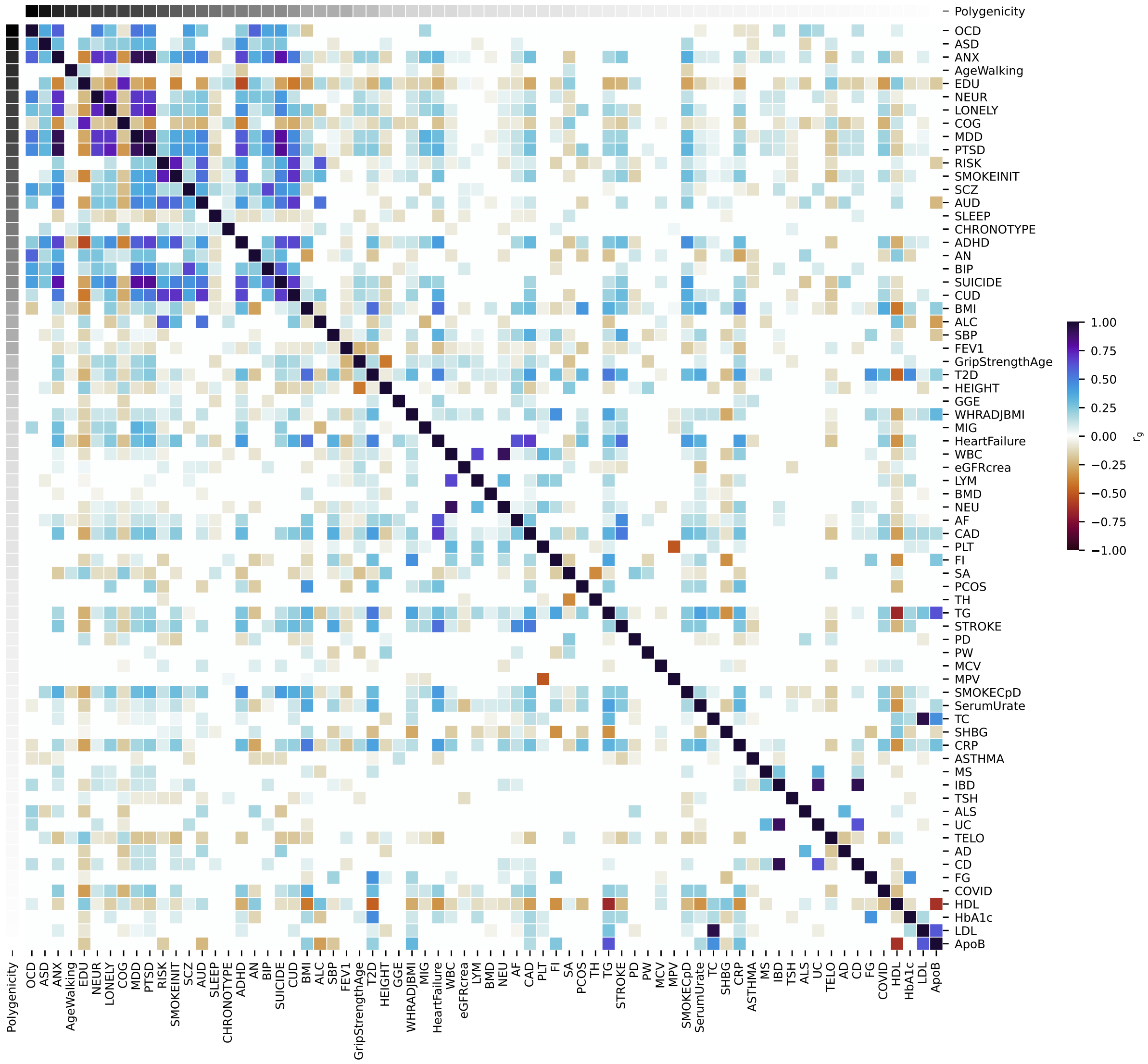


**Figure S3:** Pairwise genetic correlation coefficient $r_g$ using LDSC score regression[83] across the full phenotype panel.

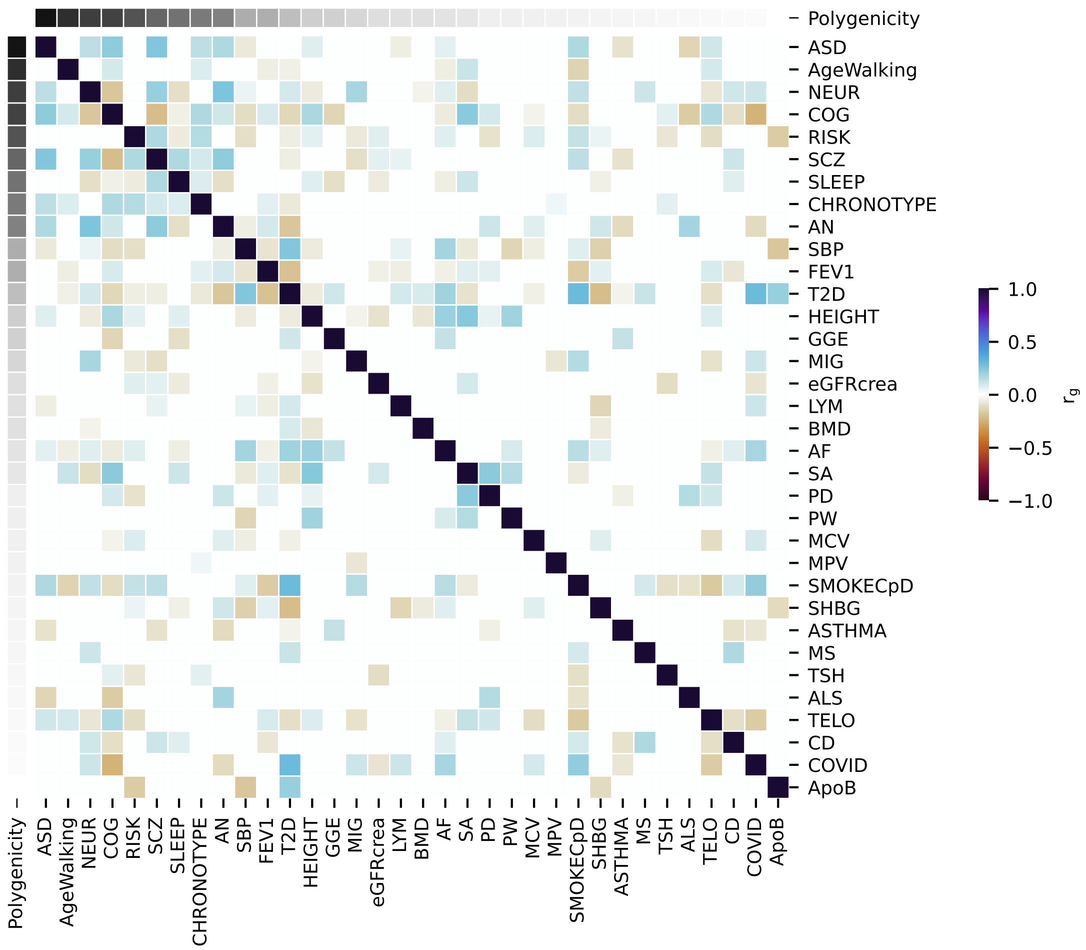


**Figure S4:** Pairwise genetic correlation coefficient $r_g$ using LDSC score regression[83] after pruning at 0.3 favoring phenotypes with higher heritability.

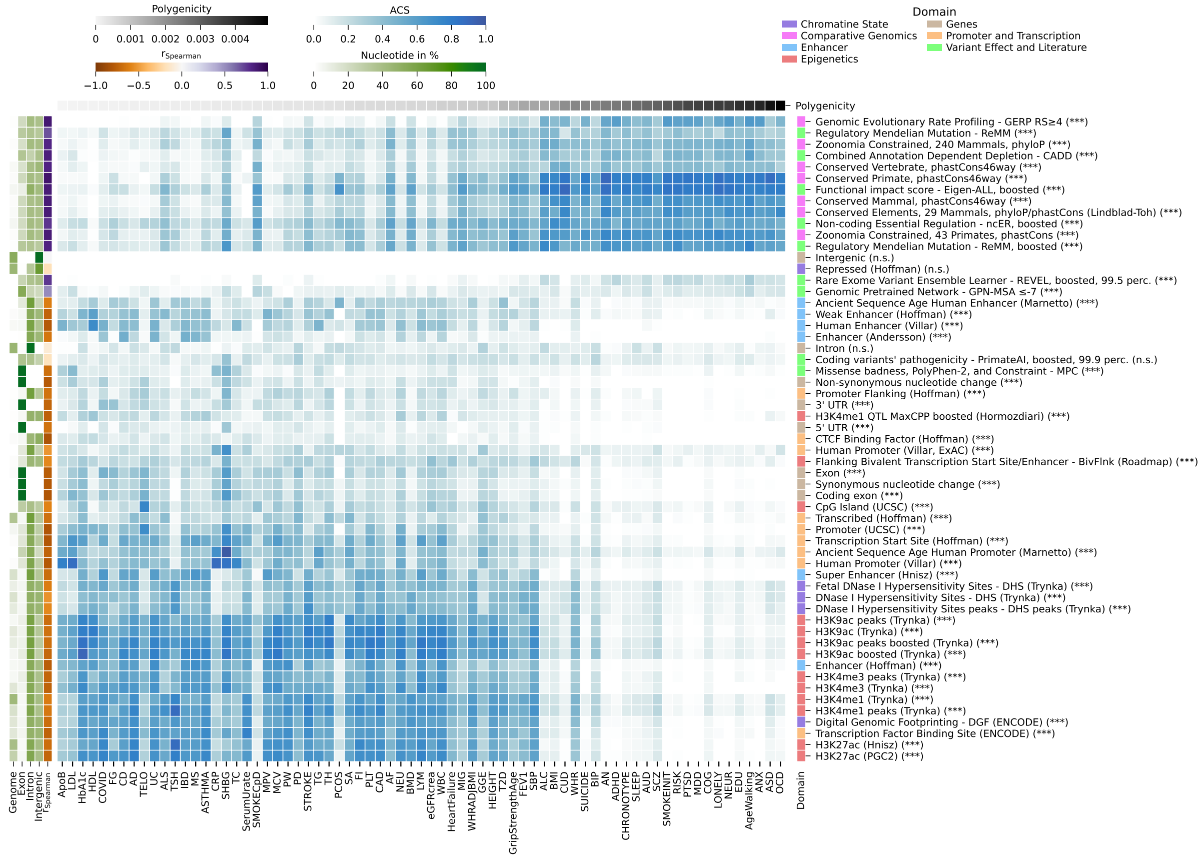


**Figure S5:** Contribution of individual functional annotations relative to the full model with 74 annotations showing all phenotypes, i.e., without pruning. Phenotypes are ordered by their polygenicity and annotations are sorted by their Euclidean distance. On the left, next to relative numbers of nucleotides (genome, exon, intron, intergenic), the Spearman rank correlation value between ACS and polygenicity across phenotypes is indicated. The symbols following each functional annotation on the right indicates the significance after Bonferroni correction for the Spearman correlation (significance: ***: p<0.001, **: p<0.01, *: p <0.05, n.s.: p≥0.05).

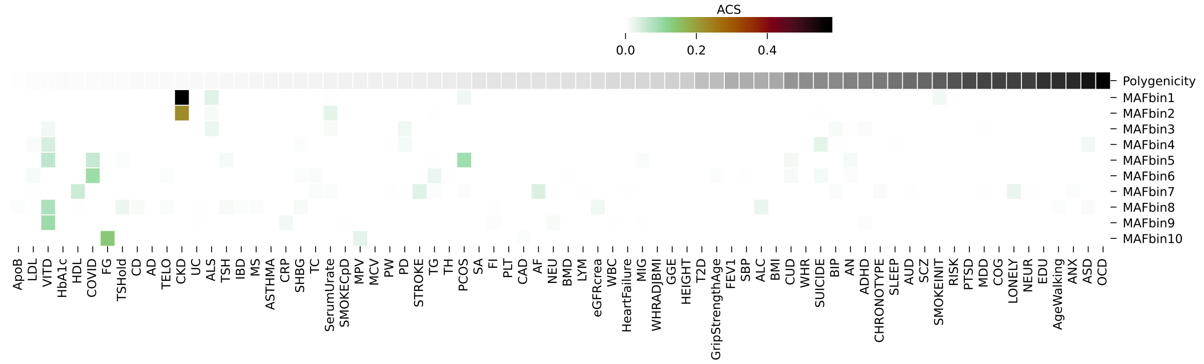


**Figure S6:** Contribution of MAF bin annotations relative to the full model with 74 annotations (phenotype pruning at rg=0.3). Phenotypes are ordered by their polygenicity. Contributions are fairly low with a max value of 0.098 for COVID.

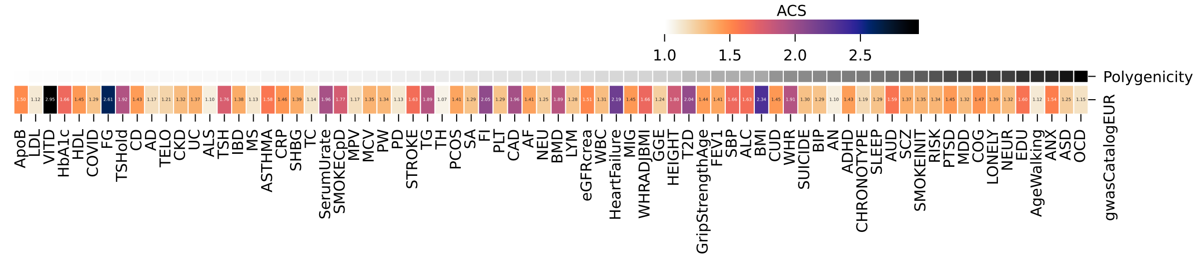


**Figure S7:** Contribution of GWAS catalouge jointly to the full model with 74 annotations across all phenotypes which are ordered by their polygenicity. For all phenotypes, its contribution is greather than one.

**(a)** Heritability estimates **(b)** Polygenicity estimates

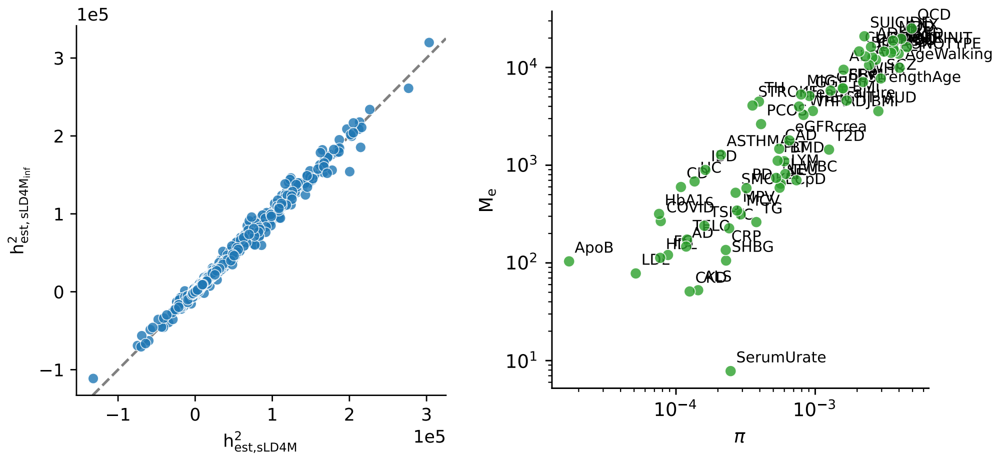


**Figure S8:** Comparison of (a) heritability fractions and (b) polygenicity estimates between sLD4M regression[1] (incorporating polygenicity estimation) and naive sLDSC style regression (SLD4M infinitesimal).

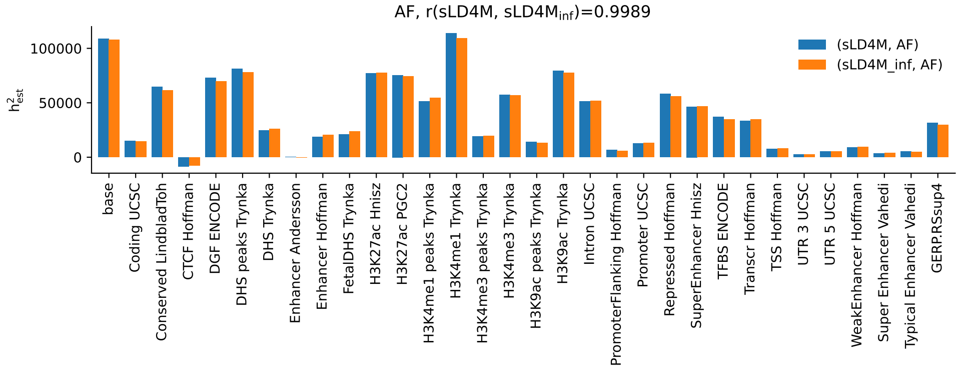


**Figure S9:** $h^2_{SNP}$ fractions from sLD4M[1] or sLDSC for pruned phenotypes. (AF)

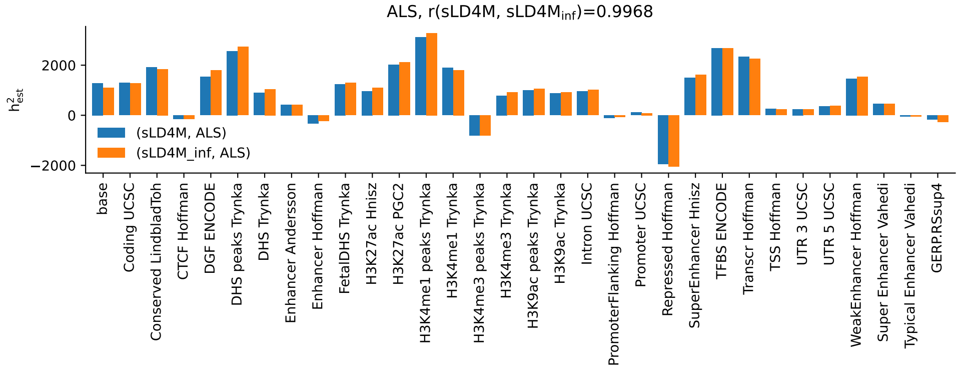


**Figure S9:** $h^2_{SNP}$ fractions from sLD4M or sLDSC for ALS

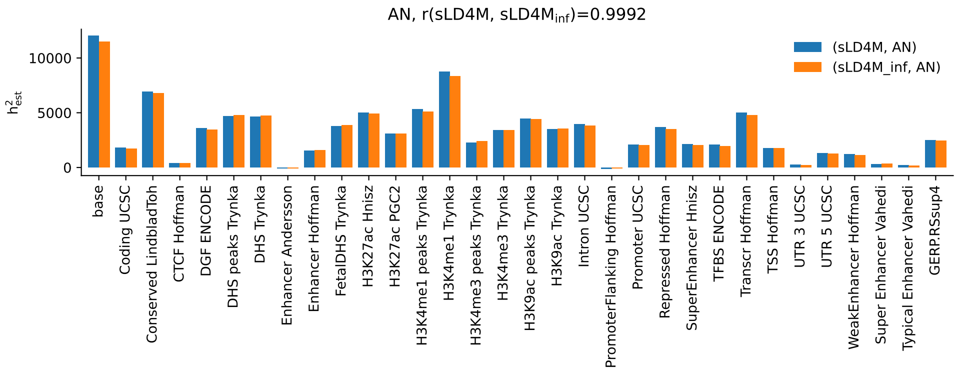


**Figure S9:** $h^2_{SNP}$ fractions from sLD4M or sLDSC for AN

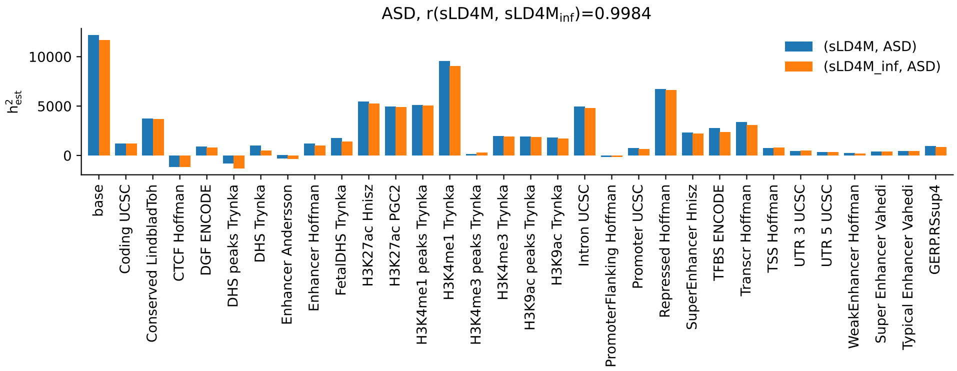


**Figure S9:** $h^2_{SNP}$ fractions from sLD4M or sLDSC for ASD

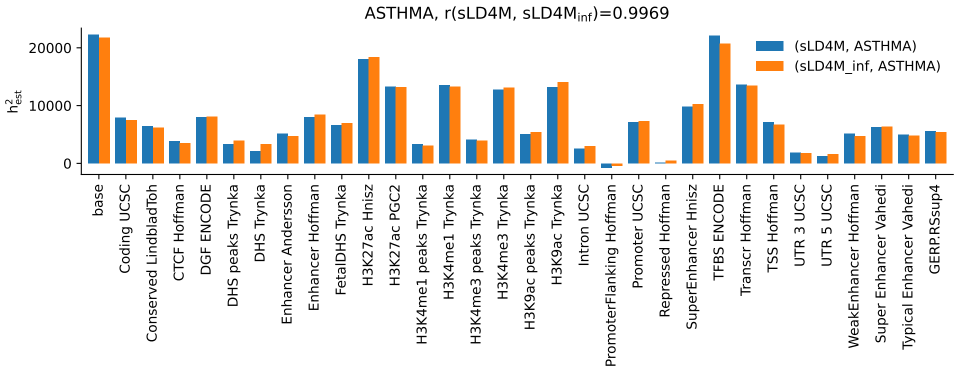


**Figure S9:** $h^2_{SNP}$ fractions from sLD4M or sLDSC for ASTHMA

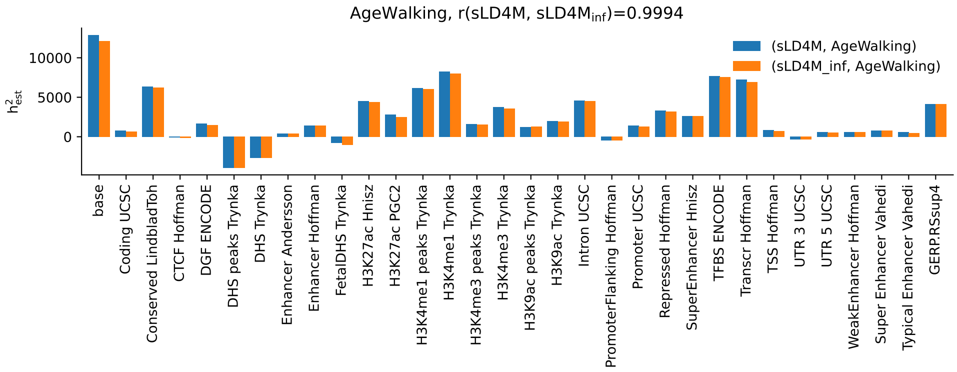


**Figure S9:** $h^2_{SNP}$ fractions from sLD4M or sLDSC for AgeWalking

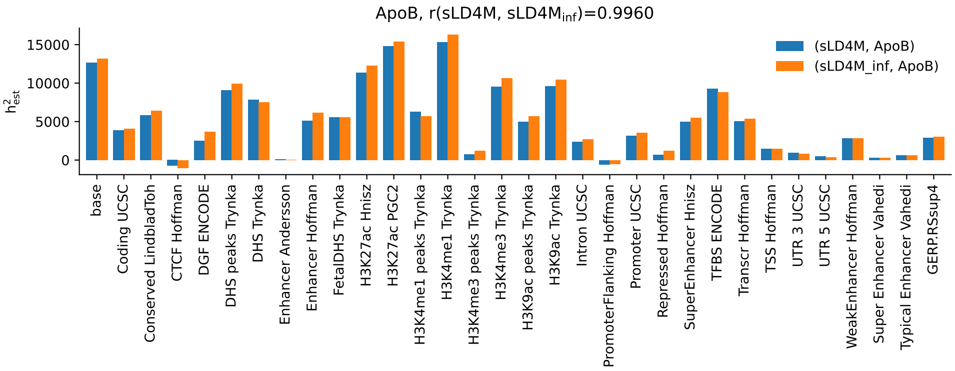


**Figure S9:** $h^2_{SNP}$ fractions from sLD4M or sLDSC for ApoB

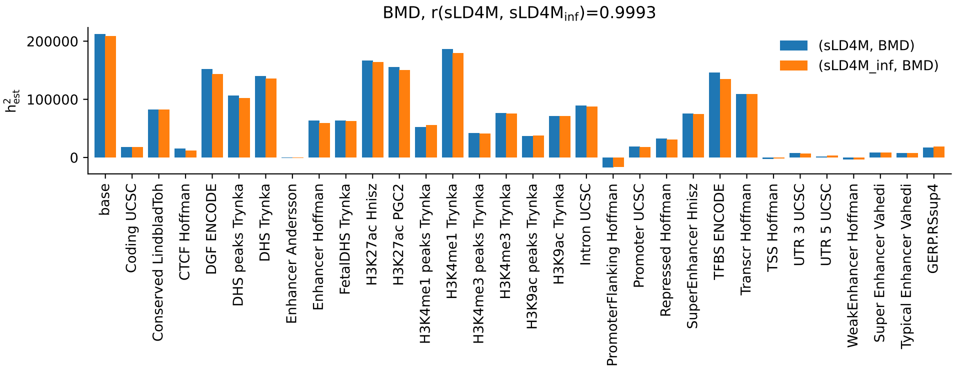


**Figure S9:** $h^2_{SNP}$ fractions from sLD4M or sLDSC for BMD

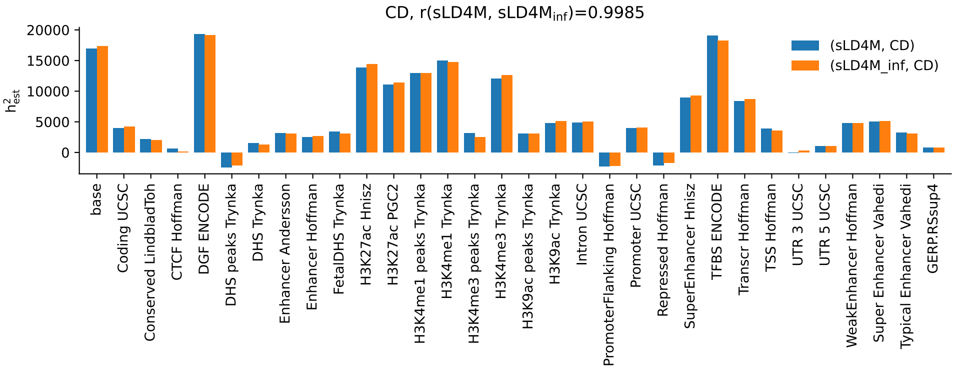


**Figure S9:** $h^2_{SNP}$ fractions from sLD4M or sLDSC for CD

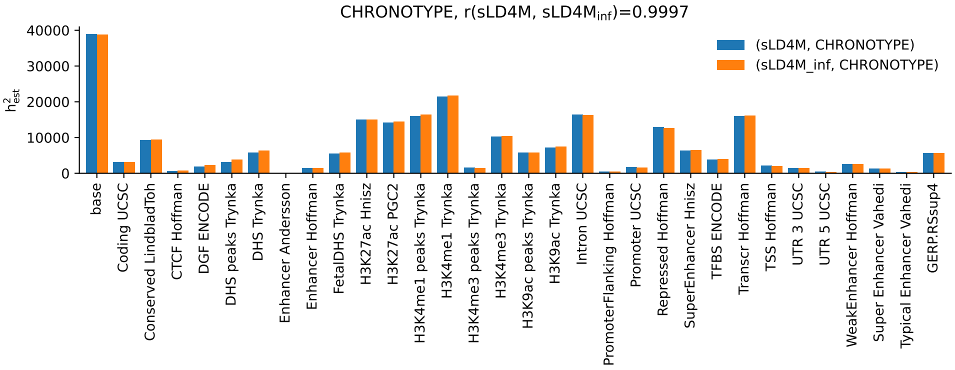


**Figure S9:** $h^2_{SNP}$ fractions from sLD4M or sLDSC for CHRONOTYPE

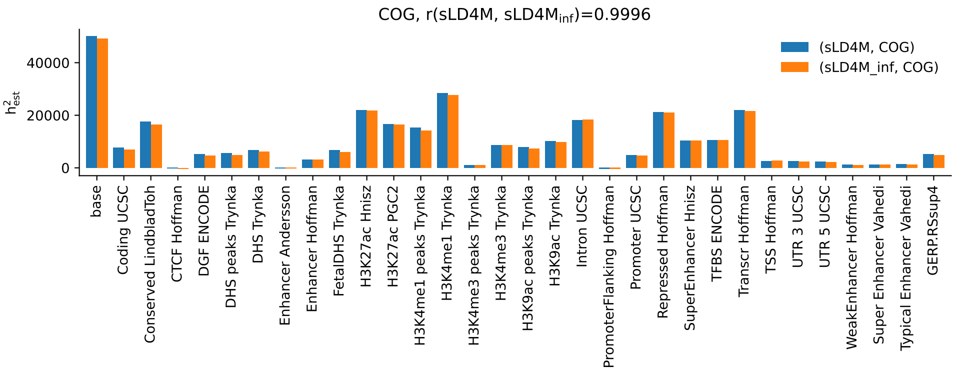


**Figure S9:** $h^2_{SNP}$ fractions from sLD4M or sLDSC for COG

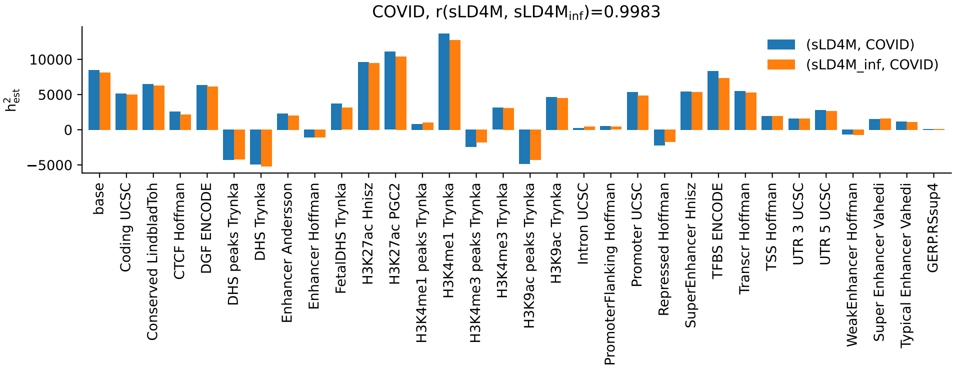


**Figure S9:** $h^2_{SNP}$ fractions from sLD4M or sLDSC for COVID

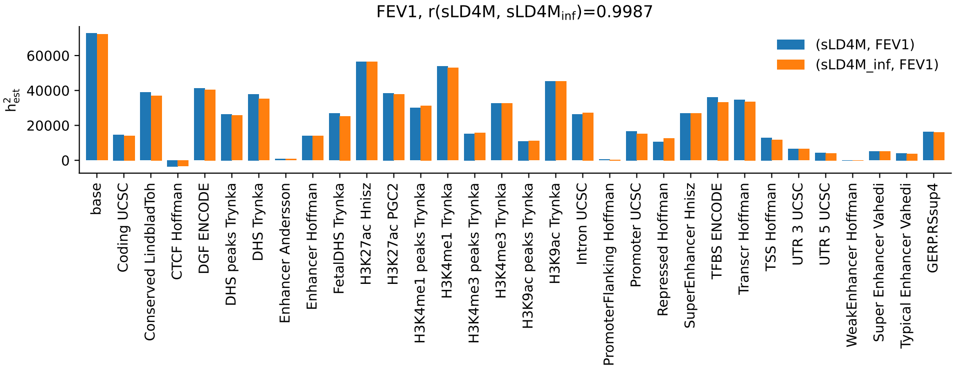


**Figure S9:** $h^2_{SNP}$ fractions from sLD4M or sLDSC for FEV1

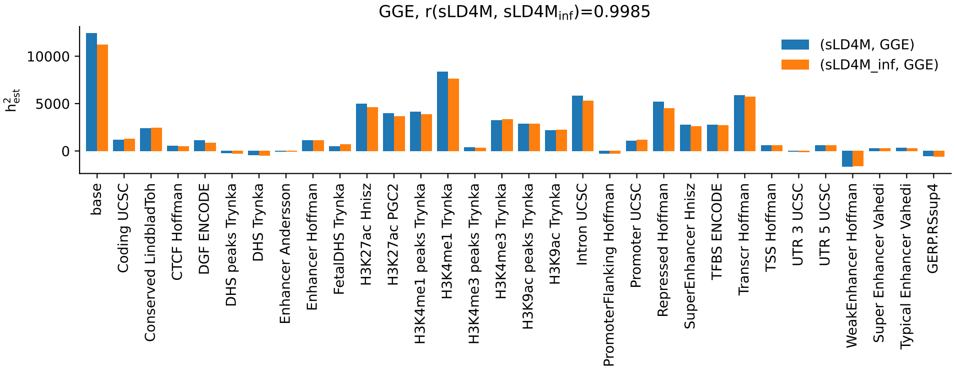


**Figure S9:** $h^2_{SNP}$ fractions from sLD4M or sLDSC for GGE

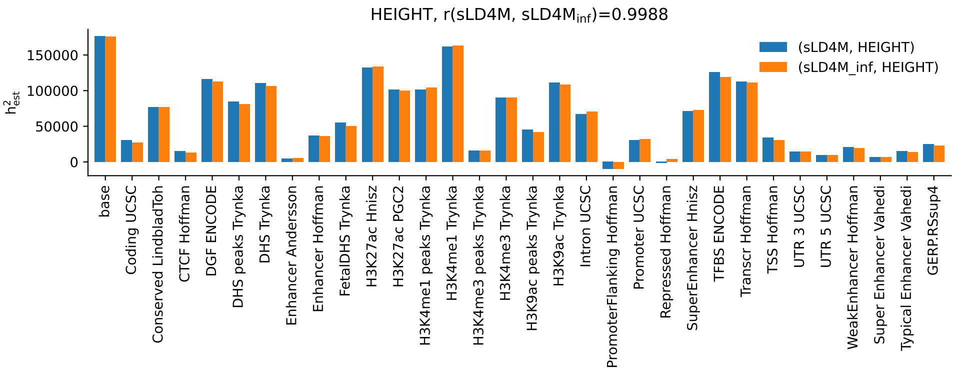


**Figure S9:** $h^2_{SNP}$ fractions from sLD4M or sLDSC for HEIGHT

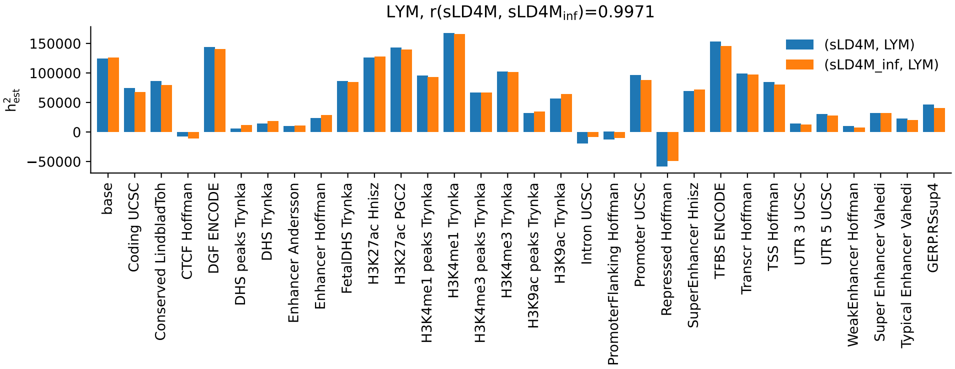


**Figure S9:** $h^2_{SNP}$ fractions from sLD4M or sLDSC for LYM

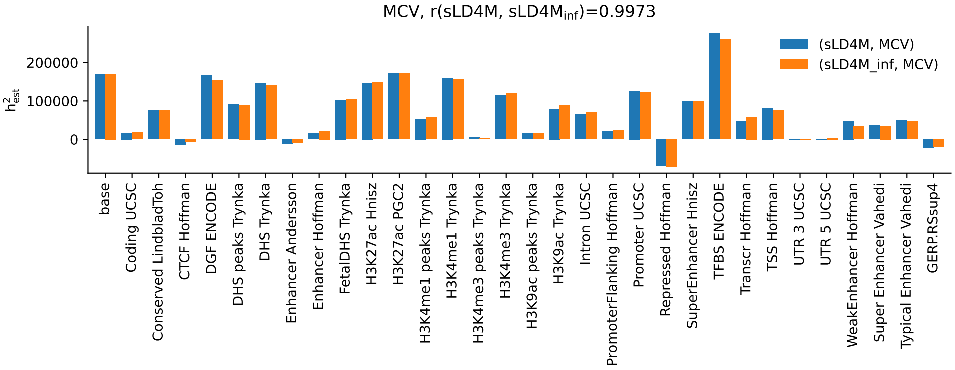


**Figure S9:** $h^2_{SNP}$ fractions from sLD4M or sLDSC for MCV

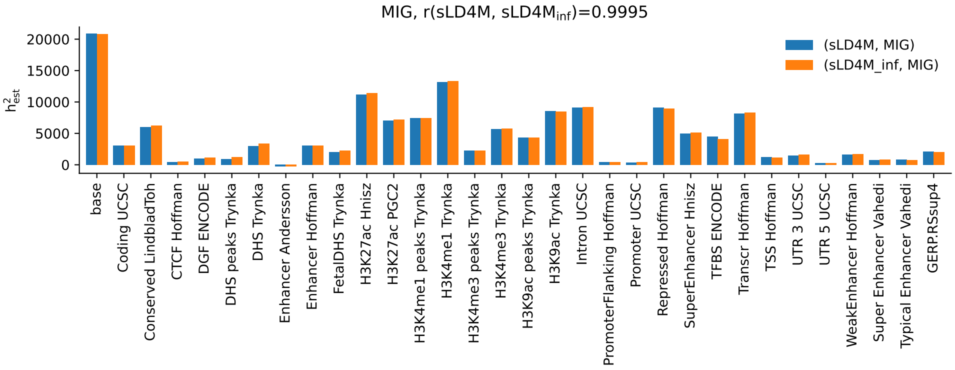


**Figure S9:** $h^2_{SNP}$ fractions from sLD4M or sLDSC for MIG

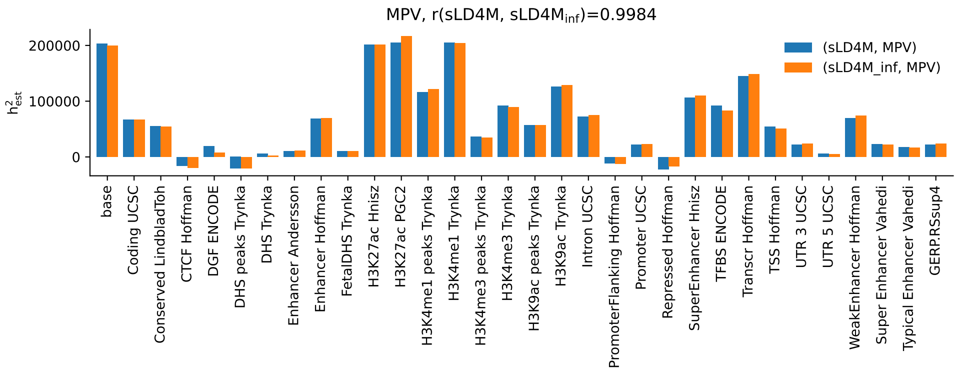


**Figure S9:** $h^2_{SNP}$ fractions from sLD4M or sLDSC for MPV

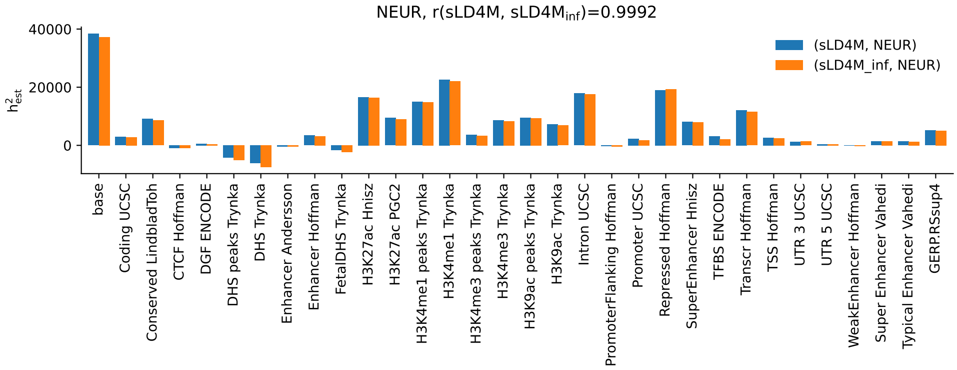


**Figure S9:** $h^2_{SNP}$ fractions from sLD4M or sLDSC for NEUR

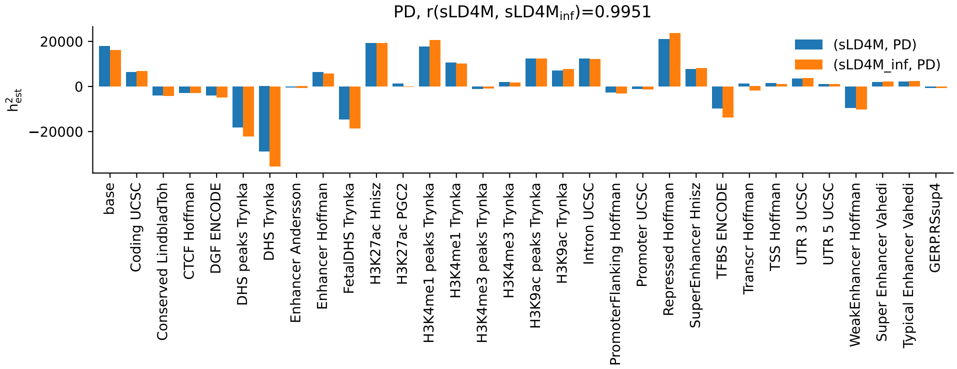


**Figure S9:** $h^2_{SNP}$ fractions from sLD4M or sLDSC for PD

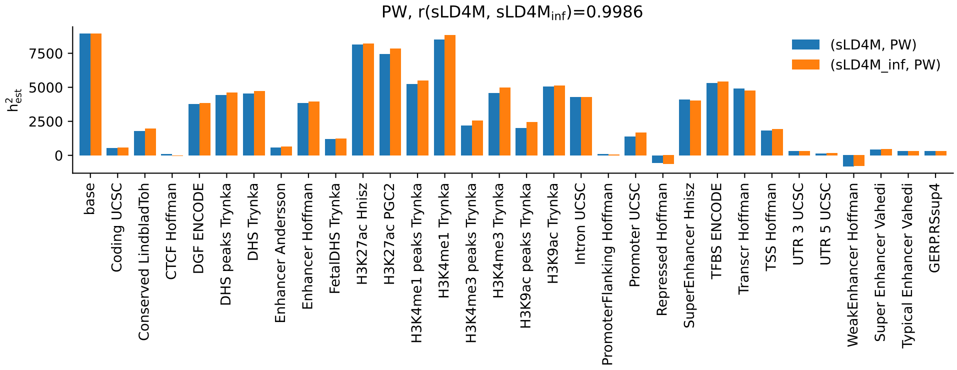


**Figure S9:** $h^2_{SNP}$ fractions from sLD4M or sLDSC for PW

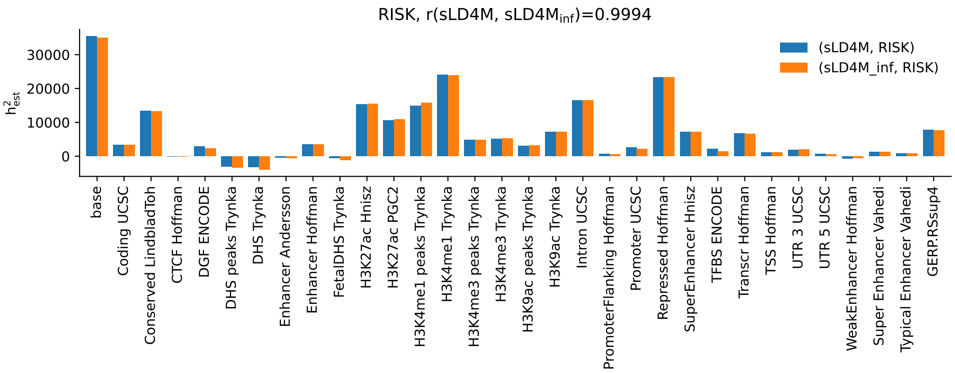


**Figure S9:** $h^2_{SNP}$ fractions from sLD4M or sLDSC for RISK

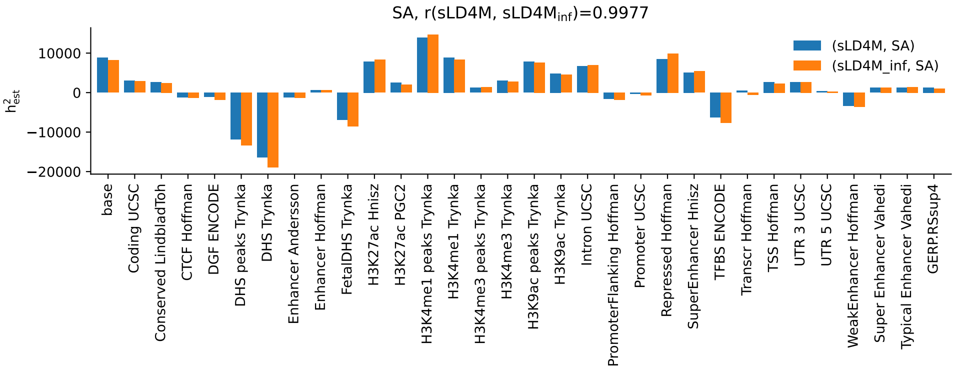


**Figure S9:** $h^2_{SNP}$ fractions from sLD4M or sLDSC for SA

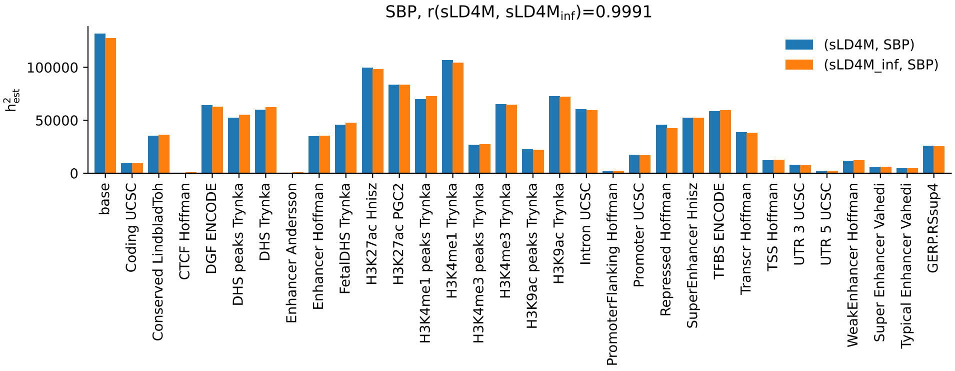


**Figure S9:** $h^2_{SNP}$ fractions from sLD4M or sLDSC for SBP

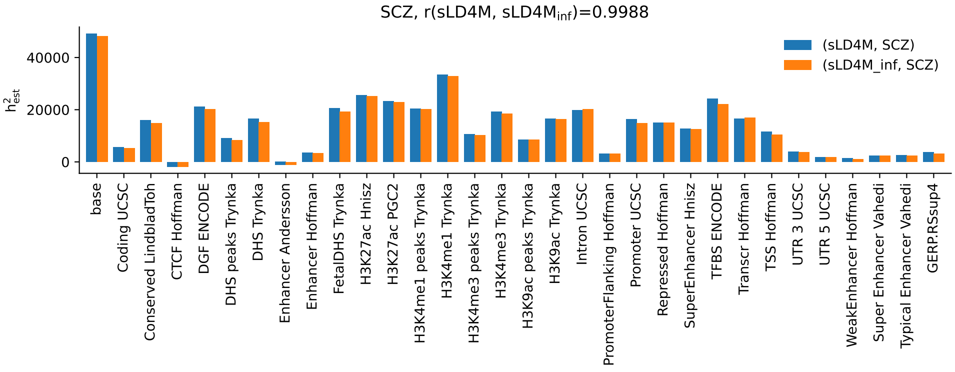


**Figure S9:** $h^2_{SNP}$ fractions from sLD4M or sLDSC for SCZ

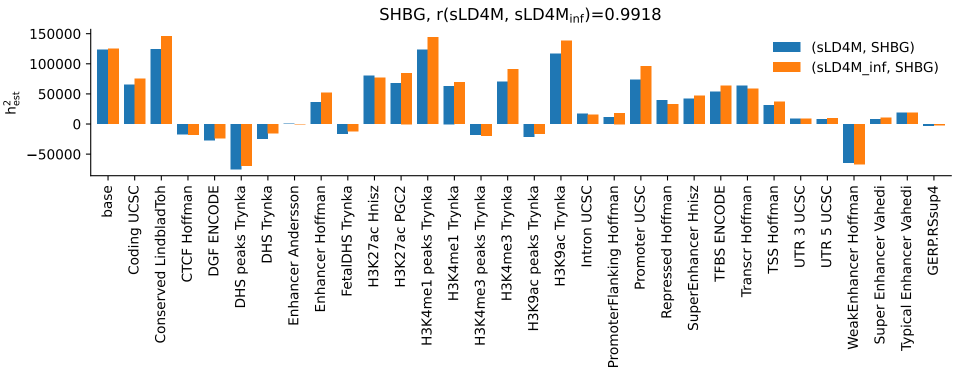


**Figure S9:** $h^2_{SNP}$ fractions from sLD4M or sLDSC for SHBG

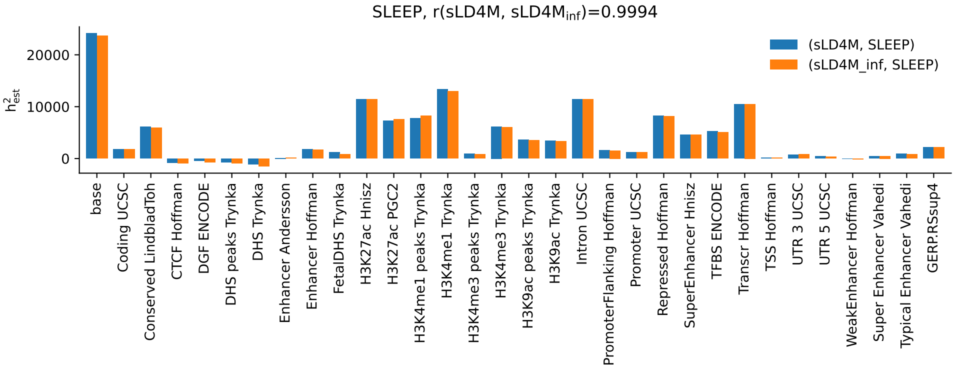


**Figure S9:** $h^2_{SNP}$ fractions from sLD4M or sLDSC for SLEEP

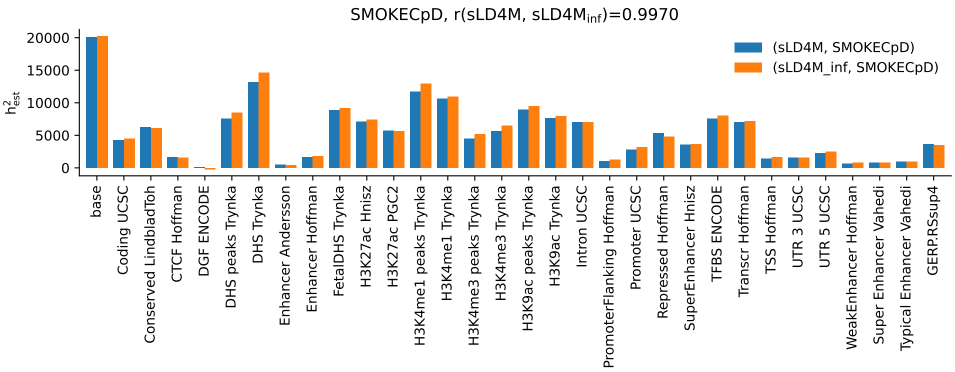


**Figure S9:** $h^2_{SNP}$ fractions from sLD4M or sLDSC for SMOKECpD

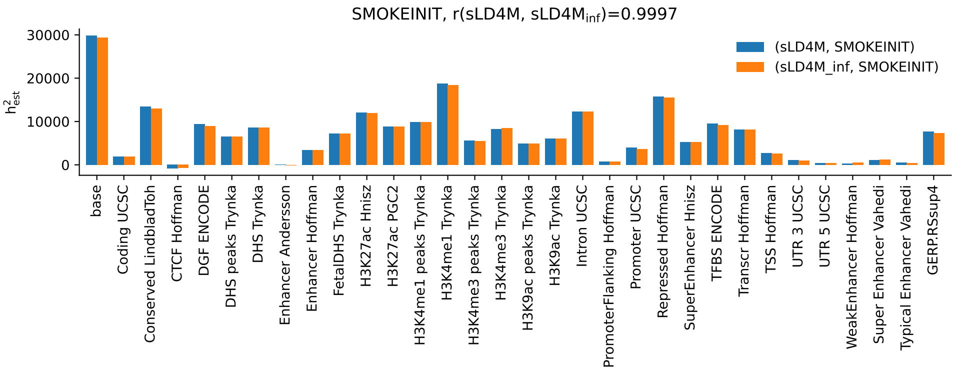


**Figure S9:** $h^2_{SNP}$ fractions from sLD4M or sLDSC for SMOKEINIT

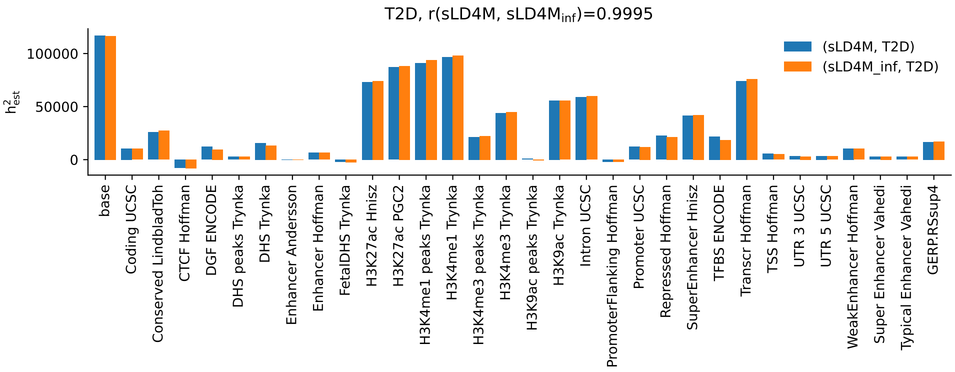


**Figure S9:** $h^2_{SNP}$ fractions from sLD4M or sLDSC for T2D

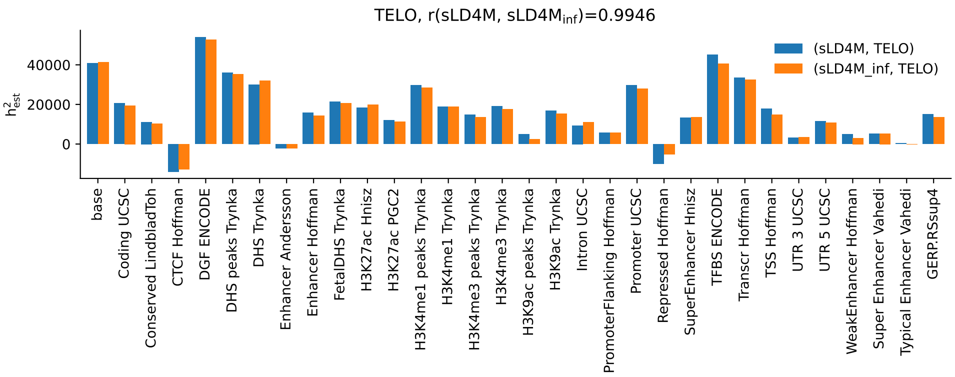


**Figure S9:** $h^2_{SNP}$ fractions from sLD4M or sLDSC for TELO

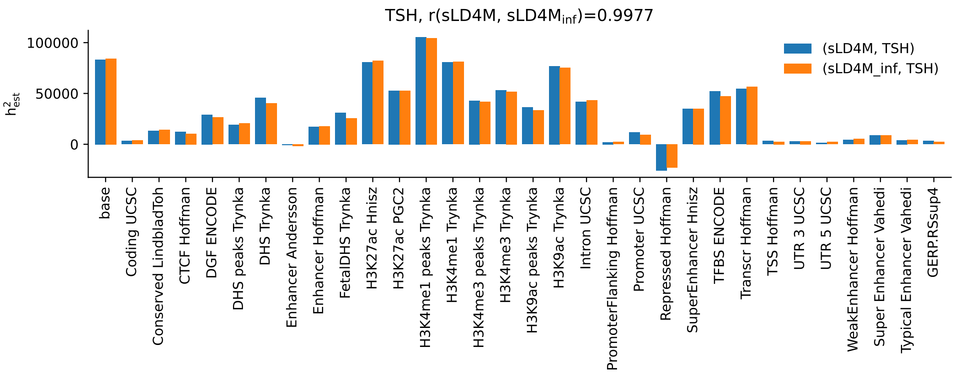


**Figure S9:** $h^2_{SNP}$ fractions from sLD4M or sLDSC for TSH

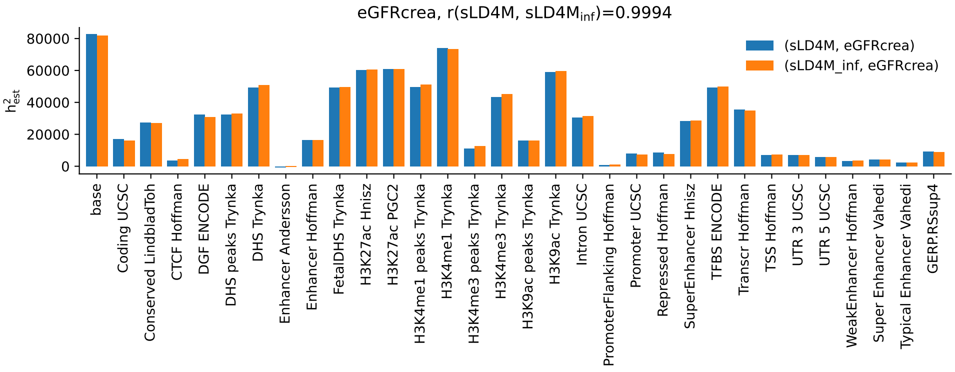


**Figure S9:** $h^2_{SNP}$ fractions from sLD4M or sLDSC for eGFRcrea

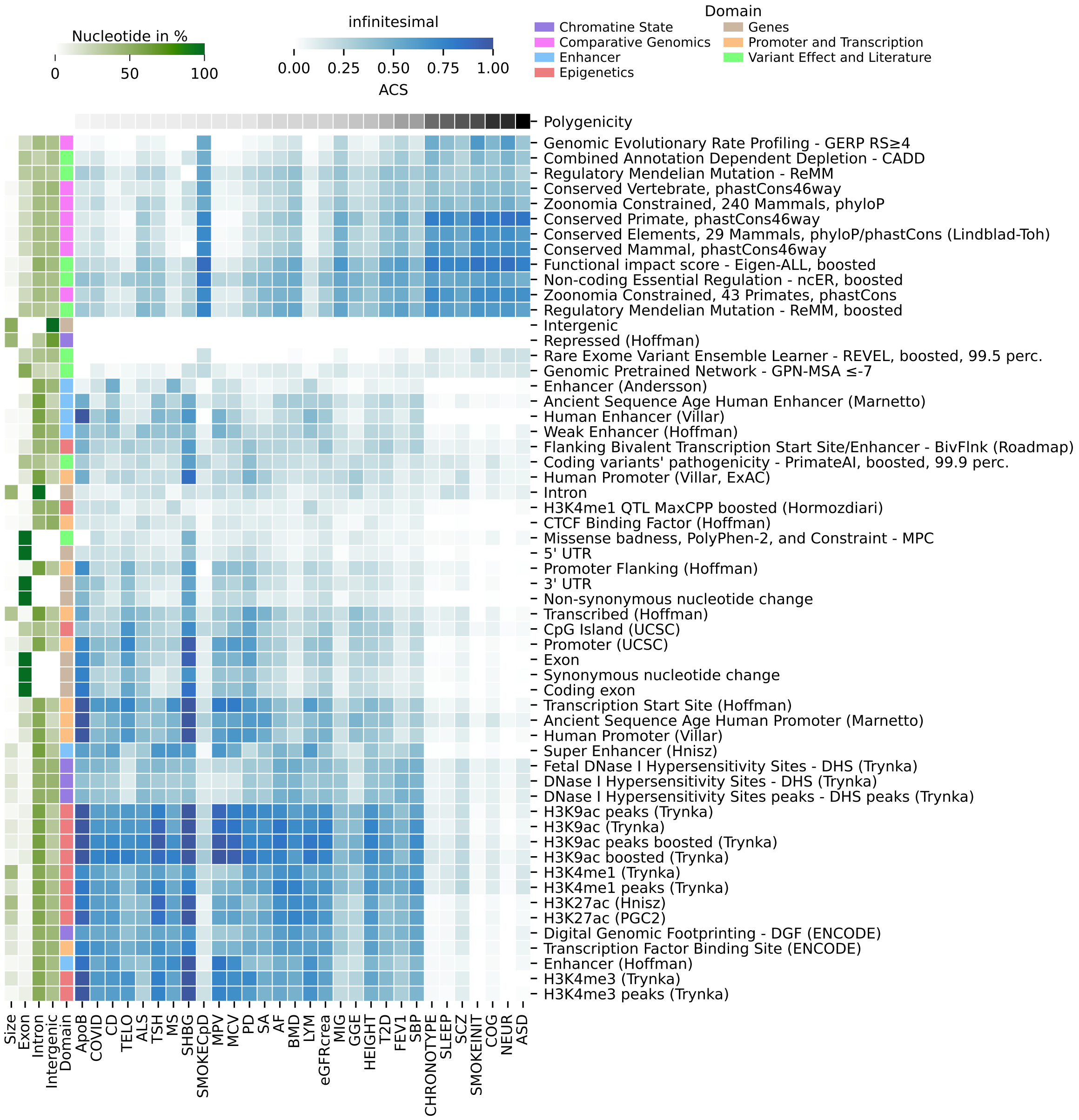


**Figure S10:** Contribution of individual functional annotations relative to the full model with 74 annotations. Phenotypes are ordered by their polygenicity remaining after pruning with $r_g$=0.3 favoring phenotypes with higher heritability. The annotation are sorted in the same way as in Fig. 2. Results show log-likelihood values based on infinitesimal assumption.

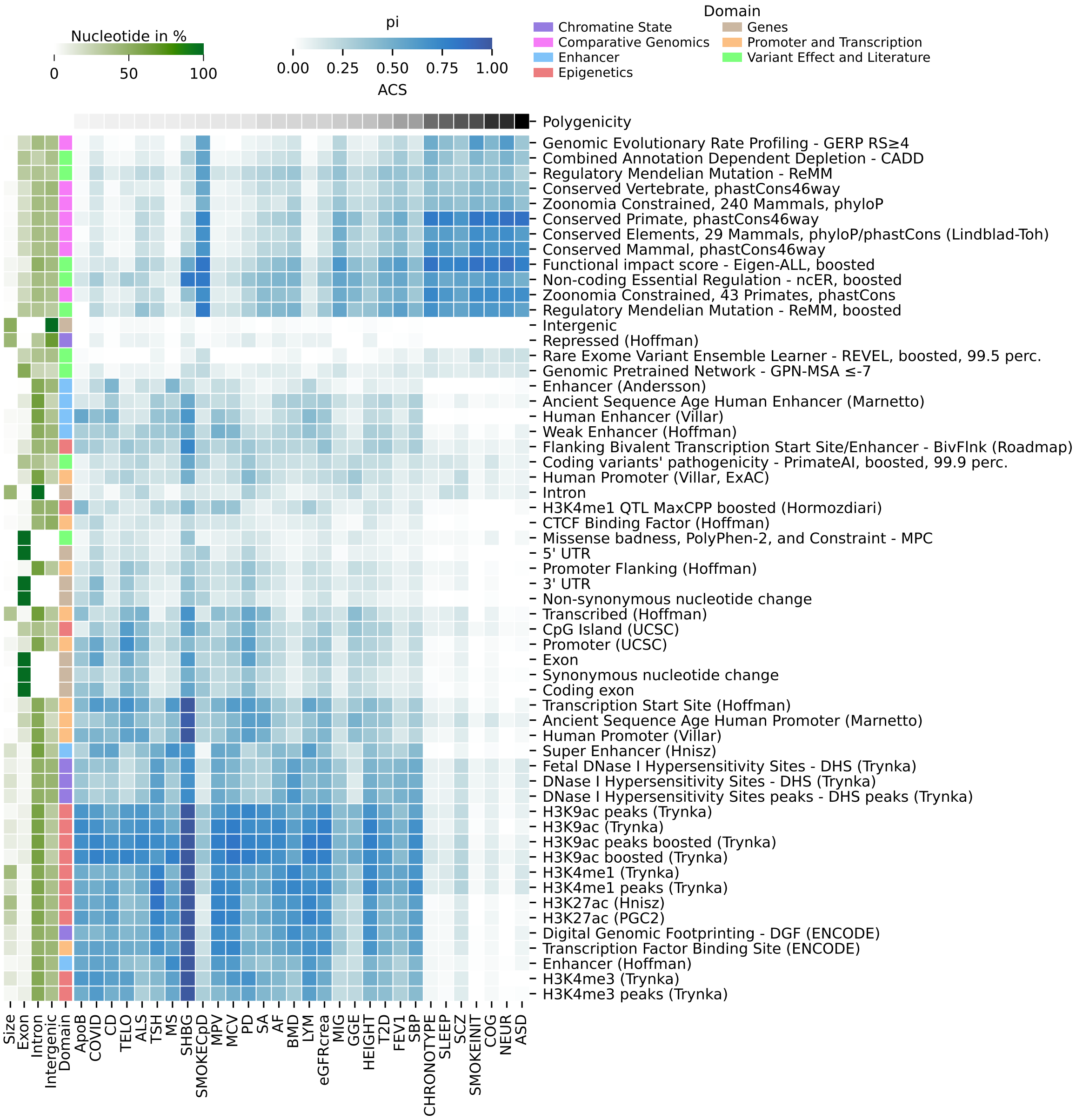


**Figure S11:** Contribution of individual functional annotations relative to the full model with 74 annotations. Phenotypes are ordered by their polygenicity remaining after pruning with $r_g$=0.3 favoring phenotypes with higher heritability. The annotation are sorted in the same way as in Fig. 2. Results show log-likelihood values based on non-infinitesimal assumption, i.e., with fitted polygenicity.

**Table S5:** Per-domain mean correlation with polygenicity. Spearman $r$ per annotation was Fisher z-transformed. Means and 95% bootstrap CIs (percentile, B=1e4) are reported for both z and r = tanh(z).

| Domain | n | $\bar{z}$ | 95% CI z | $\bar{r}$ | 95% CI r |
|---|---|---|---|---|---|
| Comparative Genomics | 7 | 1.09 | [1.01, 1.16] | 0.80 | [0.77, 0.82] |
| Variant Effect and Literature | 9 | 0.62 | [0.19, 0.97] | 0.55 | [0.18, 0.75] |
| Chromatine State | 5 | -0.47 | [-0.66, -0.23] | -0.44 | [-0.58, -0.23] |
| Genes | 8 | -0.66 | [-0.87, -0.36] | -0.58 | [-0.70, -0.35] |
| Epigenetics | 13 | -0.72 | [-0.78, -0.67] | -0.62 | [-0.65, -0.58] |
| Enhancer | 6 | -0.80 | [-0.90, -0.67] | -0.67 | [-0.72, -0.59] |
| Promoter and Transcription | 9 | -0.87 | [-0.97, -0.76] | -0.70 | [-0.75, -0.64] |

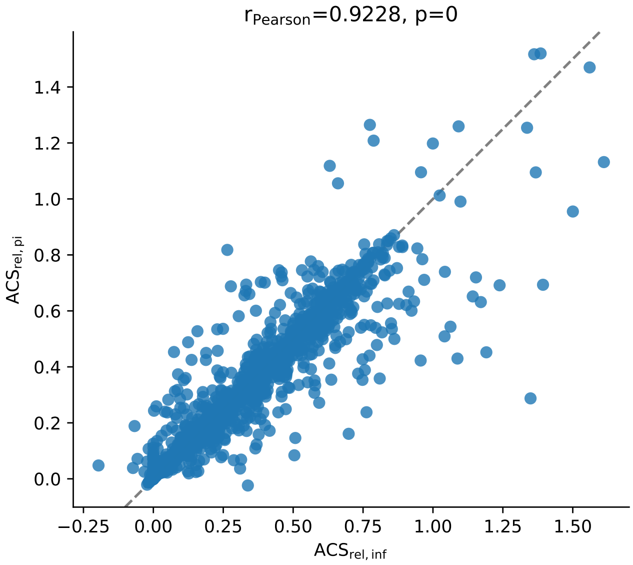


**Figure S12:** Relative log-likelihood-based ACS across pruned phenotypes and functional annotations when using an infinitesimal approach (x-axis) versus and non-infinitesimal approach (y-axis).

**Table S6:** Pairwise differences in mean correlation with polygenicity across functional domains. For each domain, r denotes the Spearman correlation between ACS and phenotype polygenicity (Fisher z-transformed and averaged across traits). $\Delta r = \tanh(\bar{z}_A) - \tanh(\bar{z}_B)$ gives the difference in mean correlation (Domain A − Domain B). CIs are 95% bootstrap (percentile, $B = 10^4$). $p$-values are obtained from 1e5 label permutations. An asterisk indicates that the comparison remains significant after Benjamini–Hochberg FDR correction at $q < 0.05$. Positive $\Delta r$ indicates Domain A > Domain B in terms of mean correlation with polygenicity.

| Domain A | Domain B | Δr [95% CI] | p |
|---|---|---|---|
| Epigenetics | Comparative Genomics | -1.41 [-1.46, -1.37] | 3.0e-05* |
| Epigenetics | Variant Effect and Literature | -1.17 [-1.37, -0.81] | 2.0e-05* |
| Promoter and Transcription | Comparative Genomics | -1.50 [-1.56, -1.43] | 1.2e-04* |
| Promoter and Transcription | Variant Effect and Literature | -1.25 [-1.46, -0.88] | 9.0e-05* |
| Genes | Comparative Genomics | -1.37 [-1.51, -1.15] | 1.4e-04* |
| Enhancer | Comparative Genomics | -1.46 [-1.52, -1.38] | 5.4e-04* |
| Genes | Variant Effect and Literature | -1.12 [-1.38, -0.71] | 5.6e-04* |
| Enhancer | Variant Effect and Literature | -1.21 [-1.42, -0.84] | 0.00125* |
| Chromatine State | Comparative Genomics | -1.24 [-1.38, -1.02] | 0.00141* |
| Chromatine State | Promoter and Transcription | 0.26 [0.11, 0.48] | 0.0153* |
| Chromatine State | Epigenetics | 0.17 [0.04, 0.39] | 0.00436* |
| Chromatine State | Variant Effect and Literature | -0.99 [-1.25, -0.58] | 0.00736* |
| Epigenetics | Promoter and Transcription | 0.09 [0.02, 0.15] | 0.179 |
| Chromatine State | Enhancer | 0.22 [0.07, 0.45] | 0.0108* |
| Variant Effect and Literature | Comparative Genomics | -0.25 [-0.60, -0.05] | 0.0236* |
| Genes | Promoter and Transcription | 0.13 [-0.02, 0.33] | 0.309 |
| Epigenetics | Enhancer | 0.05 [-0.03, 0.12] | 0.122 |
| Genes | Chromatine State | -0.13 [-0.39, 0.12] | 0.406 |
| Genes | Enhancer | 0.09 [-0.06, 0.30] | 0.385 |
| Promoter and Transcription | Enhancer | -0.04 [-0.13, 0.05] | 0.947 |
| Genes | Epigenetics | 0.04 [-0.09, 0.26] | 0.554 |

**Table S7:** $90^{th}$-percentile of $\log L_{rel}$ across annotation-specific relative nucleotide count bins. Bins were chosen to align with nucleotide distributions and maintain counts per bin, showing highest number in mid-size annotations.

| Bin | Size bin (%) | Q90 |
|---|---|---|
| 4 | (10, 15] | 0.68 |
| 5 | (15, 25] | 0.64 |
| 2 | (3, 6] | 0.63 |
| 6 | (25, ∞) | 0.60 |
| 1 | (1, 3] | 0.54 |
| 3 | (6, 10] | 0.50 |
| 0 | (0, 1] | 0.38 |

**Table S8:** MAF deciles used in the analysis based on 1000 Genome reference panel.

| MAF bin | Interval |
|---|---|
| 1 | [0.0051, 0.0072] |
| 2 | (0.0072, 0.0133] |
| 3 | (0.0133, 0.0256] |
| 4 | (0.0256, 0.0481] |
| 5 | (0.0481, 0.0869] |
| 6 | (0.0869, 0.1421] |
| 7 | (0.1421, 0.2137] |
| 8 | (0.2137, 0.2996] |
| 9 | (0.2996, 0.3967] |
| 10 | (0.3967, 0.5000] |

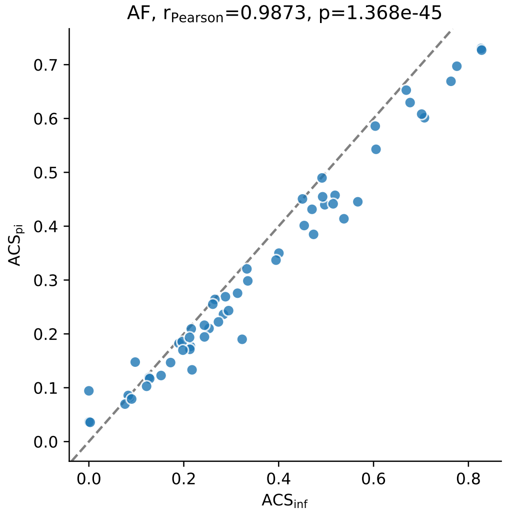


**Figure S13:** Relative log-likelihood-based ACS across individual pruned phenotypes and functional annotations when using an infinitesimal approach versus and non-infinitesimal approach (AF)

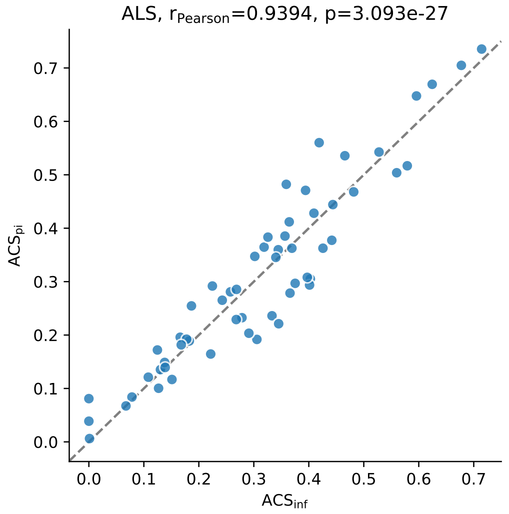


**Figure S13:** ACS comparison for ALS

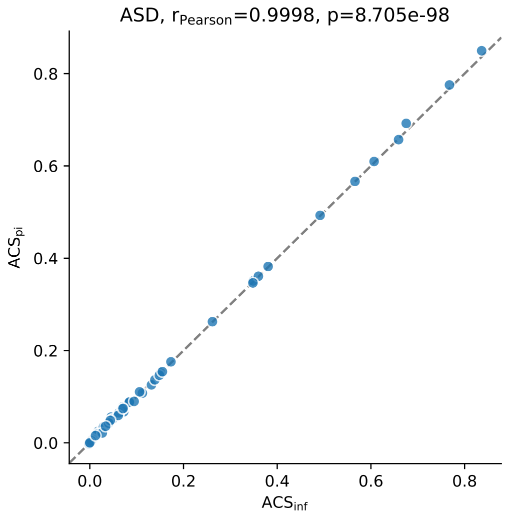


**Figure S13:** ACS comparison for ASD

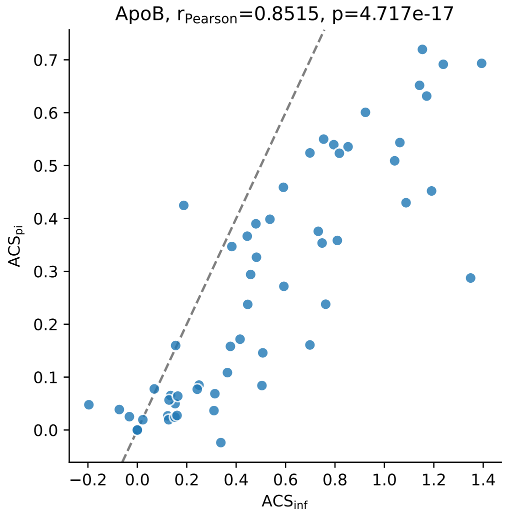


**Figure S13:** ACS comparison for ApoB

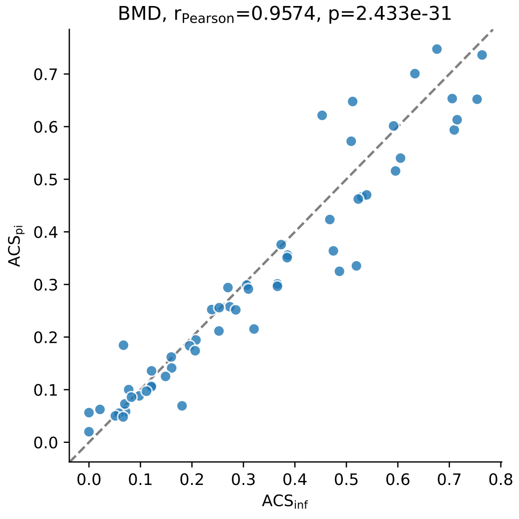


**Figure S13:** ACS comparison for BMD

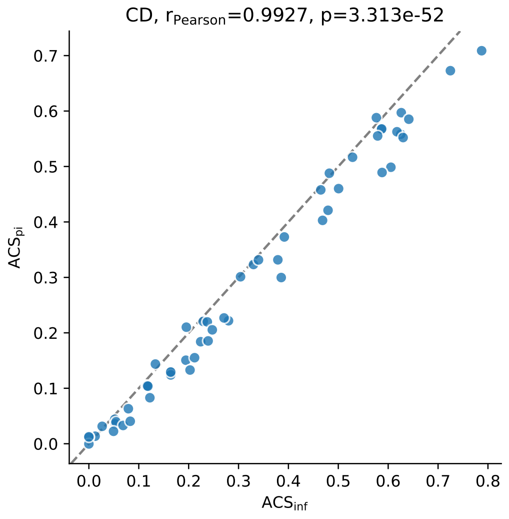


**Figure S13:** ACS comparison for CD

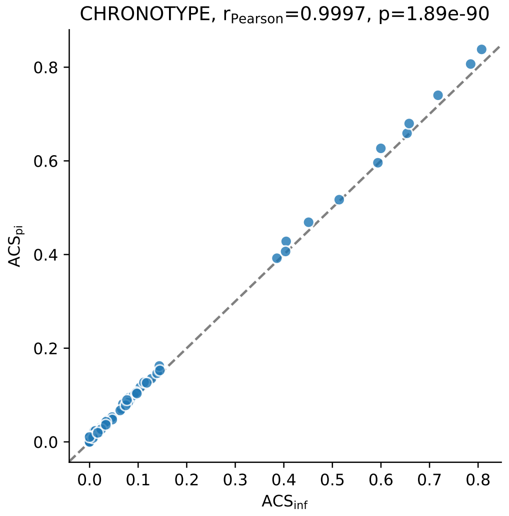


**Figure S13:** ACS comparison for CHRONOTYPE

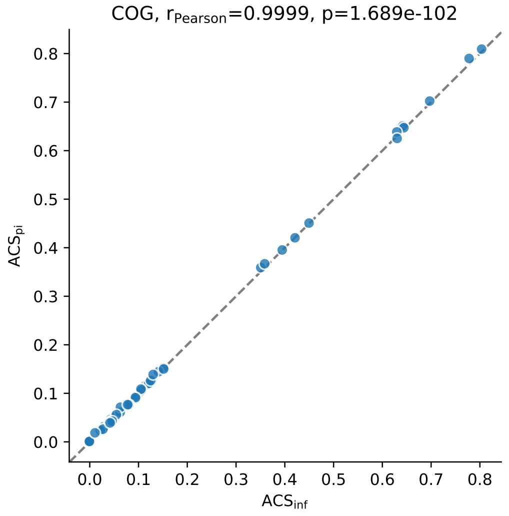


**Figure S13:** ACS comparison for COG

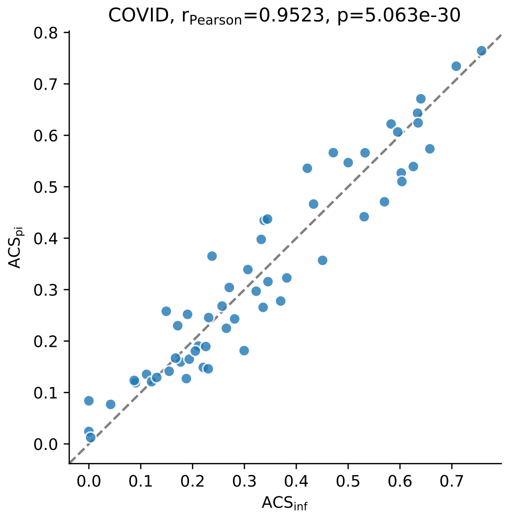


**Figure S13:** ACS comparison for COVID

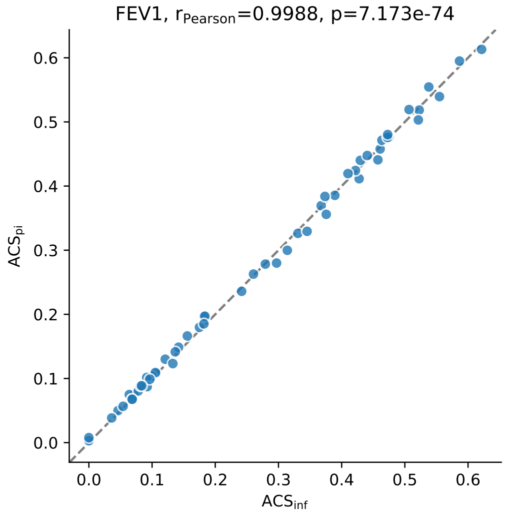


**Figure S13:** ACS comparison for FEV1

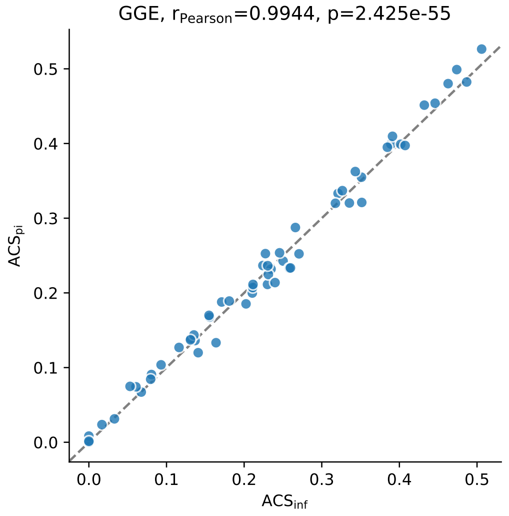


**Figure S13:** ACS comparison for GGE

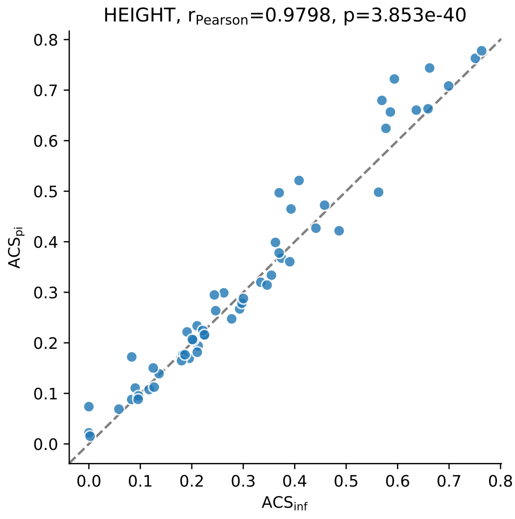


**Figure S13:** ACS comparison for HEIGHT

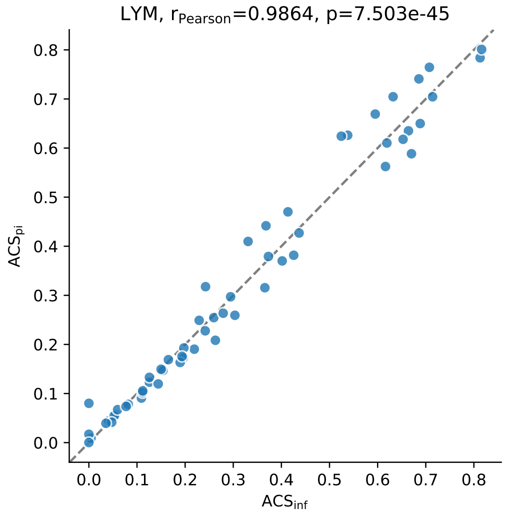


**Figure S13:** ACS comparison for LYM

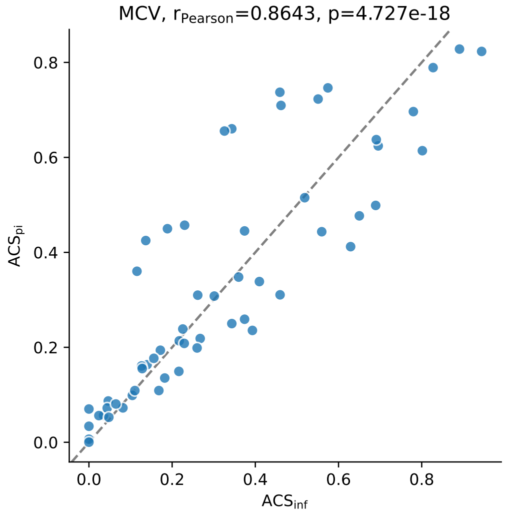


**Figure S13:** ACS comparison for MCV

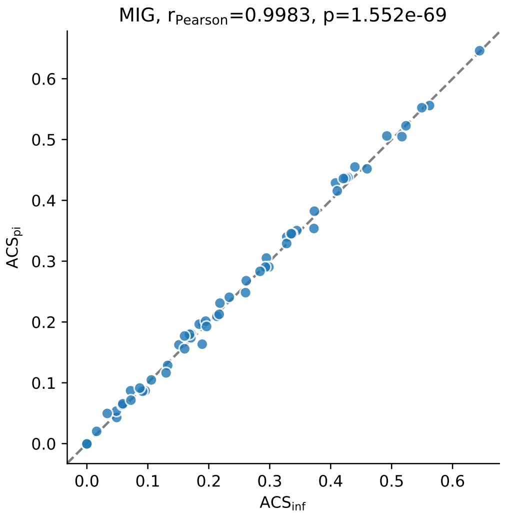


**Figure S13:** ACS comparison for MIG

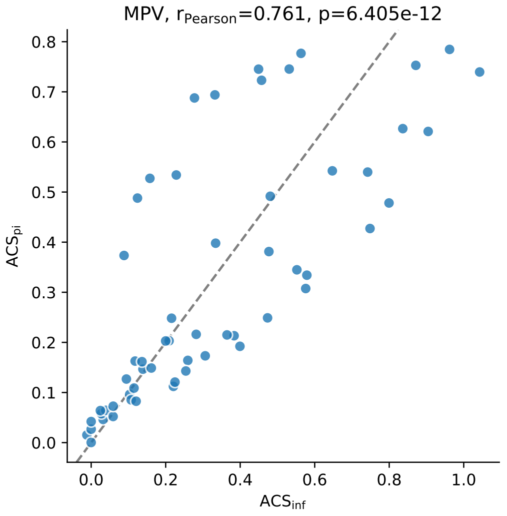


**Figure S13:** ACS comparison for MPV

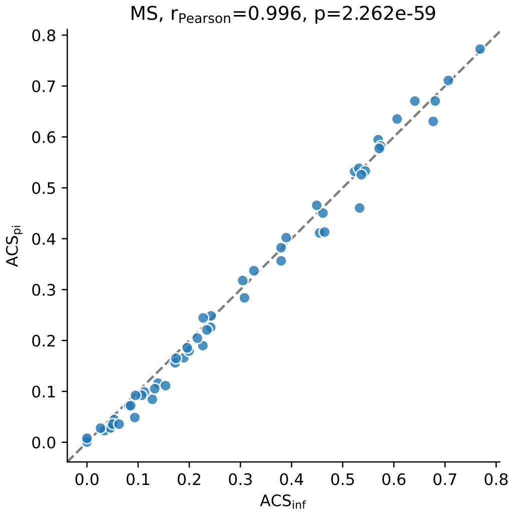


**Figure S13:** ACS comparison for MS

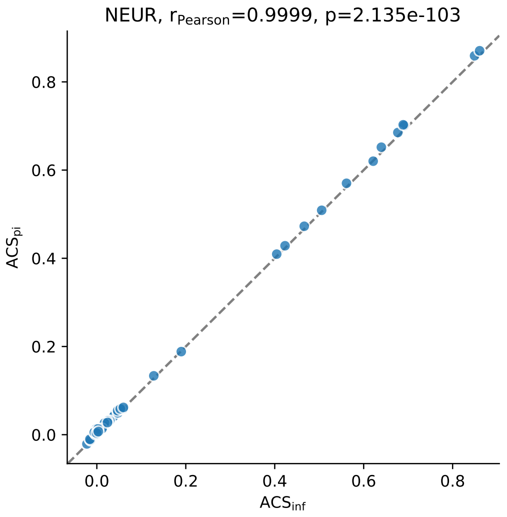


**Figure S13:** ACS comparison for NEUR

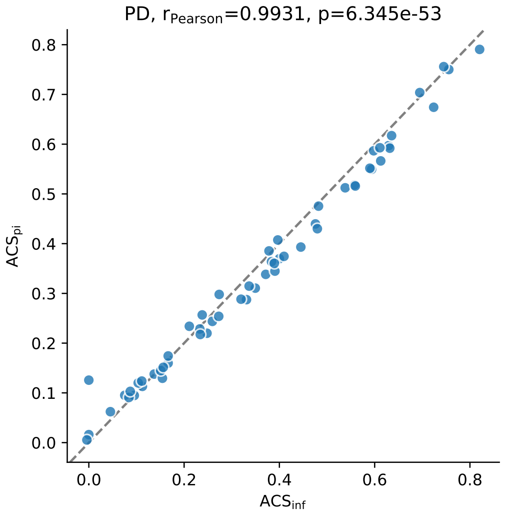


**Figure S13:** ACS comparison for PD

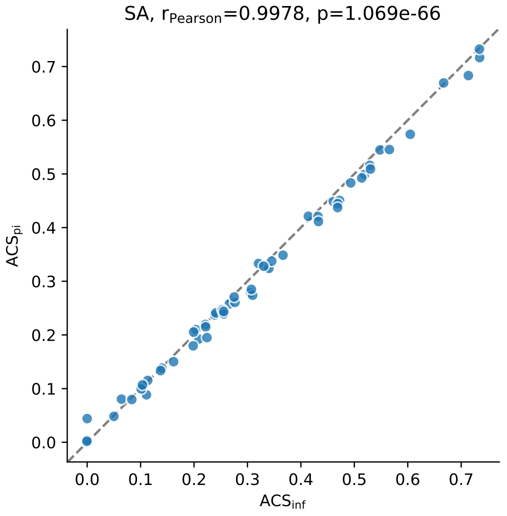


**Figure S13:** ACS comparison for SA

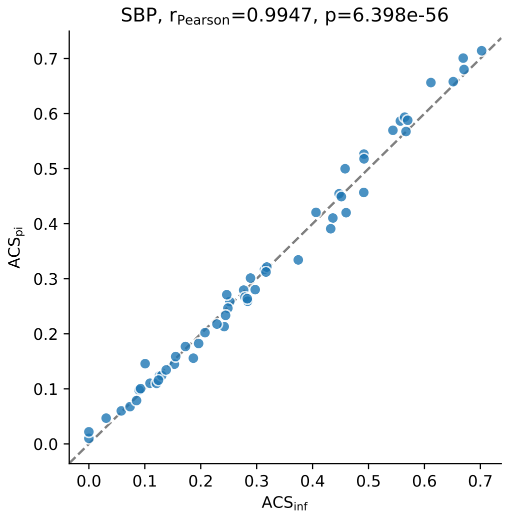


**Figure S13:** ACS comparison for SBP

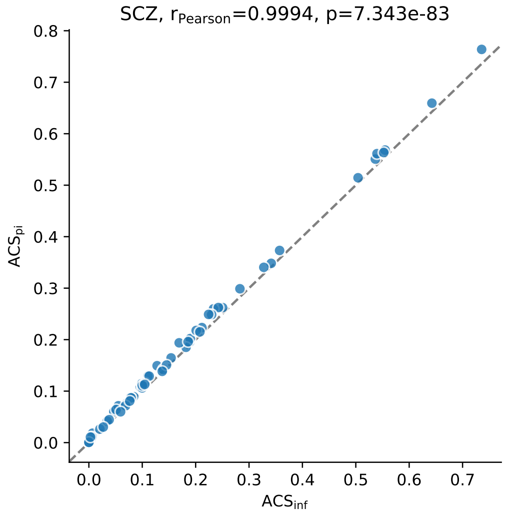


**Figure S13:** ACS comparison for SCZ

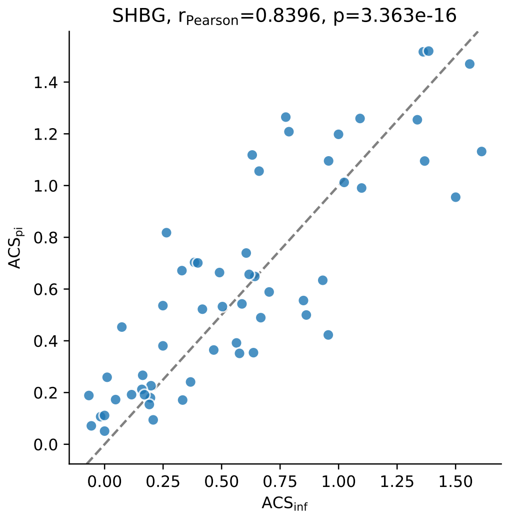


**Figure S13:** ACS comparison for SHBG

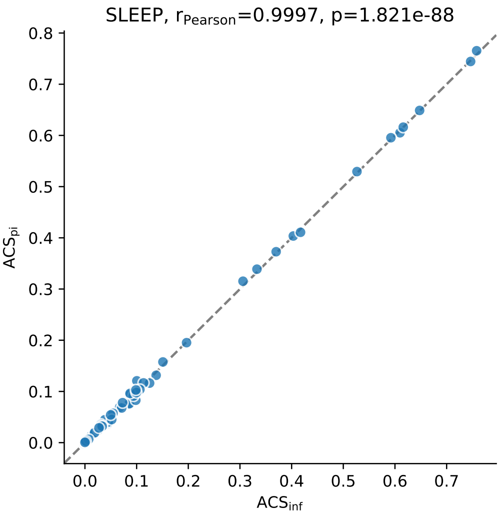


**Figure S13:** ACS comparison for SLEEP

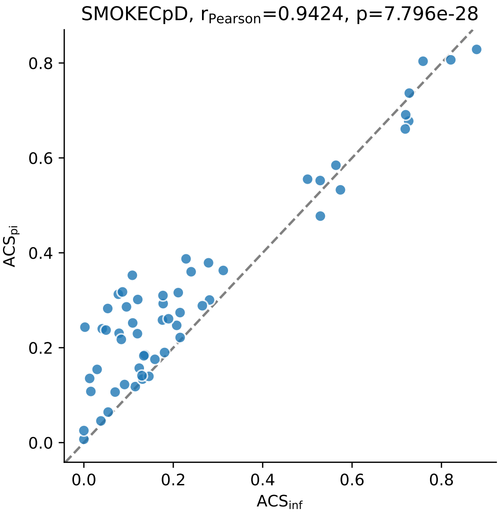


**Figure S13:** ACS comparison for SMOKECpD

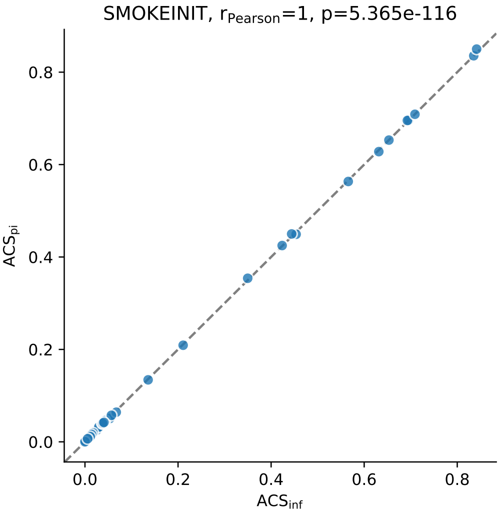


**Figure S13:** ACS comparison for SMOKEINIT

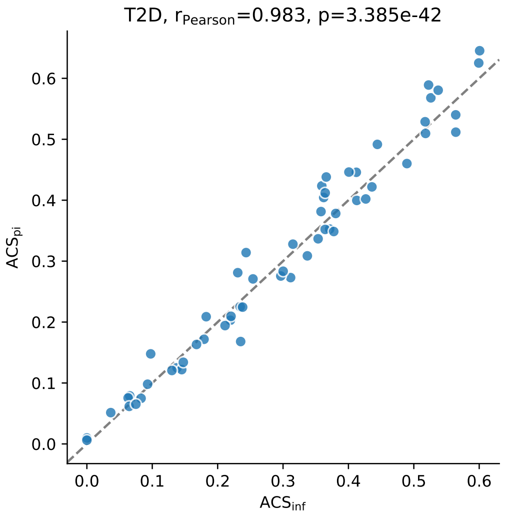


**Figure S13:** ACS comparison for T2D

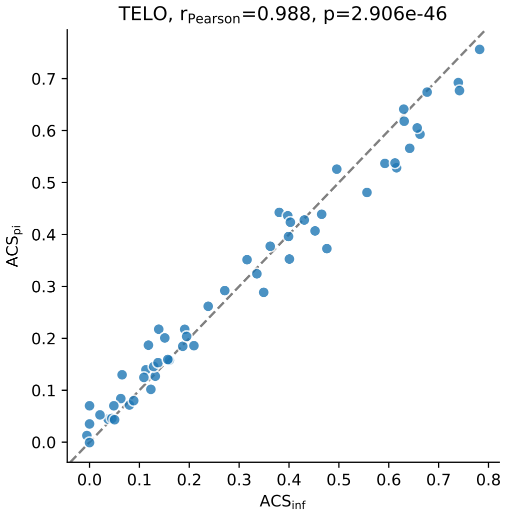


**Figure S13:** ACS comparison for TELO

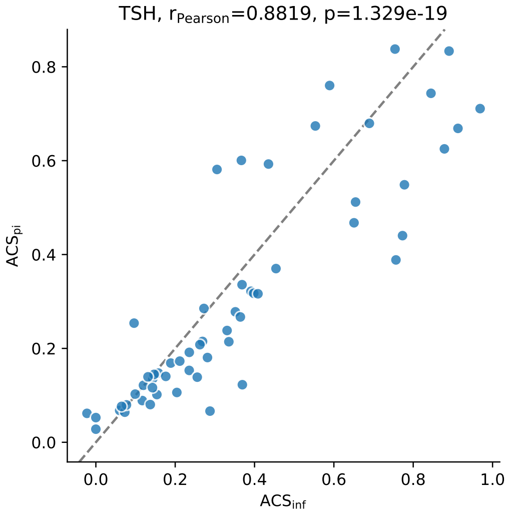


**Figure S13:** ACS comparison for TSH

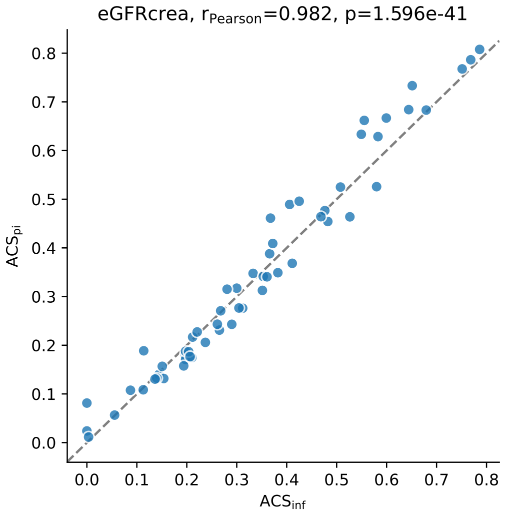


**Figure S13:** ACS comparison for eGFRcrea

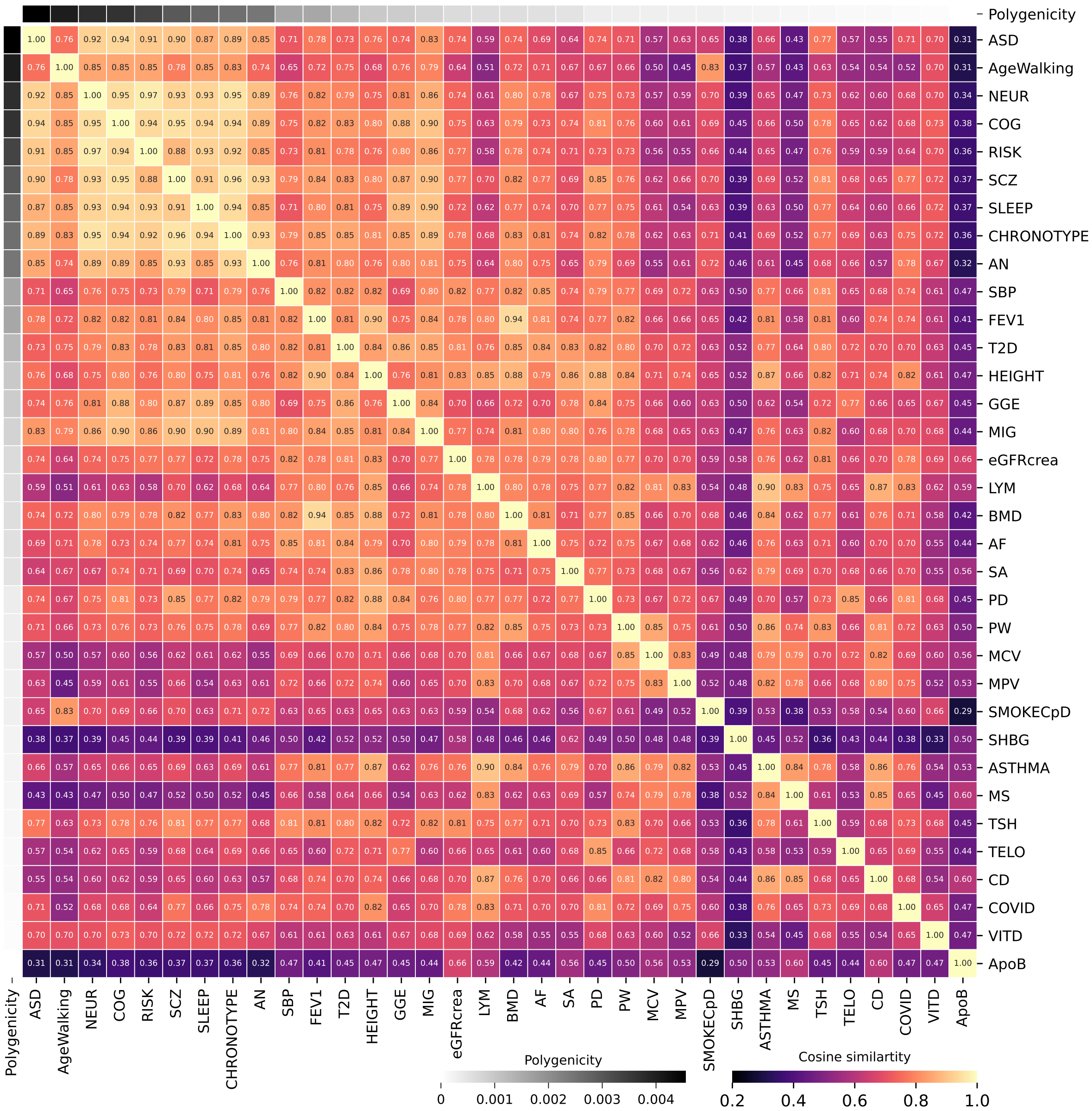


**(a)** Genome-wide.

**Figure S14:** Cross-trait SNP heritability cosine similarity heatmaps with traits ordered by decreasing polygenicity.

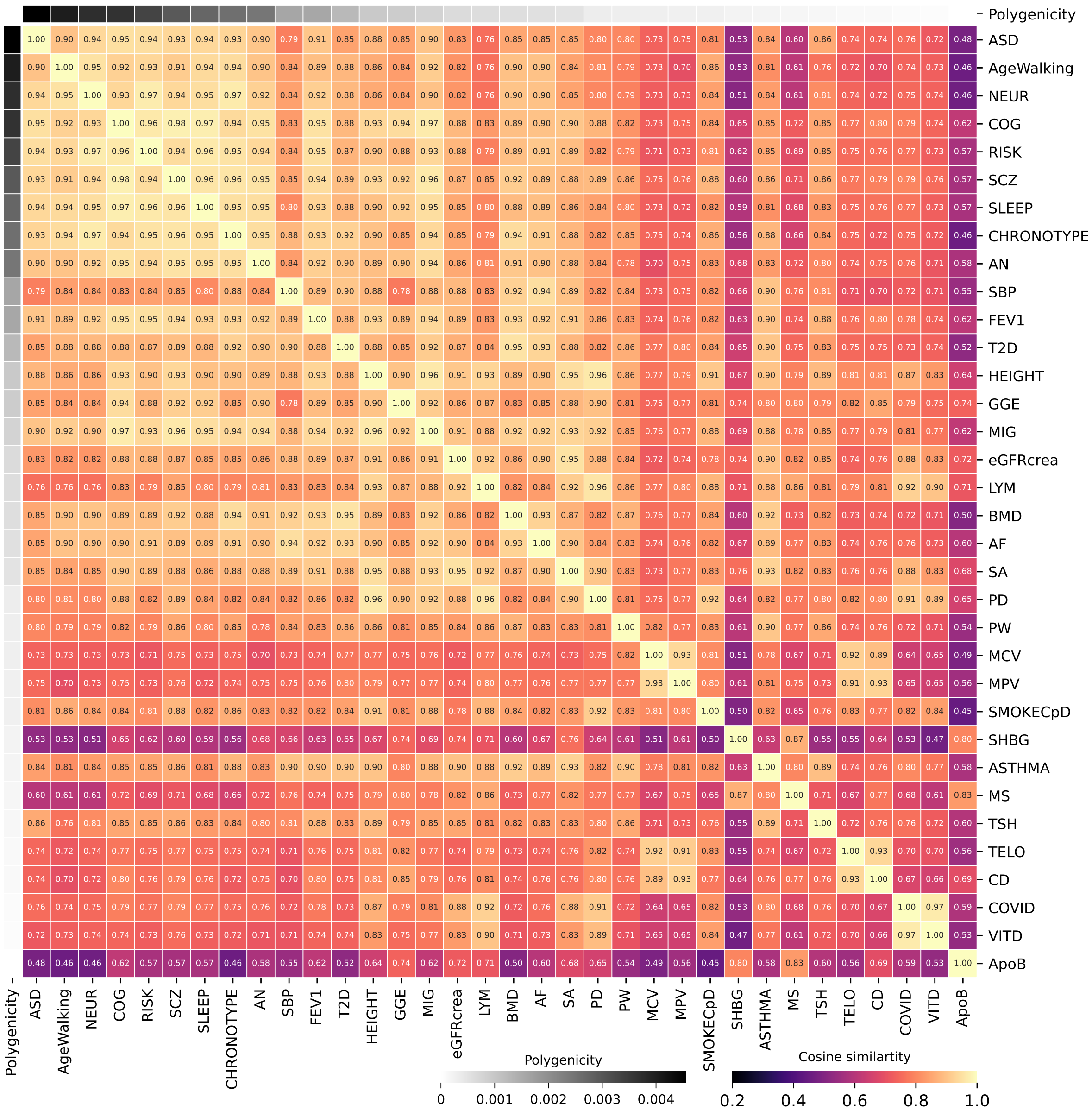

**(b)** Exon.

**Figure S14:** Cross-trait SNP heritability cosine similarity heatmaps with traits ordered by decreasing polygenicity.

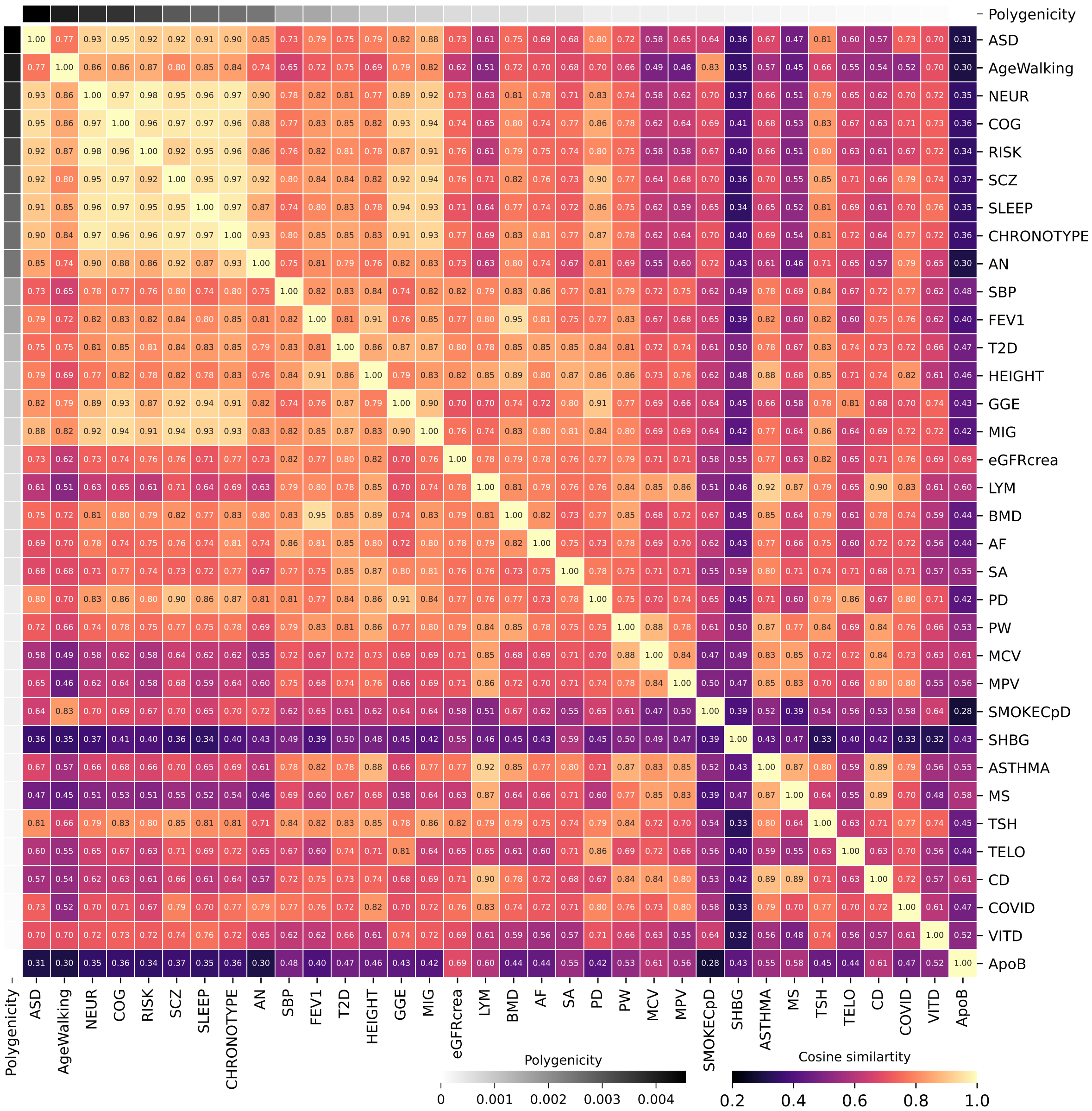


**(c)** Intron.

**Figure S14:** Cross-trait SNP heritability cosine similarity heatmaps with traits ordered by decreasing polygenicity.

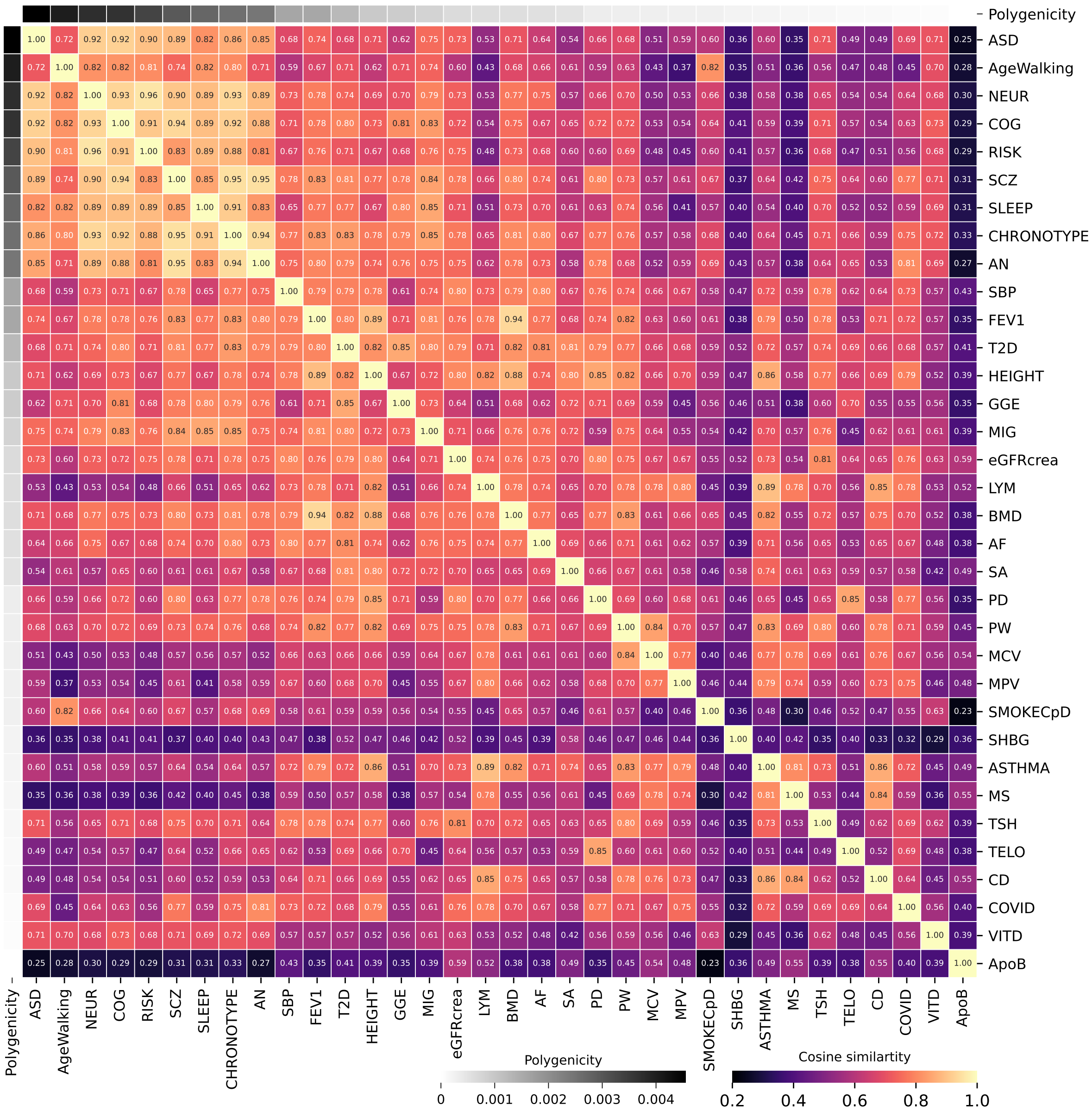

**(d)** Intergenic.

**Figure S14:** Cross-trait SNP heritability cosine similarity heatmaps with traits ordered by decreasing polygenicity.

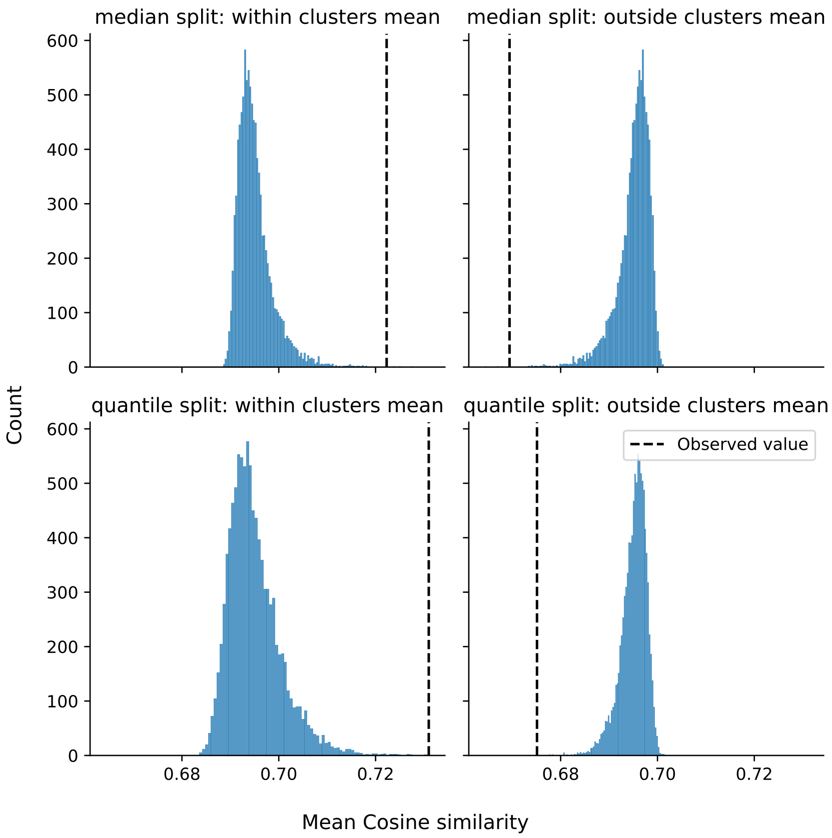


**Figure S15:** Permutation test of cross-trait per-SNP heritability similarity. Traits were binned by -$\log_{10}(\pi)$ (median: 17/17, tertiles: 12/11/11). Histograms show the null distribution of "within-between" differences of cosine similarity matrix from 1e4 label shuffles. Vertical dashed lines indicate the observed within- and between-bin means in the respective panels.

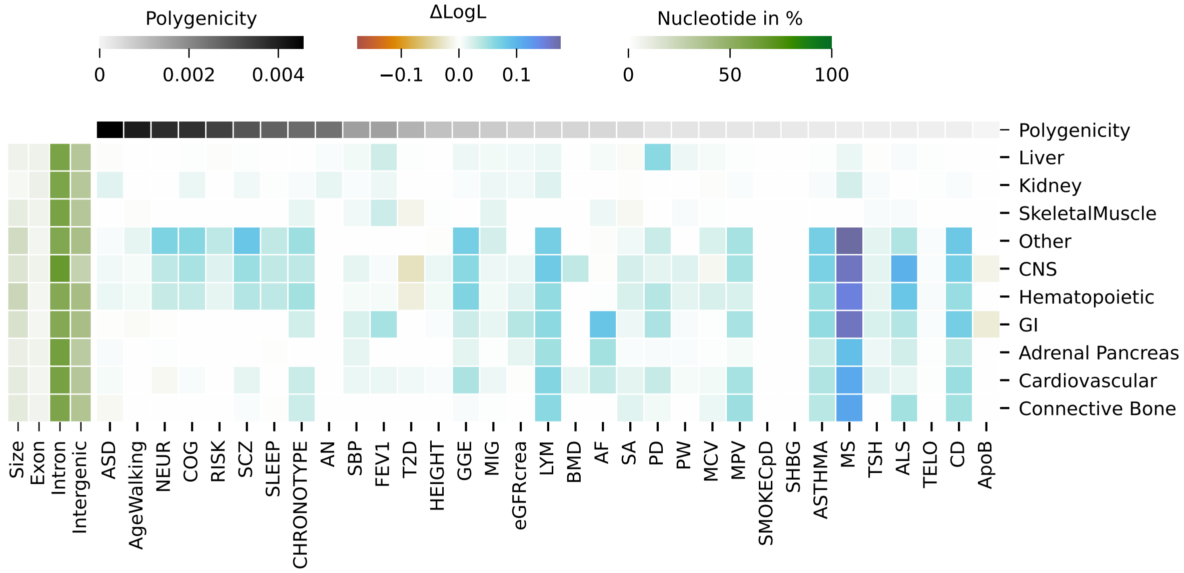


**Figure S16:** Contribution of 221 individual cell types from Finucane *et al.*[84] added jointly to the full model with 74 annotations. Phenotypes are ordered by their polygenicity remaining after pruning with $r_g$=0.3 favoring phenotypes with higher heritability. , There is an increased CNS ACS for SCZ and COG, elevated hematopoietic and gastrointestinal signals for MS and CD, and comparatively modest signals in liver or pancreas for T2D or lipids.
